# A series of spontaneously blinking dyes for super-resolution microscopy

**DOI:** 10.1101/2024.02.23.581625

**Authors:** Katie L. Holland, Sarah E. Plutkis, Timothy A. Daugird, Abhishek Sau, Jonathan B. Grimm, Brian P. English, Qinsi Zheng, Sandeep Dave, Fariha Rahman, Liangqi Xie, Peng Dong, Ariana N. Tkachuk, Timothy A. Brown, Robert H. Singer, Zhe Liu, Catherine G. Galbraith, Siegfried M. Musser, Wesley R. Legant, Luke D. Lavis

## Abstract

Spontaneously blinking fluorophores permit the detection and localization of individual molecules without reducing buffers or caging groups, thus simplifying single-molecule localization microscopy (SMLM). The intrinsic blinking properties of such dyes are dictated by molecular structure and modulated by environment, which can limit utility. We report a series of tuned spontaneously blinking dyes with duty cycles that span two orders of magnitude, allowing facile SMLM in cells and dense biomolecular structures.

## MAIN TEXT

Single-molecule localization microscopy (SMLM) circumvents the diffraction limit in fluorescence microscopy through the stochastic activation and localization of individual molecules, which enables the generation of pointillistic microcopy images^1–3^. The development of spontaneously blinking fluorophores^4–6^, such as HM-SiR^7^ (**1**; **Fig. 1a**) simplifies SMLM by eliminating the need for photoactivatable fluorophores^8^ or redox buffers^9^. The on/off ratio (*i.e.*, duty cycle) of such dyes is a product of both the molecular structure and the chemical environment; the lack of duty cycle control can limit utility depending on labeling strategy and sample type^5, 10^. Here, we synthesized and evaluated a series of hydroxymethyl-Si-rhodamine (HM-SiR) analogs based on the bright Janelia Fluor (JF) dyes^11–14^, incorporating substituted pyrrolidines or azetidines to tune the duty cycle across two orders of magnitude. We demonstrate the utility of these dyes in cells using the HaloTag system and fluorescence *in situ* hybridization (FISH), and *in vitro* with biomolecular condensates (BMCs) using direct protein labeling. This panel of new dyes will enable straightforward SMLM and other super-resolution microscopy experiments in a variety of biological samples.

**Figure 1.**
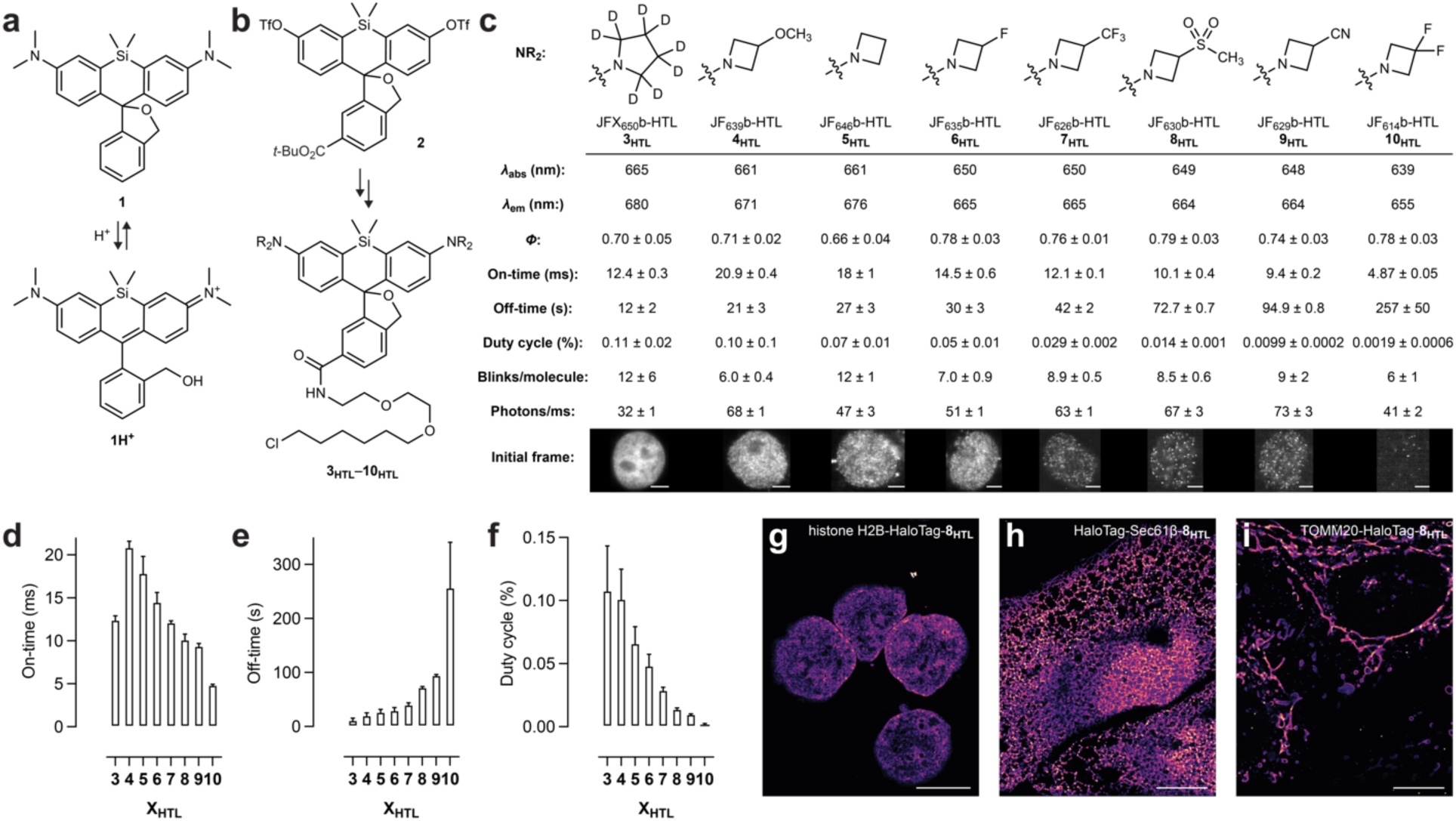
Synthesis and properties of tuned spontaneously blinking fluorophores. (**a**) The structure and dynamic protonation/deprotonation of HM-SiR (**1**). (**b**) Synthesis of HaloTag ligands **3_HTL_**–**10_HTL_** from ditriflate **2**. (**c**) Properties of HaloTag ligands **3_HTL_**–**10_HTL_** measured *in vitro* in 10 mM citrate buffer, pH 3.0 containing 150 mM NaCl and 0.1% (w/v) SDS (*λ*_abs_, *λ*_em_, *Φ*) or in paraformaldehyde-fixed COS-7 cells expressing HaloTag–histone H2B fusion proteins (other characteristics); error for *Φ* indicates ±SD (*n* = 10) and errors for other values indicate ±SEM (*n* = 3); images are the initial frame (25 ms exposure time) from an SMLM imaging session using cells expressing histone H2B-HaloTag labeled to saturation with the respective ligands; scale bars: 5 μm. (**d–f**) Mean on-times (**d**), off-times (**e**), and duty cycles (**f**) measured for HaloTag ligands **3_HTL_**–**10_HTL_**; *n* = 3; error bars indicate ±SEM. (**g**–**i**) SMLM images of paraformaldehyde-fixed cells expressing HaloTag fused to histone H2B (**g**; COS-7 cells), Sec61β (**h**; U2OS cells), or TOMM20 (**i**; U2OS cells) and labeled with JF_630_b-HTL (**8_HTL_**); localization precision (σ; mean±SD) values: 13.0 ± 5.8 nm (**g**), 10.7 ± 2.9 nm (**h**), 13.0 ± 5.5 nm (**i**); scale bars: 10 μm; these imaging experiments were repeated with similar results.

The spontaneously blinking properties of HM-SiR depend on stochastic protonation/deprotonation that switches the dye between a neutral colorless “off” state (**1**) and a cationic, fluorescent “on” state (**1H^+^**; **Fig. 1a**).^7^ Unfortunately, the duty cycle of **1** is too high for use in many biological samples. This can limit its use in SMLM to cellular structures with relatively low labeling densities like the nuclear pore^10^, or in membranes where the duty cycle is decreased by the lipophilic environment^5^. Although improved versions of **1** and the development of other spontaneously blinking fluorophores have broadened the utility of this label type^4–6^, microscopists lack a panel of systematically tuned dyes to address the diverse sample types, labeling densities, dye attachment strategies, and hardware used in modern biological imaging. To address this deficiency, we developed a new, efficient synthesis of reduced Si-fluorescein bistriflate **2** (**Supplementary Note 1**), which was derivatized to yield a panel of eight spontaneously blinking HM-SiR analogs as HaloTag ligands (HTLs; **3_HTL_**–**10_HTL_**; **Fig. 1b,c**, **Extended Data** Fig. 1**, Supplementary Note 2**). These dyes were named after their parent fluorophores^11–14^—JFX_650_, JF_646_, JF_639_, JF_635_, JF_630_, JF_629_, JF_626_, and JF_614_—with a “b” appended to the name to indicate their blinking character. For example, the hydroxymethyl derivative of JF_635_-HaloTag ligand was termed “JF_635_b-HTL” (**6_HTL_**; **Fig. 1c**).

The properties of these dye ligands were evaluated *in vitro* and in fixed cells expressing HaloTag–histone H2B fusion proteins (**Fig. 1c**, **Movie 1**). As expected^12^, stronger electron-withdrawing substituents on the auxochrome groups gave trends to lower absorption maxima (*λ*_abs_) and emission maxima (*λ*_em_) along with higher fluorescence quantum yield (*Φ*) values (**Fig. 1c**, **Extended Data** Fig. 2). We discovered that electronegative substituents modestly decreased the on-times (**Fig. 1c,d**, **Extended Data** Fig. 3a,b) and substantially increased the off-times (**Fig. 1c,e**, **Extended Data** Fig. 3c,d) yielding dyes with lower duty cycles (**Fig. 1c,f**). The number of times a molecule blinked before irreversibly photobleaching and the average photon emission rate for each blink were largely consistent across the series (**Fig. 1c**, **Extended Data** Fig. 3e–g). The methylsulfone-containing JF_630_b-HTL (**8_HTL_**) showed suitable performance in SMLM imaging of paraformaldehyde-fixed cells expressing HaloTag fusions of histone H2B (nucleus; **Fig. 1g**), which is consistent with the relatively low duty cycle = 0.014% (**Fig. 1c**). Compound **8_HTL_** could also be used to image HaloTag fusions of Sec61β (endoplasmic reticulum; **Fig. 1h**), and TOMM20 (mitochondria; **Fig. 1i**). These SMLM experiments were performed in standard phosphate-buffered saline (PBS) without any additives and provided high localization precisions (*σ* ≤ 13.0 nm; **Extended Data** Fig. 3h,i). Additionally, a JF_630_b-Hoechst (**8_HST_**) conjugate was used for SMLM (**Extended Data** Fig. 4) and the HaloTag ligands of JF_639_b (**4_HTL_**), JF_635_b (**6_HTL_**), and JF_630_b (**8_HTL_**) were deployed for super-resolution optical fluctuation imaging^15^ (SOFI), a complementary technique to SMLM that can utilize blinking fluorophores with higher duty cycles (**Extended Data** Fig. 5)^16^.

We then tested the utility of these spontaneously blinking dyes to characterize the physical properties of BMCs, which serve major roles in cellular organization, regulation, and metabolic control^17^. Cellular BMCs are dense (∼10^6^ molecules/µm^3^)^18^ and complex^17^ macromolecular assemblies; measurement of the structural and dynamical features of BMCs remains an important but challenging problem. We envisioned adding a small amount of protein labeled with a spontaneously blinking dye to probe the BMC’s structure using both SMLM and single-molecule rotational diffusion (SiMRoD) microscopy^19^; we choose the intermediate duty cycle JF_635_b (**Fig. 1c**) due to the relatively low labeling density in this experiment. We synthesized JF_635_b-maleimide (**6_MAL_**; **Fig. 2a**, **Supplementary Note 2**) and labeled the A2C variant of the fused in sarcoma (FUS) protein, yielding FUS^A2C^-JF_635_b. Condensates were then formed from wildtype FUS doped with a 0.3% mole fraction of FUS^A2C^-JF_635_b. A 3D map of all localizations obtained by astigmatic single-molecule imaging^10^ revealed a condensate with a flattened-dome shape containing regions of different densities after 12 h of aging (**Fig. 2b**, **Extended Data** Fig. 6a). Assuming a FUS concentration of ∼160 mg/mL inside the condensates,^20^ FUS^A2C^-JF_635_b was present at ∼5000 molecules/µm^3^. Even with this high number of molecules per volume, single JF_635_b fluorophores were readily imaged revealing regions of different localization densities, in some cases >10^5^ events/µm^3^ (**Fig. 2b**, **Extended Data** Fig. 6b). These variable densities within the BMC suggest local compaction or selective aggregation of the protein within the condensate. The JF_635_b label proved superior to HM-SiR for this BMC imaging experiment. Condensates doped with FUS^A2C^-JF_635_b showed a constant number of activated molecules over time, but use of HM-SiR-maleimide (**1_MAL_**; **Fig. 2a**) resulted in condensates with a higher initial fraction of molecules in the on-state along with faster photobleaching and reversible photoinactivation (**Extended Data** Fig. 6c, **Movie 2**).

**Figure 2.**
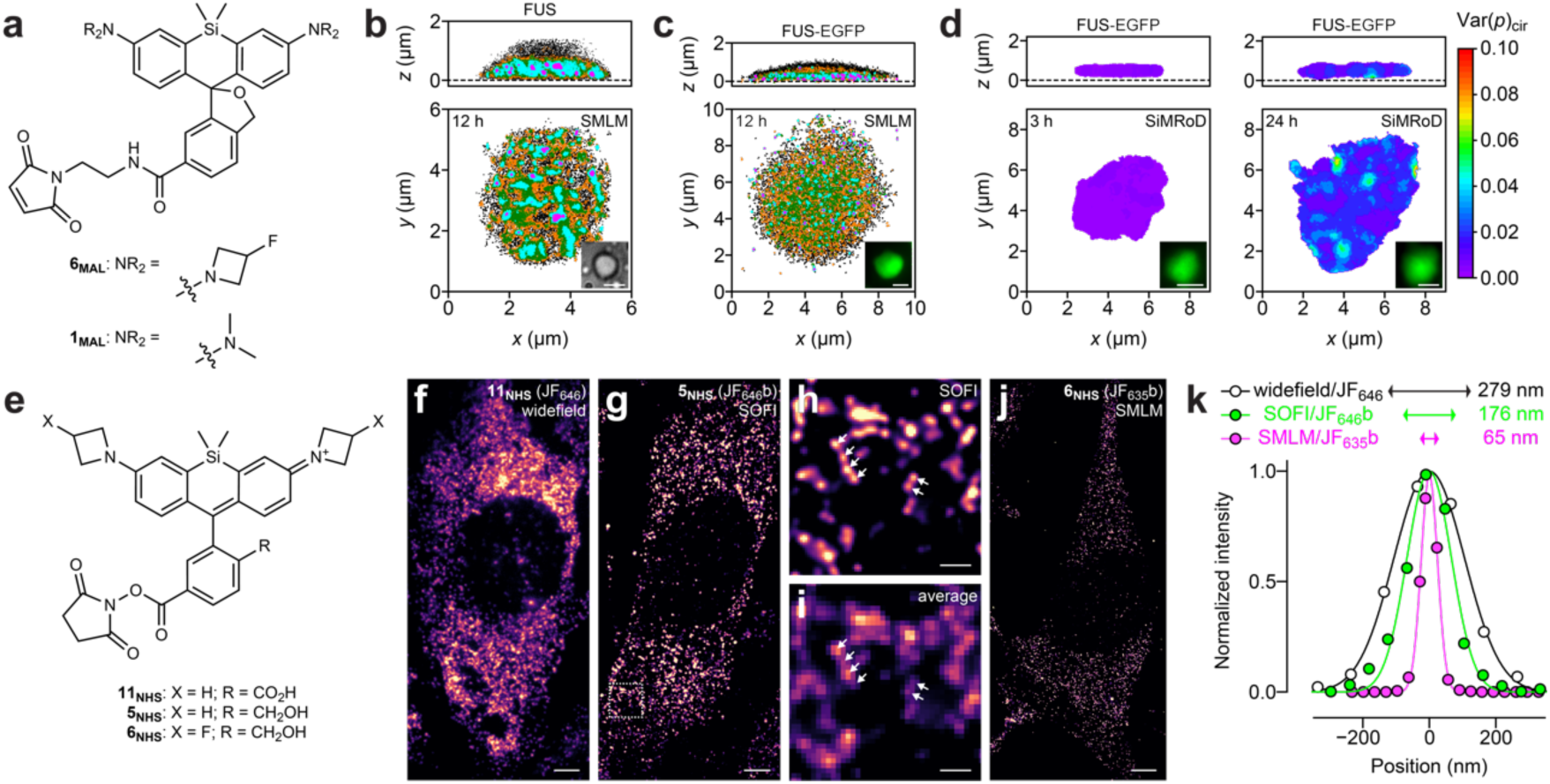
Applications of tuned spontaneously blinking fluorophores. (**a**) Structures of JF_635_b-maleimide (**6_MAL_**) and HM-SiR-maleimide (**1_MAL_**). (**b**) 3D SMLM map of a FUS coacervate aged for 12 h containing 20 nM FUS^A2C^-JF_635_b; inset shows brightfield image; scale bar = 4 μm; *n* = 31,123 total localizations; colors indicate localization density within a 100 nm radius sphere: black = 1; orange 2–10; green = 11–20; cyan = 21–40; magenta = 41–100. (**c**) 3D SMLM map of a FUS-EGFP coacervate aged for 12 h containing 20 nM FUS^A2C^-JF_635_b; inset shows wide-field fluorescence image; scale bar = 4 μm; *n* = 26,895 total localizations; colors indicate localization density within a 100 nm radius sphere: black = 1; orange 2–4; green = 5–8; cyan = 9–20; magenta = 21–40; the color ranges were adjusted to account for the lower total density of localizations compared to the condensate in **b**. (**d**) 3D SiMRoD map of FUS-EGFP coacervates aged for 3 h (left) or 24 h (right) containing 2 nM FUS^A2C^-JF_635_b; insets show wide-field fluorescence images, scale bars = 4 μm; color scale indicates the variance of single-molecule polarization measurements using circularly polarized light illumination (Var(*p*)_cir_) and calculated within a 400 nm radius sphere; 5 ms/polarization measurement. (**e**) Structures of JF_646_-NHS (**11_NHS_**), JF_646_b-NHS (**5_NHS_**), and JF_635_b-NHS (**6_NHS_**). (**f**) Widefield RNA-FISH image of MEF cells expressing MS2 in the 3′ UTR of the beta-actin gene and labeled with JF_646_-oligonucleotide from **11_NHS_**; the image is from a single frame; scale bar: 5 μm. (**g**) SOFI RNA-FISH image of MEF cells expressing MS2 in the 3′ UTR of the beta-actin gene and labeled with JF_646_b-oligonucleotide from **5_NHS_**; scale bar: 5 μm. (**h,i**) Zoomed area from **g** rendered as a SOFI image (**h**) or averaged image (**i**); scale bars: 1 μm. (**j**) SMLM RNA-FISH image of MEF cells expressing MS2 in the 3′UTR of the beta-actin gene and labeled with JF_635_b-oligonucleotide from **6_NHS_**; scale bar: 5 μm. (**k**) Intensity profiles, Gaussian fits, and FWHM measurements of RNA foci from RNA-FISH experiments using JF_646_/confocal (*n* = 3), JF_646_b/SOFI (*n* = 4), or JF_635_b/SMLM (*n* = 3). Images in **f**–**j** were taken using HILO illumination with 30 ms exposure time (33.33 Hz). All imaging experiments were repeated with similar results.

The FUS^A2C^-JF_635_b conjugate was then used to image the BMCs formed from the FUS–EGFP fusion protein. A substantially flatter shape and lower fluorophore densities were observed (**Fig. 2c**, **Extended Data** Fig. 6d), which suggests a more liquid-like structure. JF_635_b also enabled SiMRoD microscopy, which has distinct advantages over bulk polarization measurements. The variance of single-molecule polarization measurements when using circularly polarized light, Var(*p*)_cir_, can report biochemically relevant differences in the rotational diffusion coefficient (*D*_r_) across six orders of magnitude^19^. SiMRoD experiments revealed that regions of lower rotational mobility formed in the BMCs after 24 h (**Fig. 2d**, **Extended Data** Fig. 6e,f), reminiscent of the puncta observed in the particle density maps (**Fig. 2c**). Under these measurement conditions a positive Var(*p*)_cir_ value indicates a *D*_r_ < 10^4^ rad^2^/s, which is >10^3^ slower than a freely tumbling molecule. This result shows that the proteins within aged BMCs can have substantially limited rotational mobility and demonstrates the utility of JF_635_b to interrogate the internal structure of BMCs using single-molecule measurements.

These dyes were then evaluated as labels for DNA using amine-terminated oligonucleotides. The standard, nonblinking label JF_646_-*N*-hydroxysuccinimidyl ester^11^ (JF_646_-NHS; **11_NHS_**), JF_646_b-NHS (**5_NHS_**), and JF_635_b-NHS (**6_NHS_**; **Fig. 2e**, **Supplementary Note 2**) were used to label oligonucleotides with sequences that were complementary to the RNA sequence that binds the bacteriophage MS2 coat protein. These probes were used in fluorescence *in situ* hybridization (FISH) experiments in cells derived from a transgenic mouse containing an MS2 binding site in the 3′ untranslated region (UTR) of the beta-actin gene^21^. Widefield imaging experiments using JF_646_ gave diffraction-limited resolution of RNA foci (**Fig. 2f,k**). Both labels could be visualized by SOFI (**Fig. 2g–i,k**, **Extended Data** Fig. 7a) and JF_635_b enabled the highest resolution *via* SMLM due to its lower duty cycle (**Fig. 2j,k**, **Extended Data** Fig. 7b–e). JF_635_b-NHS (**6_NHS_**) was also used to label oligonucleotides for super-resolution assays of transposase-accessible chromatin (ATAC– SMLM)^22^ and showed five-fold higher localizations/cell compared to probes labeled with photoactivatable JF_646_-*N*-hydroxysuccinimidyl ester (PA-JF_646_-NHS, **12_NHS_**; **Extended Data** Fig. 8). Although counting the absolute number of transposase-accessible sites with JF_635_b is complicated by the multiple blinks/molecule (**Fig. 1c**), imaging the spatial distribution of accessible sites within the nucleus is faster and operationally simpler due to the spontaneously blinking dye’s short on-times, tuned duty cycle, and lack of requisite photoactivation.

Modern super-resolution imaging techniques utilize a broad range of biological samples, labeling strategies, imaging conditions, and hardware that must be balanced to extract the most information from an experiment. We synthesized a panel of spontaneously blinking fluorophores with different duty cycles to accommodate this experimental variety (**Fig. 1c–f**). For SMLM imaging of densely labeled samples, we advise starting with low duty cycle dyes such as JF_630_b, which proved useful for imaging highly expressed HaloTag fusions (**Fig. 1g–i**). For localization microscopy experiments with lower labeling density, we suggest the intermediate duty cycle JF_635_b, which facilitated SMLM and single-molecule polarization measurements in BMCs (**Fig. 2b–d**) along with SMLM of RNA (**Fig. 2j**) and accessible chromatin (**Extended Data** Fig. 8). For SOFI, which offers higher temporal resolution but lower spatial resolution^16^, we recommend dyes with higher duty cycles such as JF_646_b (**Fig. 2g–i**) and JF_639_b (**Extended Data** Fig. 5), which enables calculation of higher resolution images using fewer frames. Overall, this series of bright, spontaneously blinking fluorophores will allow experiment-specific optimization of the duty cycle, thereby facilitating straightforward imaging below the diffraction limit.

## Acknowledgements

1. S. Banala, P. Kumar, and A. Osowski (Janelia); S. Hutton and D. Dormann (Johannes Gutenberg University) for contributive discussions. Plasmid pMal-*Tev*-FUS(WT)-EGFP-*Tev*-His_6_ was a gift of D. Dormann. This research was supported the Howard Hughes Medical Institute (HHMI), the Janelia Visiting Scientist Program, grants from the NIH National Institutes of General Medical Sciences (NS083085 to R.H.S; GM117188 to C.G.G.; GM126190 to S.M.M.; 1DP2GM136653 to W.R.L), the Welch Foundation (579662 to S.M.M.), and the W.M. Keck Foundation (to C.G.G.). W.R.L. acknowledges additional support from the Searle Scholars program, the Beckman Young Investigator Program, and the Packard Fellowship for Science and Engineering. This article is subject to HHMI’s Open Access to Publications policy. HHMI lab heads have previously granted a nonexclusive CC BY 4.0 license to the public and a sublicensable license to HHMI in their research articles. Pursuant to those licenses, the author-accepted manuscript of this article can be made freely available under a CC BY 4.0 license immediately upon publication.

## Declarations of Interests

US Patents 9,933,417 and 11,091,643 and US Patent Applications 2021/0171490 and 2021/0085805 describing azetidine-containing fluorophores and variant compositions (with inventors J.B.G., Q.Z., and L.D.L.) are assigned to HHMI. L.D.L. is a founder and shareholder of Eikon Therapeutics. The authors declare no other competing interests.

**Extended Data Figure 1.**
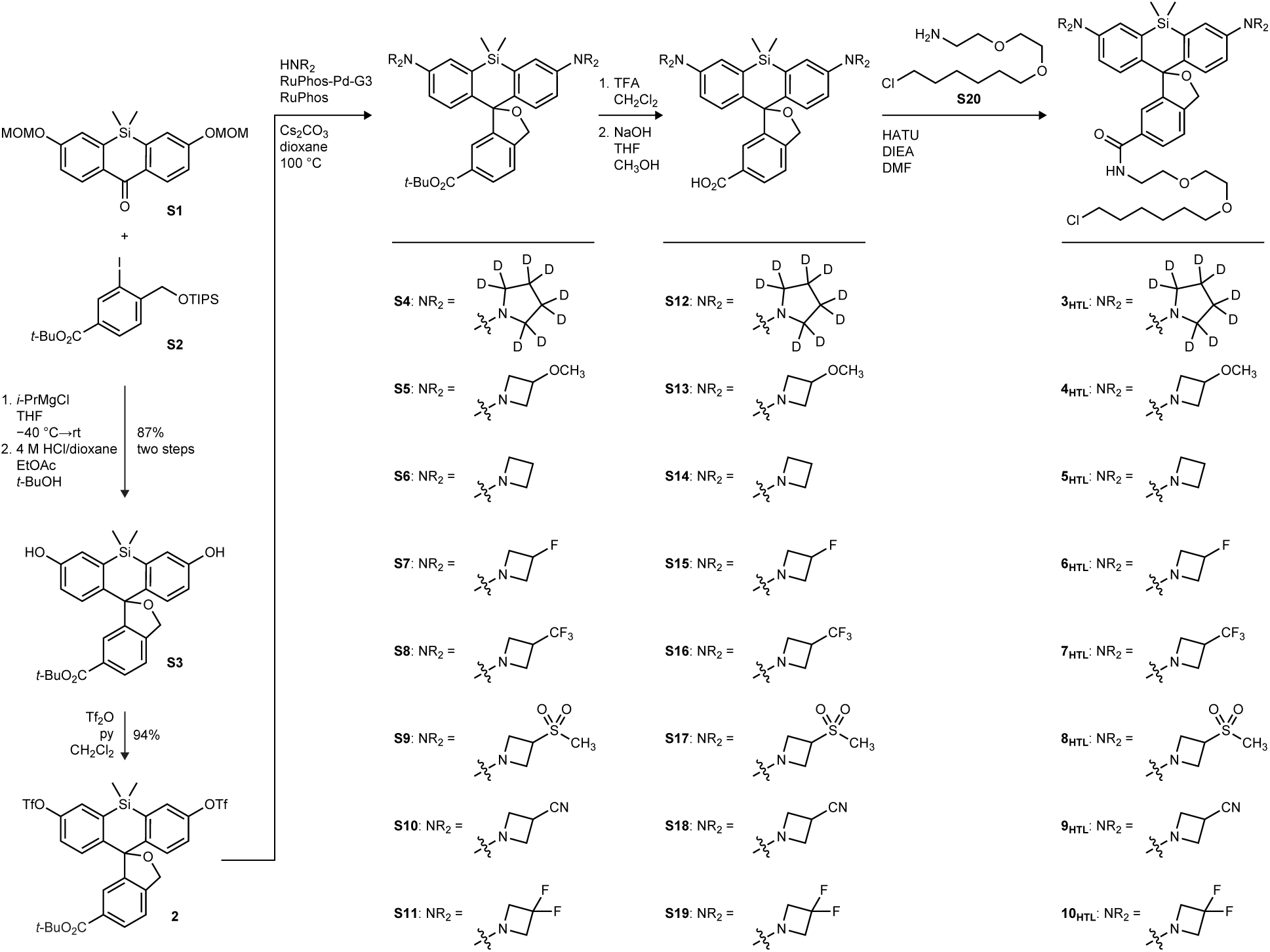
Divergent synthesis of 3_HTL_–10_HTL_.

**Extended Data Figure 2.**
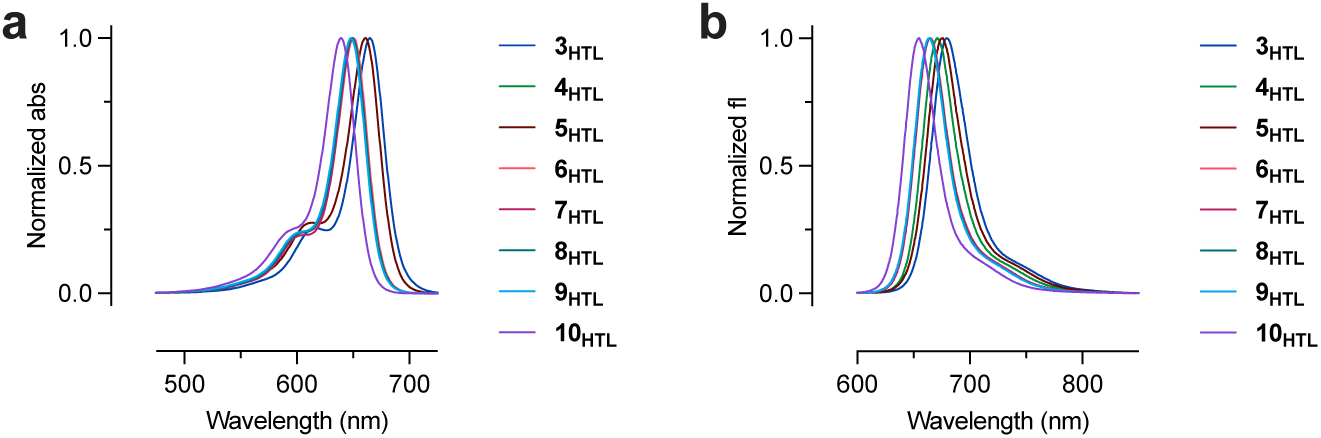
Spectral properties of 3_HTL_–10_HTL_. (**a,b**) Normalized absorbance (**a**) and fluorescence emission (**b**) of ligands **3_HTL_**–**10_HTL_** in 10 mM citrate buffer, pH 3.0 containing 150 mM NaCl and 0.1% (w/v) SDS.

**Extended Data Figure 3.**
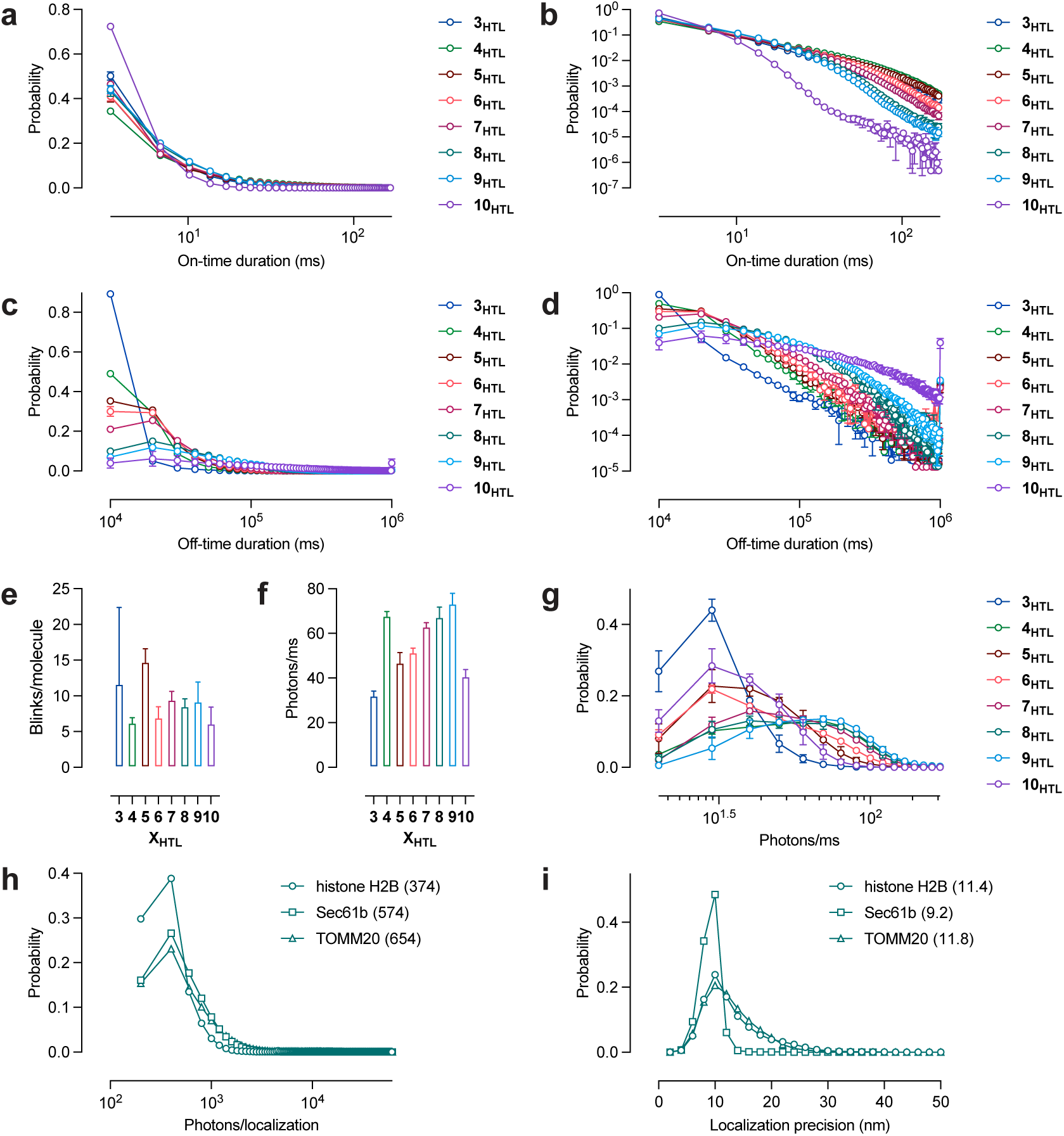
Characterization of 3_HTL_–10_HTL_. (**a,b**) Probability *vs.* on-time (ms) of ligands **3_HTL_**–**10_HTL_** in formaldehyde-fixed cells expressing histone H2B-HaloTag fusion proteins as semi-log (**a**) and log–log (**b**) plots. (**c,d**) Probability *vs.* off-time (ms) of ligands **3_HTL_**–**10_HTL_** in formaldehyde-fixed cells expressing histone H2B-HaloTag fusion proteins as semi-log (**c**) and log–log (**d**) plots. (**e,f**) Mean number of blinks per molecule (**e**) and photon emission rate (**f**) for the HaloTag conjugates of **3_HTL_**–**10_HTL_** in cells; *n* = 3; error bars indicate ±SEM. (**g**) Probability *vs.* photons/ms of ligands **3_HTL_**–**10_HTL_** in formaldehyde-fixed cells expressing histone H2B-HaloTag fusion proteins as a semi-log plot. (**h**) Probability *vs.* photons/localization of **8_HTL_** in formaldehyde-fixed cells expressing histone H2B-HaloTag, HaloTag-Sec61b, or TOMM20-HaloTag fusion proteins as a semi-log plot; numbers in parentheses indicate median values in photons. (**i**) Probability *vs.* localization precision of **8_HTL_** in formaldehyde-fixed cells expressing histone H2B-HaloTag, HaloTag-Sec61b, or TOMM20-HaloTag fusion proteins; numbers in parentheses indicate median values in nm.

**Extended Data Figure 4.**
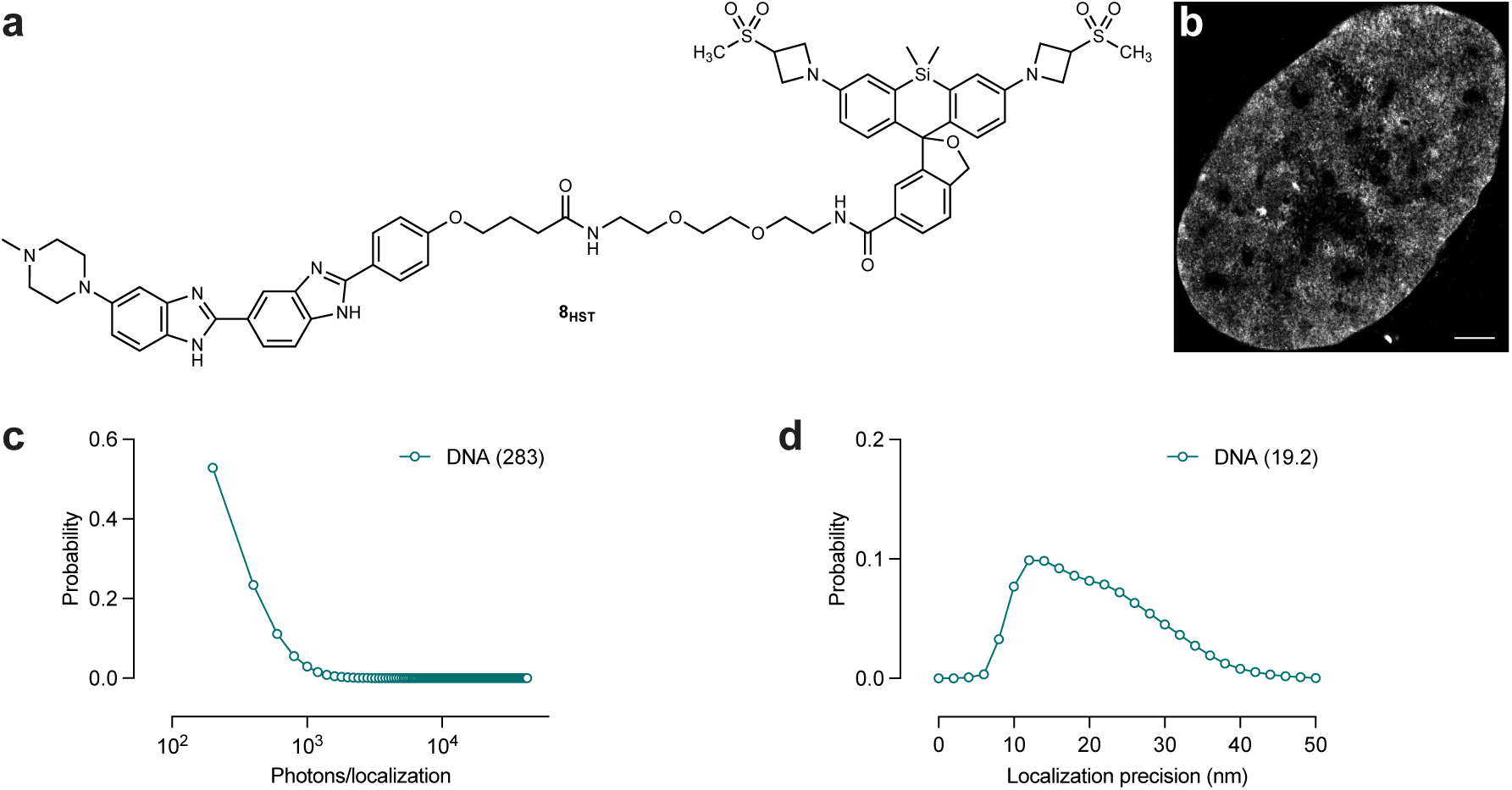
Characterization of 8_HST_. (**a**) Chemical structure of JF_630_-Hoechst (**8_HST_**). (**b**) SMLM images of paraformaldehyde-fixed hTERT RPE-1 cells labeled with **8_HST_**; scale bar: 2 μm. (**c**) Probability *vs.* photons/localization of **8_HST_** in formaldehyde-fixed cells as a semi-log plot; number in parentheses indicates median values in photons. (**d**) Probability *vs.* localization precision of **8_HST_** in formaldehyde-fixed cells; number in parentheses indicates median values in nm.

**Extended Data Figure 5.**
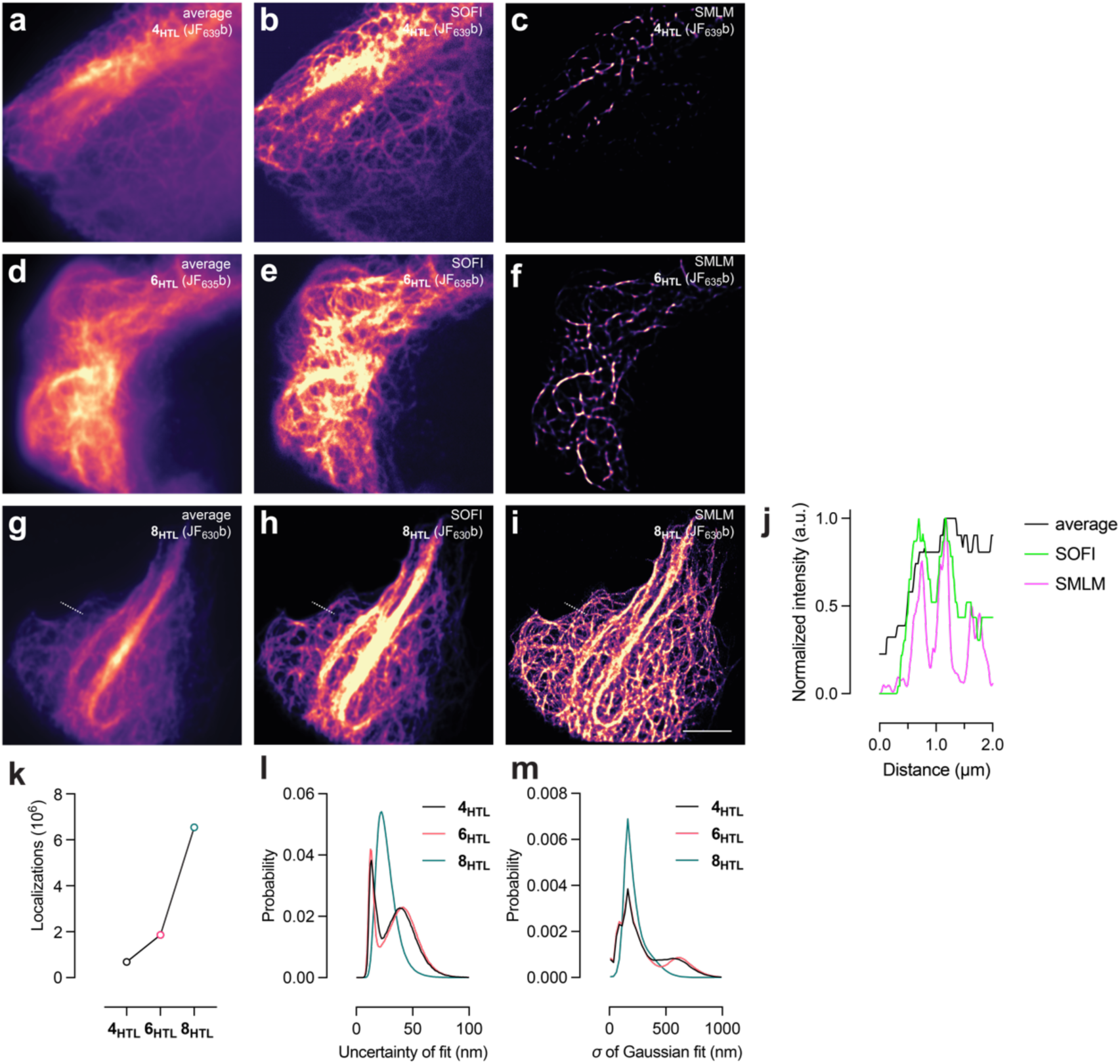
Super-resolution optical fluctuation imaging (SOFI) and single-molecule localization microscopy (SMLM) using HaloTag ligands 4_HTL_, 6_HTL_, and 8_HTL_. (**a–i**) Images of live U2OS cells expressing ensconsin-HaloTag fusion proteins and labeled with **4_HTL_** (**a–c**), **6_HTL_**, (**d–f**), and **8_HTL_** (**g–i**); panels **a**, **d**, and **g** are average images; **b**, **e**, and **h** are SOFI images reconstructed from 5000 frames taken at 100 frames/s; panels **c**, **f**, and **i** are SMLM images reconstructed from 100,000 frames; scale bar = 5 μm. (**j**) Line-scan of region shown in panels **g–i** comparing resolution of average, SOFI, and SMLM images. (**k– m**) Statistics for SMLM images shown in panels **c**, **f**, and **i** showing total localizations (**k**), histogram of uncertainty of fit (**l**), and histogram of σ of Gaussian fit. The shoulder in the σ of Gaussian fit plot for **4_HTL_** and **6_HTL_** suggests attempted fits of multiple molecules as one event, which is consistent with the higher duty cycle of these dyes.

**Extended Data Figure 6.**
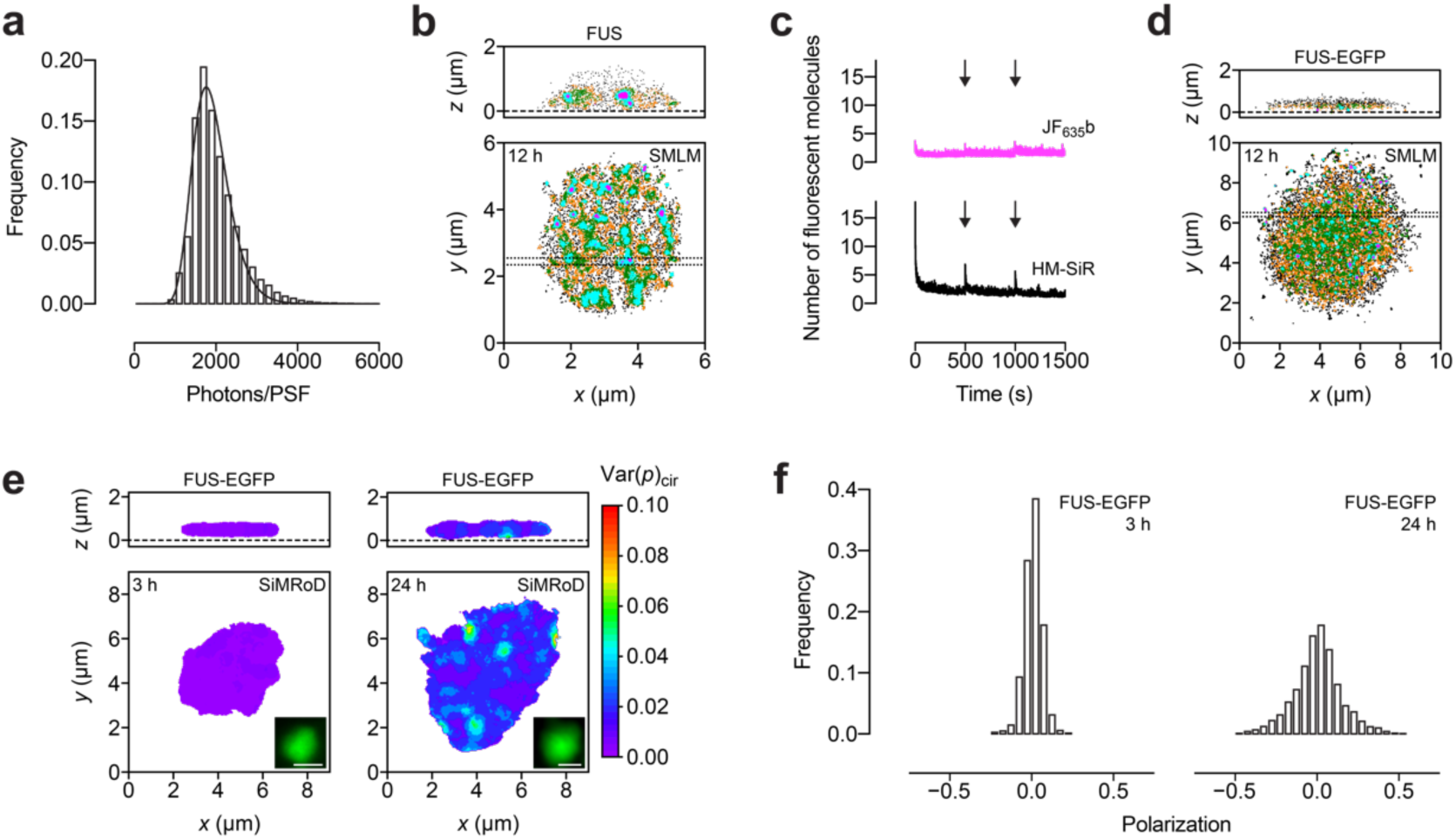
Characterization of FUS condensates. (**a**) Intensity frequency histogram for the localizations from FUS condensate doped with FUS^A2C^-JF_635_b (**Fig. 2b**; 30 ms/frame) fit with a log-normal distribution *y* = [*b*/((*xσ*(2*π*)^½^]*exp[−(ln(*x*)−*µ*)^2^/2*σ*^2^], where *b* = 200 (bin size), *µ* = 7.53, and *σ* = 0.245. Only localizations with ≥1000 photons were included in **Fig. 2b**, yielding average *x*, *y*, and *z* precisions of approximately 5.6, 9.0, and 15 nm, respectively. The difference in average *x* and *y* precisions arises from the bias in *z* values.^23^ (**b**) 200 nm sections of the 3D SMLM map from **Fig. 2b** showing a coacervate of FUS containing 20 nM FUS^A2C^-JF_635_b aged for 12 h. The bottom panel shows *z* = 100–300 nm. The top panel show the section cut through the region identified by the horizontal dashed lines in the bottom panel. (**c**) Fluorescence decay and recovery of FUS coacervates containing 5 nM FUS^A2C^-JF_635_b (magenta) or FUS^A2C^-HM-SiR (black) normalized to the number of fluorescent molecules per frame; the illumination laser was turned off for 500 s at *t* = 500 s and *t* = 1000 s (arrows) to allow depopulation of dark states. (**d**) 200 nm sections of the 3D SMLM map from **Fig. 2c** showing a coacervate of FUS-EGFP containing 20 nM FUS^A2C^-JF_635_b aged for 12 h. The bottom panels show *z* = 100–300 nm. The top panel shows the section cut through the region identified by the horizontal dashed lines in the bottom panels. (**e**) Reprint of **Fig. 2d** showing 3D SiMRoD maps of a FUS-EGFP coacervates aged for 3 h (left) and 24 h (right) containing 2 nM FUS^A2C^-JF_635_b; insets show wide-field fluorescence images, scale bar = 2 μm; color scale indicates the variance of single-molecule polarization measurements using circularly polarized light illumination (Var(*p*)_cir_) and calculated within a 400 nm radius sphere. (**f**) Polarization frequency histograms for the FUS-EGFP condensates in **e** aged for 3 h (left) and 24 h (right); the broader histogram after 24 h indicates reduced rotational mobility in aged condensates.

**Extended Data Figure 7.**
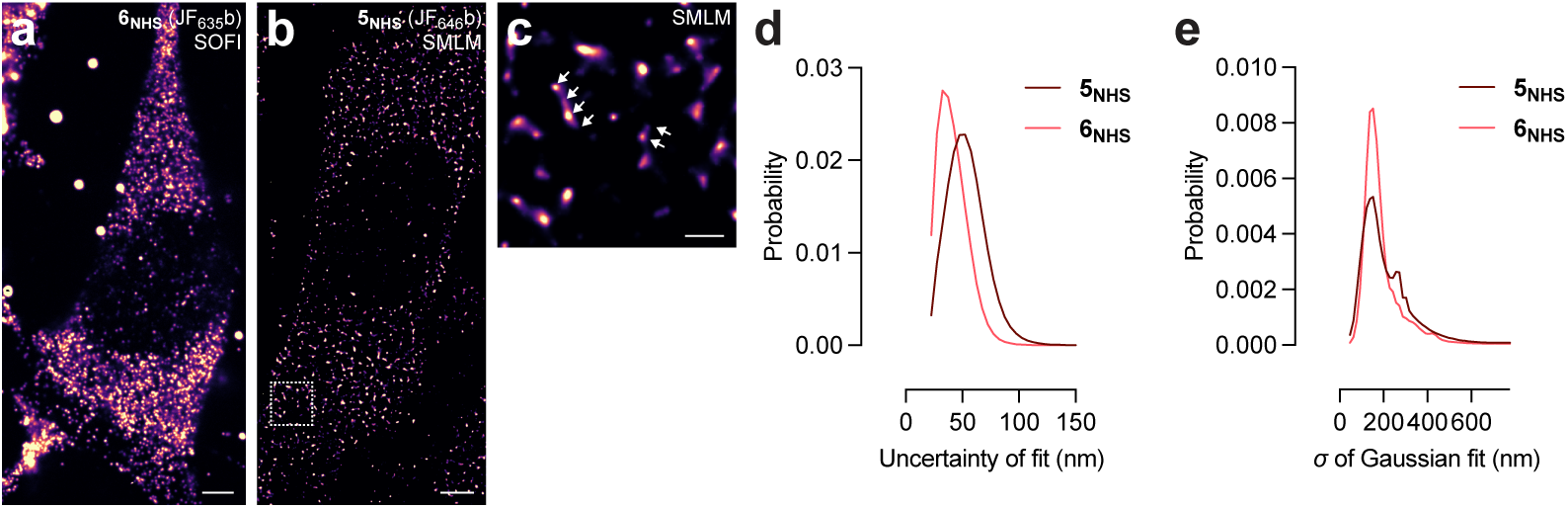
Comparison of PA-JF_646_ and JF_635_b for RNA-FISH experiments. (**a**) SOFI RNA-FISH image of MEF cells expressing MS2 in the 3′ UTR of the beta-actin gene and labeled with JF_635_b-oligonucleotide from **5_NHS_**; scale bar: 5 μm. (**b**) SMLM RNA-FISH image of MEF cells expressing MS2 in the 3′UTR of the beta-actin gene and labeled with JF_646_b-oligonucleotide from **6_NHS_**; scale bar: 5 μm. (**c**) Zoomed area from **b**; scale bar: 1 μm. (**d,e**) Statistics for SMLM images shown in panel **b** and **Fig. 2j** histogram of uncertainty of fit (**d**), and histogram of *σ* of Gaussian fit (**e**) calculated using 2,219,274 detections for the JF_646_b-oligonucleotide from **5_NHS_**, and 2,257,059 detections for JF_635_b-oligonucleotide from **6_NHS_**. The shoulder in the *σ* of Gaussian fit plot for **5_NHS_** (JF_646_b) suggests attempted fits of multiple molecules as one event, which is consistent with the higher duty cycle of this label. Images in **a**–**c** were taken using HILO illumination with 30 ms exposure time (33.33 Hz). All imaging experiments were repeated with similar results.

**Extended Data Figure 8.**
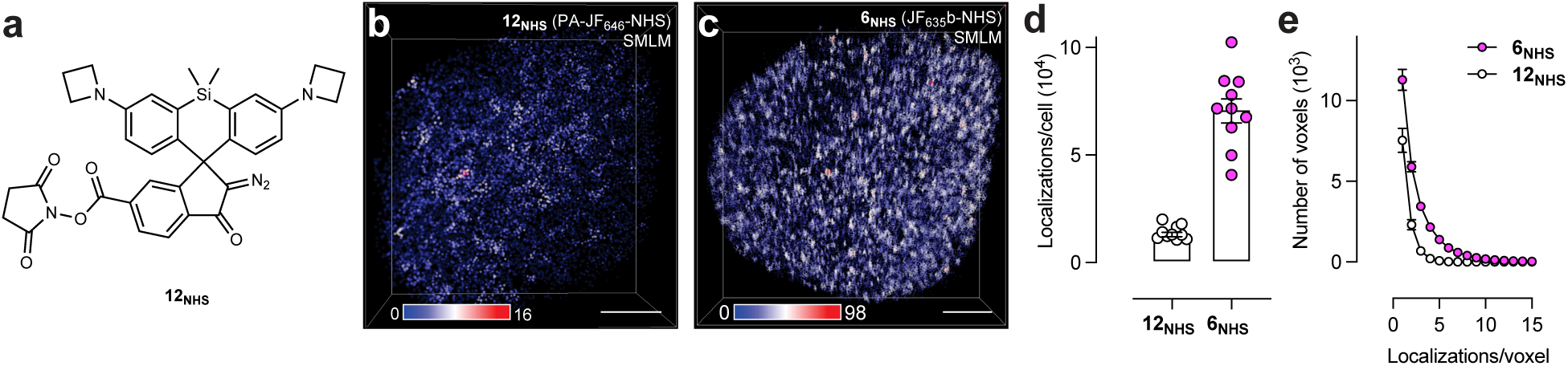
Comparison of PA-JF_646_ and JF_635_b for ATAC–SMLM experiments. (**a**) Structure of PA-JF_646_-NHS (**12_NHS_**). (**b,c**) Representative 3D ATAC–SMLM rendering of accessible chromatin localizations in wild-type mouse embryonic stem (mES) cells using oligonucleotide probes labeled with **12_NHS_** (**b**) or **6_NHS_** (**c**); the color-coded localization density was calculated with a canopy radius of 250 nm; scale bars: 2 µm. (**d**) Localizations per cell for ATAC–SMLM experiments using oligonucleotides labeled with **12_NHS_** (*n* = 10 cells) or **6_NHS_** (*n* = 10 cells); error bars indicate ±SEM. (**e**) Plot of number of voxels *vs.* localizations per voxel for ATAC–SMLM experiments using oligonucleotides labeled with **12_NHS_** (*n* = 10 cells) or **6_NHS_** (*n* = 10 cells); error bars indicate ±SEM.

## ONLINE METHODS

### Chemical synthesis

The development of a new synthetic method for Si-fluorescein derivatives including **2** is described in **Supplementary Note 1**. Experimental details and characterization for the syntheses of **3_HTL_**–**10_HTL_**, **8_HST_**, **6_MAL_**, **5_NHS_**, and **6_NHS_** can be found in **Supplementary Note 2**.

### Spectroscopy of HaloTag ligands

HaloTag ligands **3_HTL_**–**10_HTL_** were prepared as 1 mM stock solutions in DMSO. We initially attempted to measure the spectral properties of the HaloTag ligands as conjugates, but the HaloTag proteins were not stable at the low pH values necessary to promote the visible absorbing and fluorescent form of the spontaneously blinking dyes. Instead, the spectral properties of HaloTag ligands **3_HTL_**–**10_HTL_** were measured in 10 mM citrate buffer, pH 3.0, containing 150 mM NaCl and 0.1% (w/v) sodium dodecyl sulfate (SDS). The visible-absorbing form of Si-rhodamines bind to SDS micelles^4^ and, combined with the low pH, these conditions afforded solutions with measurable absorbance and fluorescence across the compound series. All measurements were taken at ambient temperature (22 ± 2 °C) using 1-cm path length, 3.5-mL quartz cuvettes (Starna Cells). Maximum absorption wavelength (*λ*_abs_) of the HaloTag ligands **3_HTL_**–**10_HTL_** were measured on a Cary Model 4000 spectrometer (Agilent). Absorption spectra are averages (*n* = 3) and normalized spectra are shown for clarity. Fluorescence spectra and maximum emission wavelength (*λ*_em_) of the HaloTag ligands **3_HTL_**–**10_HTL_** were measured on a Cary Eclipse fluorometer (Agilent); normalized spectra are shown for clarity. Absolute fluorescence quantum yield values (*Φ*) of the HaloTag ligands **3_HTL_**–**10_HTL_** were measured using a Quantaurus-QY spectrometer (model C11374, Hamamatsu). This instrument uses an integrating sphere to determine photons absorbed and emitted by a sample. Measurements were carried out using dilute samples (*A* < 0.1), and self-absorption corrections were performed using the instrument software.^24^ Reported values are averages of 10 values taken at different excitation wavelengths with two separate dye solutions.

### Cell culture and plating of cells for SMLM

COS-7 (CRL-1651), U2OS (HTB-96), and hTERT-immortalized retinal pigment epithelial cells (hTERT RPE-1; CRL-4000) were obtained from ATCC. For imaging of cell nuclei using the HaloTag system, we used COS-7 cells with an integrated HaloTag–histone H2B fusion protein expressing plasmid via the piggyBac transposon system^25^. For imaging of mitochondria, we used U2OS cells with an integrated a TOMM20–HaloTag fusion protein expressing plasmid^13^. These cell lines were kept under the selection of 500 μg/mL Geneticin (Life Technologies). For imaging the endoplasmic reticulum, we used U2OS cells stably expressing a HaloTag-Sec61b fusion protein via lentiviral transduction and integration. All cell lines undergo regular mycoplasma testing by the Legant Lab or by the Janelia Cell Culture Shared Resource. All cell lines were cultured in growth medium comprised Dulbecco’s Modified Eagle Medium (DMEM; Thermo Fisher Scientific, 11965118) supplemented with 10% (v/v) fetal bovine serum (FBS; Avantor 1300-500H) and 100 units/mL penicillin–streptomycin (Thermo Fisher Scientific, 15140122). Cell cultures were maintained in an incubator under standard conditions (37 °C, 5% (v/v) CO_2_). For all imaging experiments, cells were harvested and plated one day prior to dye labeling. U2OS cells were plated at a density of 1 × 10^5^ cells/mL and hTERT RPE-1 and COS-7 cells were plated at a density of 2.5 × 10^5^ cells/mL in 6-, 12-or 24-well glass bottomed plates (Cellvis P06-1.5H-N, P12-1.5H-N, P24-1.5H-N).

### Labeling of cells with HaloTag ligands 3_HTL_–10_HTL_

HaloTag ligands were dissolved in anhydrous dimethyl sulfoxide (DMSO; Thermo Fisher Scientific, 50-148-9333) to achieve a final concentration of 1 mM. The ligand stock solution was aliquoted and stored at −20 °C; a freshly thawed aliquot was used immediately for each experiment. The ligand stock solution was diluted into complete growth media and cells were incubated for 1 h. To achieve complete labeling of HaloTag proteins, a concentration of 0.5–2 μM of dye ligand was used. For the on-time, off-time, and duty cycle measurements, we utilized a serial dilution method to label cells with a range of different concentrations of each dye to achieve a labeling density at which multiple blinks could be unambiguously assigned to single dye molecules. After the 1 h incubation, cells were rinsed once with prewarmed phosphate-buffered saline (PBS; Corning 21-040-CMX12) and then placed in fresh growth medium for 20 min. The growth medium was refreshed after 20 minutes and incubated for an additional 20 minutes. Cells were then washed once with room temperature PBS and fixed in freshly prepared 4% paraformaldehyde (Electron Microscopy Sciences 15710) for 12 minutes. This was followed by a 10-second wash and three 5-minute washes with PBS.

Simultaneous with labeling and fixations steps, a solution of Wheat Germ Agglutinin (WGA) conjugated with nanodiamonds (WGA-ND; Adámas, NDNV1000-WGA custom order) was diluted in PBS and was sonicated for at least 1 h. Following fixation and wash steps, the cellular samples were incubated in WGA-ND solution for 20 minutes on a rocker at ambient temperature. This was followed by 10-second wash and three 5-minute washes with PBS. The sample then underwent a secondary fixation step for five minutes, followed again by a 10-second wash and three 5-minute washes with PBS. Samples were either imaged immediately following sample preparation or sealed with parafilm and stored overnight at 4 °C. For samples stored overnight, the PBS was replaced, and samples were equilibrated to room temperature prior to imaging.

### Labeling of cells with Hoechst derivative 8_HST_

Hoechst derivative **8_HST_** was dissolved in anhydrous dimethyl sulfoxide (Thermo Fisher Scientific, 50-148-9333) to achieve a final concentration of 1 mM. The ligand stock solution was aliquoted and stored at −20 °C; a freshly thawed aliquot was used immediately for each experiment. The **8_HST_** stock solution was diluted in PBS to a final concentration of 10 nM. Cells were processed as described above and then incubated with this PBS solution throughout the imaging procedure.

### SMLM imaging of HaloTag-expressing cells labeled with 3_HTL_–10_HTL_ and cells labeled with 8_HST_

All samples were imaged using a custom built highly inclined swept tile (HIST) microscope^26^, based on a Nikon TI2 platform and utilizing a 60× 1.27 numerical aperture objective (MRY10060, Nikon). Individual WGA-ND images were acquired using 11.5 mW of 560 nm laser illumination with a Semrock FF01-680/42-32 emission filter. All single molecule images were aquired using 560 nm (2RU-VFL-P-2000-560-B1R, MPB Communications Inc) and 647 nm (2RU-VFL-P-2000-642-B1R, MPB Communications Inc) laser illumination to optimally excite both the ND and dye molecules and imaged onto a camera (Fusion BT, Hamamatsu) through a bandpass emission filter (FF01-680/42, Semrock). Regions of interest (ROIs) were identified under low power illumination conditions (11.5 mW at 560 nm and 1.6 mW at 647 nm, measured at the objective back pupil). Each ROI contained at least three WGA-ND fiducial markers for saturating and blinking imaging conditions, and at least one WGA-ND fiducial for blink lifetime imaging conditions. To measure dye on-time, 20,000 frames were acquired with 2 ms exposure (3 ms cycle time) using 40 mW of 647 nm and 11.5 mW of 560 nm illumination. To measure dye off-time and for representative super-resolution renderings 100,000 frames were acquired with 25 ms exposure (27 ms cycle time) under the same illumination conditions as above.

### SMLM image processing of HaloTag-expressing cells labeled with 3_HTL_–10_HTL_ and cells labeled with 8_HST_

Individual blink events within the images were fit using the SMAP localization software package^27^ based on MATLAB (The Mathworks) with the following parameters: Peak Finder (DoG, s: 1.2, dynamic: 1.7), Fitter (PSF Fix, PSFx start: 2 pix, ROI size: 7 pix, Iterations: 25). The data were additionally filtered to keep only localizations that could be localized to <40 nm, emitted more than 100 photons per frame (for blink quantification) or 40 photons per frame (for blink lifetime measurement). Sample drift was compensated for by tracking multiple fiducial ND within the field of view and subtracting their motion over time from the localization coordinates. To quantify the photophysical properties of each dye, localizations were first assigned to individual cells using a manually outlined binary mask. To determine dye on-time, multiple localizations were considered to have come from the same dye molecule if they were observed in consecutive frames within 100 nm of a previous observation. To distinguish long-lived blinking events that occur from dye protonation/deprotonation from transient events that may be due to the Poisson nature of photon emission or noise, a “gap-closing” parameter of 5 frames (15 ms) was used such that if an “on” molecule transiently drops below our detection threshold and reappears within 5 frames, it is still considered the same on event. To determine dye off-time, sparse labeling was used such that groups of localizations could be unambiguously assigned to a single dye molecule even if the localizations occurred at distant time points. All localizations acquired within 100,000 frames (45 minutes of imaging) were plotted and groups were identified using the “dbscan” function with a grouping radius of 100 nm. Off-time events for each molecule were quantified as the intervening time experienced between localizations for each grouping.

### Super-resolution optical fluctuation imaging (SOFI) using HaloTag ligands

U2OS cells were Amaxa-transfected with 500 ng plasmid per 1 × 10^6^ cells encoding for N-terminally labeled HaloTag-ensconsin fusion protein. The cells were plated on plated in 35 mm glass-bottom MatTek dishes and allowed to grow to appropriate density (5 × 10^4^ cells/cm^2^) in phenol-free DMEM containing 10% (v/v) FBS for imaging. Cells were labeled with 100 nM of Halo-ligand **4_HTL_**, **6_HTL_**, or **8_HTL_** for 30 min. Cells were washed three times with media and then fixed with a solution of 4% (w/v) paraformaldehyde in PBS buffer, pH 7.5. The fixed cells were imaged in 1× PBS buffer at room temperature with an Olympus 100× 1.5 NA TIRF objective on a RAMM frame microscope (ASI), equipped with a tube lens (LAO-300.0, Melles Griot), resulting in 166.66× overall magnification^28^. Emitted light was collected on a Andor iXon Ultra EMCCD camera with increased sensitivity in the NIR (model DU-897UCS0-EXF) that was operated using Micro-Manager^29^ version 1.4.20 with the following settings: gain 400, cooled to −60 °C, 17 MHz EM amplifiers, preamp setting 3. The sample was illuminated using a Stradus 637-140 laser (Vortran) illuminated the sample in a highly inclined and laminated optical sheet (HILO) configuration with an illumination power density of 400 W/cm^2^. 20,000 frames were recorded at 100 frames/s, and a subset of 5000 frames were analyzed with Igor Pro v9.05 (WaveMetrics) using the Localizer plugin^30^ to reconstruct second-order SOFI images. SMLM images were calculated from the full 20,000 frames using the ThunderSTORM plugin^31^ for Fiji^32^ with maximum likelihood integrated gaussian fitting option. SMLM images were visualized using normalized gaussians.

### Expression and purification of FUS variants

MBP-FUS-EGFP was overproduced from pMBP-*tev*-FUS-EGFP-*tev*-His_6_ [*aka* pMal-*Tev*-FUS(WT)-EGFP-*Tev*-His_6_] and purified as described previously^33^. pMBP-*tev*-FUS-*tev*-His_6_ and pMBP-*tev*-FUS^A2C^-*tev*-His_6_ were generated by PCR amplifying the FUS coding sequence in pMBP-*tev*-FUS-EGFP-*tev*-His_6_ with SalI/HindIII ends without or with the A2C mutation and ligating the digested fragment into the original plasmid digested with SalI/HindIII, which removes the FUS-EGFP coding sequence. Inserted sequences were confirmed by DNA sequencing. Plasmids pMBP-*tev*-FUS-*tev*-His_6_ and pMBP-*tev*-FUS^A2C^-*tev*-His_6_ were used to overproduce MBP-FUS and MBP-FUS^A2C^, respectively, which were purified identically as MBP-FUS-EGFP, except as indicated. Plasmids pMBP-*tev*-FUS-*tev*-His_6_ and pMBP-*tev*-FUS^A2C^-*tev*-His_6_ were submitted to Addgene, and their construction is described in the history of the linked SnapGene files.

All FUS-containing proteins were overproduced in *E. coli* strain Rosetta 2(DE3)pLysS (Novagen) transformed with the appropriate plasmid using lysogeny broth (LB) with 50 µg/mL ampicillin. After incubating a 5 mL starter culture overnight at 37 °C, it was centrifuged (4000×*g*, 5 min, 25 °C) and resuspended in 1 L fresh media, which was incubated at 37 °C. At an OD_600_ of ∼0.8, protein overproduction was induced with 1 mM isopropyl β-D-1-thiogalactopyranoside (IPTG) for 24 h at 20 °C with shaking at 250 rpm before pelleting the culture (4,000×*g*, 20 min, 4°C). Cells were recovered (4,000×*g*, 20 min, 4 °C) and resuspended in 25 mL resuspension buffer consisting of 50 mM Na_2_HPO_4_/NaH_2_PO_4_, 500 mM NaCl, 10 μM ZnCl_2_, 10 mM imidazole, 4 mM β-mercaptoethanol (βME), 10% glycerol, pH 7.5, and the protease inhibitors: 10 mM phenylmethane sulfonyl fluoride (PMSF), 100 µg/mL trypsin inhibitor, 20 µg/mL leupeptin, and 100 µg/mL pepstatin. The suspension was lysed by French press (3× at 16,000 psi), the lysate was centrifuged (15,000×*g*, 20 min, 4 °C), and the supernatant was incubated with 500 µL of Ni-NTA resin (Qiagen) for 30 min at 25 °C. The suspension was transferred to a gravity column. The resin was washed with 10 column volumes (CVs) of resuspension buffer with 2 mM PMSF, followed by 5 CV of wash buffer A consisting of: 50 mM Na_2_HPO_4_/NaH_2_PO_4_, 500 mM NaCl, 10 μM ZnCl_2_, 10 mM imidazole, 4 mM βME, 10% glycerol, pH 7.5. Fractions (1 mL) were eluted with elution buffer A consisting of: 50 mM Na_2_HPO_4_/NaH_2_PO_4_, 500 mM NaCl, 10 μM ZnCl_2_, 400 mM imidazole, 4 mM βME, 10% (v/v) glycerol, pH 7.5. Protein-containing fractions (<15 mL) were combined and diluted with 100 mL dilution buffer consisting of: 50 mM Na_2_HPO_4_/NaH_2_PO_4_, 150 mM NaCl, 10 μM ZnCl_2_, 10 mM imidazole, 10% (v/v) glycerol, 4 mM βME, pH 7.5. After gentle shaking at room temperature for 5 min, the protein mixture was loaded onto a gravity flow column with 2 mL amylose resin (New England Biolabs, #E8021). The resin was washed with 5 CVs of wash buffer B consisting of: 50 mM Na_2_HPO_4_/NaH_2_PO_4_, 200 mM NaCl, 10 μM ZnCl_2_, 10 mM imidazole, 10% (v/v) glycerol, pH 7.5, and the protein was eluted with amylose elution buffer consisting of: 20 mM Tris, 150 mM NaCl, 20 mM maltose, 1 mM EDTA, 5% (v/v) glycerol, pH 7.5. For MBP-FUS^A2C^, 2 mM tris(2-carboxyethyl)phosphine (TCEP) was added before storage. The eluted protein was typically ∼20–30 µM, and it was flash frozen in liquid nitrogen and stored at −80°C in 10 µL aliquots until use. Protein concentrations were determined by the densitometry of bands on SDS-PAGE gels stained with Coomassie Blue R-250 using carbonic anhydrase as a standard and a ChemiDoc MP imaging system (Bio-Rad Laboratories). The purity of dye-labeled proteins was assayed by direct in-gel fluorescence imaging using the same ChemiDoc imaging system and was determined to be >95%.

### Labeling of FUS variants

MBP-FUS^A2C^ was reacted with a 15-fold molar excess of the spontaneously blinking dye HM-SiR-maleimide^7^ (**1_MAL_**; *aka* SaraFluor 650B-maleimide; Goryo Chemical, Japan, #A209-01) or JF_635_b-maleimide (**6_MAL_**) at room temperature for 15 min. The reaction was quenched with 10 mM βME. The dye-protein mixture was incubated with 0.2 mL Ni-NTA resin for 15 min and loaded into a gravity flow column. Excess dye was removed by washing the resin-bound protein with 100 CVs of wash buffer C consisting of: 50 mM Na_2_HPO_4_/NaH_2_PO_4_, 10 mM imidazole, 10 µM ZnCl_2_, 0.5 M NaCl, 0.2% (v/v) Triton X-100, 1 mM dithiothreitol (DTT), 10% (v/v) glycerol, pH 7.5, and 50 CVs of wash buffer D consisting of: 50 mM Na_2_HPO_4_/NaH_2_PO_4_, 10 mM imidazole, 10 µM ZnCl_2_, 150 mM NaCl, 2 M urea, 1 mM DTT, 10% (v/v) glycerol, pH 7.5. To assess removal of the free dye, Ni-NTA resin (10 µL) was incubated with reducing SDS sample buffer and the protein was electrophoresed. The acrylamide gel was incubated in ∼5% (v/v) acetic acid (pH 2–3) for 5 min to convert the blinking dye molecules into the protonated fluorescent state, and the sample was analyzed by direct in-gel fluorescence imaging (Epi Red, *λ*_ex_ = 625–650 nm). If the free dye was still present, the resin was washed again and re-assayed. Once free dye was no longer detected in the gel image (< 2%), the protein was eluted with elution buffer B consisting of: 50 mM Na_2_HPO_4_/NaH_2_PO_4_, 250 mM imidazole, 10 µM ZnCl_2_, 150 mM NaCl, 10% (v/v) glycerol, pH 7.5. The eluted protein was typically 1–2 µM, and it was flash frozen in liquid nitrogen and stored at −80°C in 3 µL aliquots until use.

### FUS condensate formation and sample preparation

FUS protein stock solutions were thawed at room temperature and centrifuged (14,000×*g*, 10 min, 25 °C). Supernatants were used immediately. MBP-FUS or MBP-FUS-EGFP solutions (diluted to 14 µM) were mixed with MBP-FUS^A2C^-JF_635_b (40 or 4 nM) in droplet buffer A consisting of: 20 mM Na_2_HPO_4_/NaH_2_PO_4_, 2.5% (v/v) glycerol, 1 mM DTT, 150 mM NaCl, pH 7.5. Protein mixtures were then diluted 2-fold with no salt droplet buffer B consisting of: 20 mM Na_2_HPO_4_/NaH_2_PO_4_, 2.5% (v/v) glycerol, 1 mM DTT, pH 7.5 to yield a final salt concentration of 75 mM and 7 µM MBP-FUS or MBP-FUS-EGFP. Phase separation was induced by the addition of 0.3 µL (3 U) of acTEV protease (10 U/µL; Invitrogen, #12575015) to 10 µL of the FUS protein mixture, which resulted in proteolytical removal of the MBP solubility tag and the 6×His-tag on the various FUS constructs.^33^ After incubation at room temperature for 10 min, the solution was added to a microscope slide flow chamber (∼10 µL), which was then sealed with clear nail polish to avoid evaporation. Flow chambers were made from large coverslips (Gold Seal 3243; 24×60 mm; ThermoFisher Scientific, #50-189-9138) that were argon plasma cleaned and pretreated with 1.5% mPEG-silane (LaysanBio, #MPEG-SIL-5000) for 30 min at room temperature to passivate the surface, followed by 3×10 µL washes with droplet buffer. Small coverslips (10.5×22 mm; Electron Microscopy Sciences, #72191-22) with double-sided tape on their short edges were pasted on top of a large coverslip to construct flow chambers.

### 3D#super-resolution microscopy of FUS condensates

The set-up, sensitivity, and precision of our microscope system using a tunable astigmatism to obtain three-dimensional (3D) spatial information was described previously.^10, 23^ Briefly, a Zeiss Axiovert 200M equipped with an oil-immersion objective (Zeiss Alpha Plan-Apochromat, 1.40 NA) and sCMOS camera (Teledyne Photometrics, Prime 95B, square pixel dimension = 120 nm at the camera plane) was used to capture the fluorescence emission from individual HM-SiR or JF_635_b molecules during their rare on-time events within FUS condensates. Laser excitation was converted to circularly polarized light with half (*λ*/2) and a quarter (*λ*/4) waveplates (Newport, #10RP42-1 and #10RP44-1) before entering the microscope as a slightly converging illumination beam. A self-configured adaptive optics system (Imagine Optic, AOKit Bio) incorporated into the emission path generated the *z*-dependent spot ellipticity needed for 3D information using 60 nm root mean square (rms) astigmatism. Green and red emission channels were collected with a double-bandpass filter set (Chroma, ZT532/640/NIR-RPC-UF2). A self-configured TIRF-Lock system (Mad City Labs) provided a *z*-stability for the coverslip of <3 nm for the duration of the experiment. The GFP fluorescence in FUS-EGFP condensates was acquired with 488 nm diode laser illumination (Coherent, Saphhire 488 LPX, 400 mW) with ∼0.05 kW/cm^2^ at the sample.

### Density maps from 3D localizations in FUS condensates

To generate 3D density maps, condensates were formed with 20 nM FUS^A2C^-JF_635_b. Localizations (one localization/blink) were acquired with 647 nm diode laser illumination (Coherent, OBIS 647, 120 mW) with ∼10 kW/cm^2^ at the sample and 30 ms/frame for 20 min. Image sequences were analyzed using the ThunderSTORM plug-in^31^ for Fiji.^32^ The maximum likelihood method was used to determine all single molecule spot centroids (*x*, *y*) and the “elliptical gaussian” fitting option was used to fit the astigmatic point spread functions (PSFs), which yielded the *z*-positions. The calibration file required for the *z*-dependent PSF was obtained from *z*-stacks (−500 nm to +500 nm; 25 nm steps) of 0.1 µm fluorescent microspheres (ThermoFisher Scientific, T7279) and were fit as described by Huang *et al.*^2^ and implemented in the ThunderSTORM plug-in. The post-processing option “remove duplicates” was used to eliminate overlapping molecules and “density filters” was used to estimate the number of spots within a sphere around each localization (search radius = 100 nm). The estimated local spot densities are indicated by the color-coding in **Fig. 2b,c**.

### SiMRoD maps from 3D localizations in FUS condensates

FUS condensates mature over time from a high mobility liquid state to a low mobility solid state.^34, 35^ This phase maturation can, in principle, be detected by the change in rotational mobility of a fluorescent dye attached to FUS molecules. In our case, the JF_635_b dye was attached via a flexible linker to an intrinsically disordered region of FUS.^36^ Thus, rotational mobility measurements were expected to depend more heavily on the local environment of the dye and less on the mobility of the macromolecule to which it was attached, although these mobilities could be related. Probes within a completely solid structure attached to the surface of the chamber are expected to be rotationally immobile on any timescale. In contrast, probes within a large solid assembly (*i.e.*, an aggregated particle) diffusing in a viscous but liquid-like medium would likely be rotationally immobile in a fluorescence anisotropy experiment but have rotational mobilities distinct from a completely solid structure. Single molecule rotational diffusion (SiMRoD) microscopy, originally reported as polarization PALM or “p-PALM”^19^ and renamed here, can distinguish such slow rotational mobilities. Since SiMRoD is a relatively new technique, we first describe the basic principles of the approach before describing the measurements on FUS condensates.

In SiMRoD, the fluorescence emission from a single fluorescent molecule is split with a 50% polarizing beam splitter and both emission channels are simultaneously recorded.^19^ Single molecule polarization values, *p*, are calculated from the spot intensity pairs using equation 1:

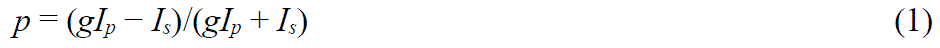

where *I*_p_ and *I*_s_ are the intensities (photons) measured in the two detection channels, and *g* is a system-dependent factor that corrects for differences in photon collection efficiency of the *p*– and *s*-channels. Each individual polarization value estimates the time average of the polarizations from the rotational trajectory of the individual molecule during the acquisition time. Thus, *p* values near zero are expected for fast rotational motion relative to the image collection timescale. In contrast, *p* values ranging from −1 to 1 are expected for immobile molecules. In SiMRoD, average rotational mobility information is extracted not from the average bulk polarization, but from polarization measurements obtained from thousands of individual molecules that are pooled to generate polarization frequency histograms (**Extended Data** Fig. 5f). The primary experimental readouts from these data are the average polarization,

<*p*>, and the variance of the polarization distribution, Var(*p*). The overall shape of polarization histograms and photon scatterplots can provide additional clues as to the underlying physical constraints on the probe’s rotational mobility.^19^ Var(*p*) provides a measure of the width of polarization histograms, and is formally calculated using equation

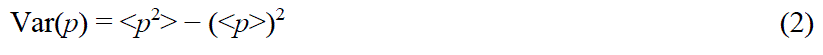

where <*p*^2^> is the average square of the measured polarization values. For a properly calibrated system using circularly polarized excitation,

<*p*> = 0 and hence Var(*p*) = <*p*^2^>. Thus, we assumed that Var(*p*)_cir_ was equivalent to the average of the squares of the experimentally determined *p* values, <*p*^2^>_cir_. The requirement that

<*p*>_cir_ = 0 (except under unusual circumstances) provides a convenient internal check for each dataset on instrument alignment and calibration.

For SiMRoD experiments in condensates containing FUS^A2C^-JF_635_b, the emission from a single emitter was split with a 50% polarizing beam splitter cube (PBS201, ThorLabs, Newton, NJ) mounted in an Optosplit III beamsplitter (Cairn Research, Kent, UK), which allowed the two polarization components to be imaged simultaneously on the two halves of the sCMOS camera. To estimate *g*, we measured the intensities of 0.1 µm fluorescent microspheres (ThermoFisher Scientific, #T7279) embedded in 2% agarose. Assuming that the dyes were randomly oriented in the microspheres, <*p*>_cir_ was expected to be 0. Under these conditions, we found that *g* = <*I*_s_/*I*_p_> = 0.94 (from equation 1, assuming <*p*> = 0). Since SiMRoD depends only on the intensity (photons) measured in the two channels, it is insensitive to the astigmatic PSFs or minor spatial misalignments of the two channels. For obtaining spatial coordinates, the brighter spot in the two channels for each molecule was used. Data were collected at 5 ms/frame for 8 min using ∼50 kW/cm^2^ (*λ*_ex_ = 647 nm), measured at the sample plane. SiMRoD heatmaps were generated using an Excel script and Origin 8.5 by calculating Var(*p*)_cir_ within a 400 nm search radius assuming a minimum of 20 polarization values and a minimum of 500 photons in one of the channels.

### Labeling of oligonucleotides for RNA-FISH

The oligonucleotides used to prepare RNA-FISH probes were synthesized by Integrated DNA Technologies with a 5′ primary amino group and a six-carbon spacer arm (C6). The oligonucleotide sequences and their modifications are:

5′-amino-C6-TTTCTAGAGTCGACCTGCAG-3′

5′-amino-C6-CTAGGCAATTAGGTACCTTAG-3′

5′-amino-C6-CTAATGAACCCGGGAATACTG-3′

The oligonucleotides (500 nmol) were dissolved in 500 µL deionized water and extracted three times with chloroform. The aqueous solution was separated, and the oligonucleotide was precipitated by adding 50 µL of 3 M NaCl(aq) and 1250 µL EtOH and incubating at −20 °C for 30 min. The sample was centrifuged (13,000 rpm, 30 min) and the resulting precipitate was washed with 70% (v/v) EtOH and dried. Purified oligonucleotides (17 nmol) were dissolved in 0.1 M sodium tetraborate, pH 8.5 (36.5 µL) and **11_NHS_**, **5_NHS_**, or **6_NHS_** (25 nmol) in DMSO (3.5 µL) was added. The reaction was incubated for 48 h at ambient temperature with constant shaking and protection from light. The oligonucleotide was precipitated by adding 10 µL of 3 M NaCl(aq) and 250 µL EtOH and incubating at −20 °C for 30 min. The sample was centrifuged (13,000 rpm, 30 min) and the resulting precipitate was washed twice with 70% (v/v) EtOH ethanol and then dissolved in deionized water. Purification of labeled oligonucleotides was performed on Agilent 1200 high-performance liquid chromatography (HPLC) purification system equipped with an autosampler, diode array detector and fraction collector. The sample was introduced onto a 4.6 × 150 mm, 5 μm, Eclipse XDB-C18 column (Agilent) using the system autosampler and eluted with a linear gradient of 0→100% (v/v) CH_3_CN/H_2_O (10 min, 3 mL/min) containing a constant 10 mM TEAA additive. The purification was monitored using the system diode array detector (*λ*_abs_ = 260 nm) and collected using the system fraction collector. Product-containing fractions were pooled and lyophilized to obtain the label oligonucleotide products as white solids.

### Cell culture and sample preparation for RNA-FISH experiments

The protocol for RNA-FISH using immortalized mouse embryo fibroblasts (MEFs) was described previously.^37^ Briefly, MEF cells derived from a transgenic mouse containing a MS2 binding site (MBS) cassette targeted to the 3′ untranslated region of the essential beta-actin gene were cultured in growth medium comprised Dulbecco’s Modified Eagle Medium (DMEM; Thermo Fisher Scientific, 11965118), supplemented with 10% (v/v) fetal bovine serum (FBS; Avantor 1300-500H) and 100 units/mL penicillin–streptomycin (Thermo Fisher Scientific, 15140122). Cell cultures were grown on MatTek dishes to 50% confluency. Cells were washed with PBSM consisting of PBS supplemented with 5 mM MgCl_2_, and then fixed in 4% (w/v) paraformaldehyde for 10 min at ambient temperature. The cells were then washed with PBSM (3×5 mins) and then with PBSM supplemented with 50 mM glycine (1×5 mins). Before hybridization, the cells were permeabilized by incubation with 0.5% (w/v) Triton X-100 in PBS for 10 min, washed with PBSM (3×5 min), and then incubated for 10 min in prehybridization solution consisting of: 10% (v/v) formamide, 10% (w/v) dextran sulfate, 2× saline-sodium citrate (SSC) buffer, 0.2 mg/mL BSA, 1 mg/mL *Escherichia coli* tRNA, 2 mM ribonucleoside vanadyl complexes (New England Biolabs), and 10 U/mL Superase (Invitrogen). The cells were then incubated with 200 µL prehybridization solution supplemented with 20 ng DNA probe at 37 °C for 18 h. The cells were washed with a solution of 10% v/v formamide in 2× SSC buffer at 37 °C (2×15 mins), and then washed with 2× SSC buffer at ambient temperature (3×10 mins).

### Fluorescence microscopy for RNA-FISH experiments

All samples were imaged at room temperature in PBS buffer. Widefield, SOFI, and SMLM experiments were performed in an ELYRA system (Carl Zeiss) with a 63×/1.4 NA Plan-Apo objective augmented with a Zeiss 1.6× magnification lens resulting in a pixel size of 158.7 nm. The system contained an EMCCD camera (Andor Ultra 897, Gain 300) able to detect single photons, and the samples were illuminated with a 642 nm excitation laser in a HILO configuration at a frame rate of 33.33 Hz with a constant illumination power density of 100 W/cm^2^ for widefield and SOFI experiments and 300 W/cm^2^ for SMLM experiments. Second-order SOFI images were reconstructed using Igor Pro 7 software (WaveMetrics) with the Localizer plugin^30^ using a 2000-frame movie. SMLM experiments were performed as previously described.^4, 8^ For correction of sample drift, 100-nm TetraSpeck Microspheres (Invitrogen) were affixed to the cover-glass by incubation in the medium for 30 min at room temperature. SMLM images were generated using the ThuderSTORM plug-in^31^ for Fiji^32^ from a 5000– 20,000-frame movie with maximum likelihood integrated gaussian fitting option. SMLM images were visualized using normalized gaussians.

### Labeling of oligonucleotides for 3D ATAC–SMLM experiments

The 3D ATAC–SMLM experiments including purification of the Tn5 transposase followed our previous protocol^22^. The mosaic ends (ME) adaptors for Tn5 transposase were synthesized by Integrated DNA Technologies with attachment of a 5′ end primary amino group by a six-carbon spacer arm (C6). The oligonucleotide sequences and their modifications are:

Tn5ME-A: 5′-amino-C6 TCGTCGGCAGCGTCAGATGTGTATAAGAGACAG3′

Tn5ME-B: 5′-amino-C6-GTCTCGTGGGCTCGGAGATGTGTATAAGAGACAG-3′

Tn5MErev: 5′-(phos)CTGTCTCTTATACACATCT-3′

The *N*-hydroxysuccinimide (NHS) ester PA-JF_646_ (PA-JF_646_-NHS, **12_NHS_**)^8^ or JF_635_b-NHS (JF_635_b-NHS, **6_NHS_**) was dissolved in anhydrous DMSO immediately before conjugation. The Tn5ME-A and Tn5ME-B oligos were first dissolved in deionized water and then extracted three times with chloroform. The upper aqueous solution was carefully extracted and precipitated by 3 M sodium acetate (pH 5.2) and EtOH. The pellet was washed with 70% (v/v) EtOH, dried and dissolved in ultrapure H_2_O. The purified oligonucleotides were reacted with excess **12_NHS_** or **6_NHS_** or in anhydrous DMSO (mass ratio 1:2.5 oligo:dye) in 0.1 M sodium tetraborate, pH 8.5. The reaction was incubated at room temperature overnight (>12 h) with constant stirring and protection from light. The crude labeled oligonucleotides were precipitated by addition of 3 M sodium acetate, pH 5.2 and ethanol. The solid was collected by centrifugation and the pellet was dissolved in 0.1 M TEAA (triethylammonium acetate; Thermo Fisher Scientific, 400613). Purification of oligonucleotide-dye conjugates was performed on an Agilent 1200 analytical high-performance liquid chromatography (HPLC) system equipped with an autosampler, diode array detector and fraction collector. The column used was Eclipse XDB-C18 column (4.6 × 150 mm, 5 µm; Agilent) and eluted with a linear gradient of 0–100% (v/v) CH_3_CN/H_2_O with constant 10 mM TEAA additive; 30 min run; 1 mL/min, detection at 260 nm. Sample fractions were pooled and lyophilized to obtain the product as colorless solid.

### 3D#ATAC–SMLM

We assembled the Tn5 transposome with dye-labeled oligonucleotide duplex according to previously published work^22^. Briefly, the HPLC-purified Tn5ME-A-dye or Tn5ME-B-dye oligonucleotides were dissolved in H_2_O to yield stock solutions of 100 μM. These oligonucleotides were then annealed to the Tn5MErev oligonucleotide by adding an equimolar amount of Tn5ME-A-dye or Tn5ME-B-dye to a solution of Tn5MErev in 10 mM Tris-HCl, 50 mM NaCl, 1 mM EDTA, pH 8.0. These solutions were denatured on a benchtop thermocycler at 95 °C for 5 min and then slowly cooled down to 25 °C at the rate of −1 °C /min. We prepared wild-type mouse embryonic stem (mES) cells for 3D ATAC-SMLM experiments as reported previously^22^. Briefly, cells were plated onto 5 mm coverslips (Warner Instruments, Cat. 64-0700) at around 80% confluency with proper coating one day before experiment. Cells were fixed with 4% paraformaldehyde (Electron Microscopy Sciences, Cat. 15710) for 10 min at room temperature. After fixation, cells were washed three times with 1× PBS for 5 min and then permeabilized with ATAC lysis buffer (10 mM Tris-Cl, pH 7.4, 10 mM NaCl, 3 mM MgCl_2_, 0.1% (v/v) Igepal CA-630) for 10 min at room temperature. After permeabilization, the slides were washed twice in 1× PBS and put inside a humidity chamber box at 37 °C. The transposase mixture solution (1× Tagmentation buffer: 10 mM Tris-HCl, pH 7.6, 5 mM MgCl_2_, 10% (v/v) dimethylformamide, 100 nM Tn5-dye conjugated oligo complex) was added to the cells and incubated for 30 min at 37 °C inside the humidity chamber. After the transposase reaction, slides were washed three times with 1× PBS containing 0.01% (w/v) SDS and 50 mM EDTA for 15 min at 55 °C before mounted onto the lattice light-sheet microscope slot for ATAC–SMLM imaging.

The 3D ATAC–SMLM data were acquired by the lattice light-sheet microscopy at room temperature^38^. The light sheet was generated from the interference of highly parallel beams in a square lattice and dithered to create a uniform excitation sheet. The inner and outer numerical apertures of the excitation sheet were set to be 0.44 and 0.55, respectively. A variable-flow peristaltic pump (Fisher Scientific, Cat. 13-876-1) was used to connect a 2 L reservoir with the imaging chamber with 1× PBS circulating through at a constant flow rate. Labelled cells seeded on coverslips were placed into the imaging chamber and each imaging volume took 100–200 image frames, depending on the depth of the field of view. The specimen was illuminated using a custom 0.65 NA excitation objective (Special Optics, Wharton, NJ) and emitted light was collected by a detection objective (CFI Apo LWD 25×W, 1.1 NA, Nikon), filtered through a 440/521/607/700nm BrightLine quad-band bandpass filter (Semrock) and N-BK7 Mounted Plano-Convex Round cylindrical lens (f = 1000mm, Ø 1’, Thorlabs, Cat. LJ1516RM), and recorded by an ORCA-Flash 4.0 sCMOS camera (Hamamatsu, Cat. C13440-20CU). The cells were imaged under sample scanning mode and the dithered light sheet at 500 nm step size, thereby capturing a volume of ∼25µm × 51µm × (27–54) µm, considering the 32.8° angle between the excitation direction and the stage moving plane. The stained samples were initially photobleached by scanning the whole imaging volume with a 2 W 640 nm laser (MPB Communications Inc., Canada) to achieve sparse labelling. For the photoactivatable PA-JF_646_ label, the samples were imaged by iteratively photoactivating each plane with low-intensity 405 nm light (<0.05 mW power at the rear aperture of the excitation objective; 6 W/cm^2^ power at the sample) for 8 ms and by alternatively exciting each plane with a 2 W 640 nm laser at its full power (26 mW power at the rear aperture of the excitation objective and 3466 W/cm^2^ power at the sample) for 20 ms exposure time. For the spontaneously blinking JF_635_b label, the samples were imaged by alternatively exciting each plane with 640 nm laser at its full power for 20 ms exposure time.

To analyze the 3D ATAC–SMLM data, nano-gold fiducials were embedded within the coverslips for drift correction as previously described^22^. ATAC–SMLM images were taken to construct a 3D volume when the sample was moving along the “s” axis. Individual volumes per acquisition were automatically stored as TIFF stacks, which were then analyzed by in-house developed scripts in MATLAB. The cylindrical lens introduced astigmatism in the detection path and recorded each isolated single molecule with its ellipticity, thereby encoding the 3D position of each molecule relative to the microscope focal plane. All processing was performed by converting all dimensions to units of x–y pixels, which were 100 nm × 100 nm after transformation due to the magnification of the detection objective and tube lens. We estimated the localization precision by calculating the standard deviation of all the localizations coordinates (x, y and z) after the nano-gold fiducial correction. The localization precision was 26 ± 3nm and 53 ± 5nm for x–y and z, respectively.

## REFERENCES

1. Betzig, E. et al. Imaging intracellular fluorescent proteins at nanometer resolution. Science 313, 1642–1645 (2006).

2. Huang, B., Wang, W., Bates, M. & Zhuang, X. Three-dimensional super-resolution imaging by stochastic optical reconstruction microscopy. Science 319, 810–813 (2008).

3. Sahl, S.J., Hell, S.W. & Jakobs, S. Fluorescence nanoscopy in cell biology. Nat. Rev. Mol. Cell Biol. 18, 685–701 (2017).

4. Zheng, Q. et al. Rational design of fluorogenic and spontaneously blinking labels for super-resolution imaging. ACS Cent. Sci. 5, 1602–1613 (2019).

5. Tyson, J. et al. Extremely bright, near-IR emitting spontaneously blinking fluorophores enable ratiometric multicolor nanoscopy in live cells. ACS Cent. Sci. 7, 1419–1426 (2021).

6. Chi, W., Tan, D., Qiao, Q., Xu, Z. & Liu, X. Spontaneously blinking rhodamine dyes for single-molecule localization microscopy. Angew. Chem. Int. Ed. Engl. 62, e202306061 (2023).

7. Uno, S. et al. A spontaneously blinking fluorophore based on intramolecular spirocyclization for live-cell super-resolution imaging. Nat. Chem. 6, 681–689 (2014).

8. Grimm, J.B. et al. Bright photoactivatable fluorophores for single-molecule imaging. Nat Methods 13, 985–988 (2016).

9. Heilemann, M., van de Linde, S., Mukherjee, A. & Sauer, M. Super-resolution imaging with small organic fluorophores. Angew. Chem. Int. Ed. Engl. 48, 6903-6908 (2009).

10. Chowdhury, R., Sau, A. & Musser, S.M. Super-resolved 3D tracking of cargo transport through nuclear pore complexes. Nat. Cell Biol. 24, 112–122 (2022).

11. Grimm, J.B. et al. A general method to improve fluorophores for live-cell and single-molecule microscopy. Nat. Methods 12, 244–250 (2015).

12. Grimm, J.B. et al. A general method to fine-tune fluorophores for live-cell and in vivo imaging. Nat. Methods 14, 987–994 (2017).

13. Grimm, J.B. et al. A general method to optimize and functionalize red-shifted rhodamine dyes. Nat. Methods 17, 815–821 (2020).

14. Grimm, J.B. et al. A general method to improve fluorophores using deuterated auxochromes. JACS Au 1, 690–696 (2021).

15. Dertinger, T., Colyer, R., Iyer, G., Weiss, S. & Enderlein, J. Fast, background-free, 3D super-resolution optical fluctuation imaging (SOFI). Proc. Natl. Acad. Sci. U. S. A. 106, 22287–22292 (2009).

16. Deschout, H. et al. Complementarity of PALM and SOFI for super-resolution live-cell imaging of focal adhesions. Nat. Commun. 7, 13693 (2016).

17. Alberti, S. & Hyman, A.A. Biomolecular condensates at the nexus of cellular stress, protein aggregation disease and ageing. Nat. Rev. Mol. Cell. Biol. 22, 196–213 (2021).

18. Handwerger, K.E., Cordero, J.A. & Gall, J.G. Cajal bodies, nucleoli, and speckles in the *Xenopus* oocyte nucleus have a low-density, sponge-like structure. Mol. Biol. Cell 16, 202–211 (2005).

19. Fu, G., Tu, L.-C., Zilman, A. & Musser, S.M. Investigating molecular crowding within nuclear pores using polarization-PALM. eLife 6, e28716 (2017).

20. Ahlers, J. et al. The key role of solvent in condensation: Mapping water in liquid-liquid phase-separated FUS. Biophys. J. 120, 1266–1275 (2021).

21. Wu, B., Chao, J.A. & Singer, R.H. Fluorescence fluctuation spectroscopy enables quantitative imaging of single mRNAs in living cells. Biophys. J. 102, 2936–2944 (2012).

22. Xie, L. et al. 3D ATAC-PALM: Super-resolution imaging of the accessible genome. Nat. Methods 17, 430–436 (2020).

23. Chowdhury, R., Sau, A., Chao, J., Sharma, A. & Musser, S.M. Tuning axial and lateral localization precision in 3D super-resolution microscopy with variable astigmatism. Opt. Lett. 47, 5727–5730 (2022).

24. Suzuki, K. et al. Reevaluation of absolute luminescence quantum yields of standard solutions using a spectrometer with an integrating sphere and a back-thinned CCD detector. Phys. Chem. Chem. Phys. 11, 9850–9860 (2009).

25. Grimm, J.B. et al. Optimized red-absorbing dyes for imaging and sensing. J. Am. Chem. Soc. 145, 23000–23013 (2023).

26. Tang, J. & Han, K.Y. Extended field-of-view single-molecule imaging by highly inclined swept illumination. Optica 5, 1063–1069 (2018).

27. Ries, J. SMAP: A modular super-resolution microscopy analysis platform for SMLM data. Nat. Methods 17, 870–872 (2020).

28. English, B.P. & Singer, R.H. A three-camera imaging microscope for high-speed single-molecule tracking and super-resolution imaging in living cells. Proc. SPIE Int. Soc. Opt. Eng. 9550, 955008 (2015).

29. Edelstein, A., Amodaj, N., Hoover, K., Vale, R. & Stuurman, N. Computer control of microscopes using microManager. Curr. Protoc. Mol. Biol. **Chapter** 14, 20 (2010).

30. Dedecker, P., Duwe, S., Neely, R.K. & Zhang, J. Localizer: Fast, accurate, open-source, and modular software package for superresolution microscopy. J. Biomed. Opt. 17, 126008 (2012).

31. Ovesný, M., Křížek, P., Borkovec, J., Švindrych, Z. & Hagen, G.M. ThunderSTORM: A comprehensive ImageJ plugin for PALM and STORM data analysis and super-resolution imaging. Bioinformatics 30, 2389–2390 (2014).

32. Schindelin, J., et al. Fiji: An open-source platform for biological-image analysis. Nat. Methods 9, 676-682 (2012).

33. Hofweber, M. et al. Phase separation of FUS is suppressed by its nuclear import receptor and arginine methylation. Cell 173, 706–719 (2018).

34. Patel, A. et al. A liquid-to-solid phase transition of the ALS protein FUS accelerated by disease mutation. Cell 162, 1066–1077 (2015).

35. Qamar, S. et al. FUS phase separation is modulated by a molecular chaperone and methylation of arginine cation-pi interactions. Cell 173, 720–734 (2018).

36. Burke, K.A., Janke, A.M., Rhine, C.L. & Fawzi, N.L. Residue-by-residue view of in vitro FUS granules that bind to C-terminal domain of RNA polymerase II. Mol. Cell 60, 231–241 (2015).

37. Lionnet, T. et al. A transgenic mouse for in vivo detection of endogenous labeled mRNA. Nat. Methods 8, 165–170 (2011).

38. Chen, B.C. et al. Lattice light-sheet microscopy: Imaging molecules to embryos at high spatiotemporal resolution. Science 346, 1257998 (2014).

