## Supplementary Note 1 for "A series of spontaneously blinking dyes for super-resolution microscopy"

### A series of spontaneously blinking dyes for super-resolution microscopy

Katie L. Holland, Sarah E. Plutkis, Timothy A. Daugird, Abhishek Sau, Jonathan B. Grimm, Brian P. English, Qinsi Zheng, Sandeep Dave, Fariha Rahman, Liangqi Xie, Peng Dong, Ariana N. Tkachuk, Timothy A. Brown, Robert H. Singer, Catherine G. Galbraith, Siegfried M. Musser, Wesley R. Legant, and Luke D. Lavis*

*

#### Experimental Information: Synthetic Efforts towards HM-SiFl Ditriflate

| **Page** | **Contents** |
|---|---|
| S2 | Schemes S1–S4 and Table S1 |
| S4 | Summary: Development of a Concise Synthetic Route to HM-Xanthenes |
| S8 | General Experimental Information for Synthesis |
| S9–S19 | Experimentals and Characterization Data for All Compounds |
| S9 | Ketone and Aryl Iodide Intermediates |
| S13 | Xanthenes *via* Optimized Grignard Additions |
| S18 | HM-Si-Fluorescein Ditriflate **2** |
| S20 | References |

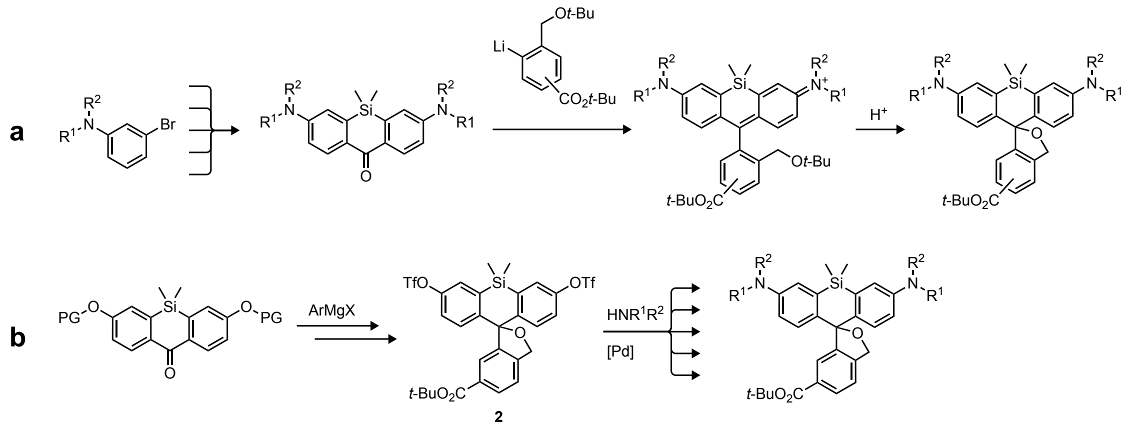


**Scheme S1.** Synthetic approaches to hydroxymethyl-Si-rhodamines (HM-SiR). (a) Existing strategy for HM-SiR synthesis via aryllithium addition to diamino-Si-xanthones. (b) Divergent C–N cross-coupling approach to HM-SiRs employing a common, late-stage ditriflate intermediate **2**.

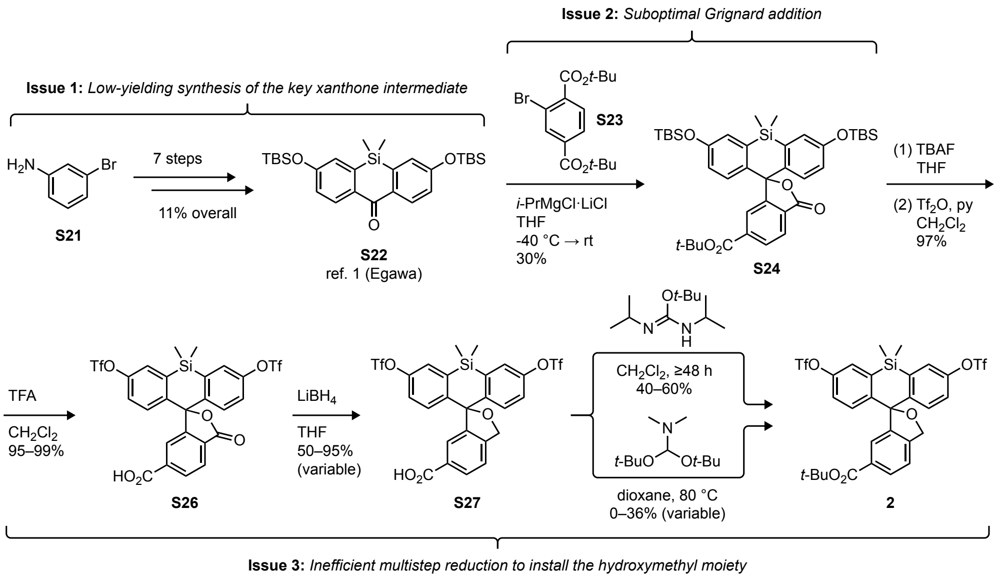


**Scheme S2.** First-generation synthesis of ditriflate intermediate **2**, annotated to highlight shortcomings targeted for optimization: (1) xanthone synthesis[1]; (2) Grignard addition; and (3) hydroxymethyl installation.

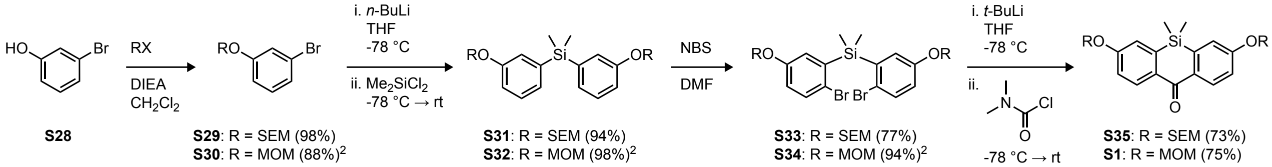


**Scheme S3.** Four-step synthesis of acetal-protected Si-xanthones **S1** and **S35** from 3-bromophenol (**S28**); synthesis of intermediates **S30**, **S32**, and **S34** from previous work.[2]

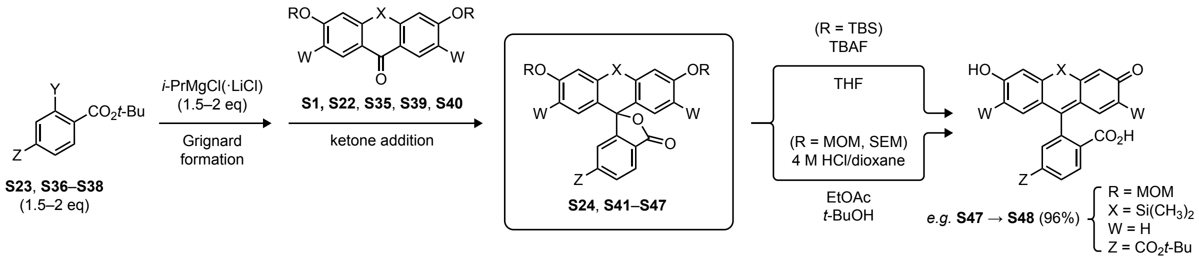


| entry | halide | Y | Z | ketone | X | R | W | Grignard formation | ketone addition | product | yield (%) | |
|---|---|---|---|---|---|---|---|---|---|---|---|---|
| 1 | **S36** | Br | H | **S39** | $C(CH_3)_2$ | TBS | H | *i*-PrMgCl·LiCl, -15 °C; -5 °C, 6 h | -5 °C; rt, 1 h | **S41** | 56 ref 3 | "Turbo Grignard" + ArBr |
| 2 | **S23** | Br | $CO_2$*t*-Bu | **S39** | $C(CH_3)_2$ | TBS | H | *i*-PrMgCl·LiCl, -15 °C; -10 °C, 4 h | -10 °C; rt, 2 h | **S42** | 17 ref 4 | |
| 3 | **S36** | Br | H | **S40** | $C(CH_3)_2$ | TBS | F | *i*-PrMgCl·LiCl, -15 °C; -5 °C, 6 h | -5 °C; rt, 0.5 h | **S43** | 50 ref 5 | |
| 4 | **S23** | Br | $CO_2$*t*-Bu | **S40** | $C(CH_3)_2$ | TBS | F | *i*-PrMgCl·LiCl, -50 °C; -40 °C, 4 h | -40 °C; rt, 2 h | **S44** | 31 ref 6 | |
| 5 | **S36** | Br | H | **S22** | $Si(CH_3)_2$ | TBS | H | *i*-PrMgCl·LiCl, -15 °C; -5 °C, 5 h | -5 °C; rt, 0.5 h | **S45** | 56 ref 7 | |
| 6 | **S23** | Br | $CO_2$*t*-Bu | **S22** | $Si(CH_3)_2$ | TBS | H | *i*-PrMgCl·LiCl, -50 °C; -40 °C, 2 h | -40 °C; rt, 2 h | **S24** | 30 ref 7 | |
| 7 | **S37** | I | H | **S39** | $C(CH_3)_2$ | TBS | H | *i*-PrMgCl, -40 °C; ↑ -25 °C, 2 h | -25 °C; rt, 1 h | **S41** | 72 | standard Grignard + ArI |
| 8 | **S38** | I | $CO_2$*t*-Bu | **S39** | $C(CH_3)_2$ | TBS | H | *i*-PrMgCl, -40 °C; ↑ -25 °C, 2 h | -25 °C; rt, 3 h | **S42** | 50 | |
| 9 | **S37** | I | H | **S40** | $C(CH_3)_2$ | TBS | F | *i*-PrMgCl, -40 °C; ↑ -25 °C, 2 h | -25 °C; rt, 0.5 h | **S43** | 80 | |
| 10 | **S38** | I | $CO_2$*t*-Bu | **S40** | $C(CH_3)_2$ | TBS | F | *i*-PrMgCl, -40 °C; ↑ -25 °C, 2 h | -25 °C; rt, 2 h | **S44** | 68 | |
| 11 | **S37** | I | H | **S22** | $Si(CH_3)_2$ | TBS | H | *i*-PrMgCl, -40 °C; ↑ -25 °C, 2 h | -25 °C; rt, 1 h | **S45** | 94 | |
| 12 | **S38** | I | $CO_2$*t*-Bu | **S22** | $Si(CH_3)_2$ | TBS | H | *i*-PrMgCl, -40 °C; ↑ -25 °C, 2 h | -25 °C; rt, 3 h | **S24** | 79 | |
| 13 | **S38** | I | $CO_2$*t*-Bu | **S35** | $Si(CH_3)_2$ | SEM | H | *i*-PrMgCl, -40 °C; ↑ -25 °C, 2 h | -25 °C; rt, 2 h | **S46** | 81 | |
| 14 | **S38** | I | $CO_2$*t*-Bu | **S1** | $Si(CH_3)_2$ | MOM | H | *i*-PrMgCl, -40 °C; ↑ -25 °C, 2 h | -25 °C; rt, 2 h | **S47** | 85 | |

**Table S1.** Optimization of RMgX addition to anthrones and Si-xanthones for the synthesis of carbo- and Si-fluoresceins. Entries 1–6 summarize our prior work[3-7] using the Turbo Grignard reagent with aryl bromides; entries 7–14 illustrate the substantial improvement seen in this report with standard *i*-PrMgCl and the analogous aryl iodides.

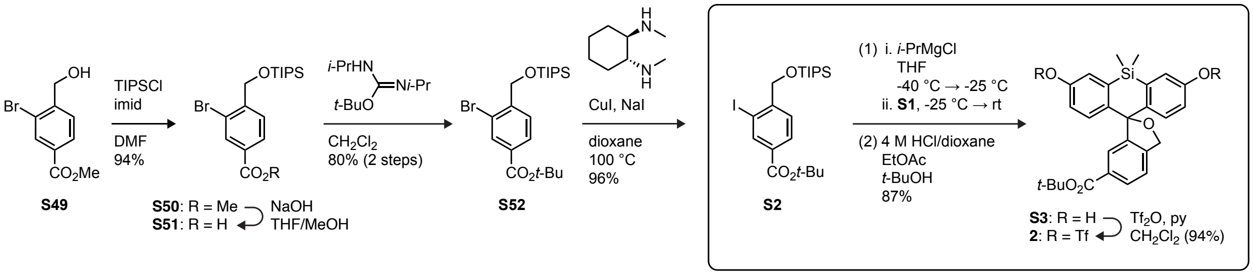


**Scheme S4.** Second-generation route to ditriflate **2** featuring direct installation of the hydroxymethyl functionality using the improved Grignard protocol with iodide **S2** and subsequent addition to the readily accessible ketone **S1**.

#### Summary: Development of a Concise Synthetic Route to HM-Xanthenes

Fine-tuning the blinking properties of the JFb scaffold was contingent on our ability to efficiently synthesize a variety of hydroxymethyl-Si-rhodamines (HM-SiRs) containing diversely substituted azetidine and pyrrolidine auxochromes. To that end, we sought a synthetic route that would enable incorporation of the diversity element—the 3′/6′ amine auxochrome substituent—at a late stage (**Scheme S1**). Unfortunately, the previously reported HM-SiR syntheses begin with a bromoaniline (**Scheme S1a**), such that the auxochrome (*i.e.*, the amine substituent $NR^1R^2$) is set during the first step of the synthesis.[8-11] The bromoaniline is typically converted into a diamino-Si-xanthone through a 3–4 step synthesis; subsequent installation of the pendant phenyl ring is generally achieved (often in modest yield) via addition of an aryllithium reagent bearing an *ortho*-hydroxymethyl substituent protected as a *tert*-butyl ether. Deprotection of the *tert*-butyl ether to give the hydroxymethyl dye requires strong acid and extended reaction time (*e.g.*, neat TFA, >24 h; **Scheme S1a**). This was also problematic for our purposes, as azetidinyl rhodamines—particularly those substituted on the azetidine ring with electron-withdrawing groups—are susceptible to ring-opening and other side reactions under harshly acidic conditions.

We instead pursued a more divergent synthetic strategy focusing on late-stage incorporation of the amine substituents through Pd-catalyzed C–N cross-coupling (**Scheme S1b**). In previous reports, we have extensively shown that the cross-coupling of fluorescein ditriflates with nitrogen nucleophiles is an efficient method for the synthesis of traditional lactone (*o*-carboxyl) rhodamines. Based on this—as well as preliminary work[12] describing the synthesis of a spontaneously blinking green dye (HM-$JF_{526}$)—we surmised that hydroxymethyl-Si-fluorescein (HM-SiFl) ditriflate **2** could serve as the common intermediate for cross-coupling with various amines, thereby allowing direct incorporation of the auxochrome at the latest possible stage. We expected that **2** would be most efficiently accessed by reaction of a protected dihydroxy-Si-xanthone with an appropriately functionalized aryl Grignard (**Scheme S1b**), which would better accommodate a wider range of protective functionality than an aryllithium reagent.

The route to ditriflate **2** underwent several optimizations, and the evolution of its synthesis is presented in **Schemes S2–S4** and **Table S1**. Our initial approach (**Scheme S2**) proceeded through the previously reported[7] Si-fluorescein ditriflate **S25**, which is itself prepared through addition of the Grignard derived from di-*tert*-butyl 2-bromoterephalate (**S23**) to the known[1] Si-xanthone **S22**. Consistent with our previous synthesis of an HM-rhodamine,[12] we found that the lactone could be selectively reduced (in modest and variable yield) to the cyclic ether with $LiBH_4$, but only by temporarily removing the *tert*-butyl protection on the 6-carboxyl group (**S25**→**S26**→**S27**). This then required reinstallation of the *tert*-butyl ester in the presence of the hydrolytically sensitive triflates to give the desired intermediate (**2**). Although this relatively long, modest-yielding synthesis was initially adequate for accessing small amounts of select JFb dyes, more substantial SAR exploration necessitated a better route. Hence, we optimized this route by addressing three distinct issues (**Schemes S3–S4**, **Table S1**) summarized below.

(1) **Low-yielding synthesis of the key xanthone intermediate (Scheme S3).** Si-xanthone **S22** was a crucial intermediate in our previous synthetic efforts towards azetidinyl-Si-rhodamines. To access this material, we made use of what was, at the time, the only published synthesis: Nagano's 2011 report of a 7-step route with an overall yield of

11% (**Scheme S2**).[1] Since that time, several shorter and more efficient syntheses have been reported.[13-16] Most of these rely on an approach similar to one first described by our laboratory[2] for the synthesis of Si-fluoresceins and -rhodamines; namely, the double lithium-bromide exchange of bis(2-bromophenyl)silanes and addition to a phosgene equivalent such as dimethylcarbamoyl chloride (**Scheme S3**). However, the use of silyl ether protecting groups on the phenols—particularly TBS—often results in diminished yields[15] or requires reprotection of the crude product following ketone installation[16] (*i.e.*, the lithium-bromide exchange and electrophile addition step).

With this in mind, we chose to protect the phenols with more robust acetal protecting groups. As shown in **Scheme S3**, the acetals (MOM and SEM) were installed in the first step through facile alkylation of 3-bromophenol with MOM-Br or SEM-Cl. Lithium-bromide exchange with *n*-butyllithium and addition to $Me_2SiCl_2$ afforded effectively quantitative yields (94–98%) of silanes **S31** and **S32**, which were regioselectively brominated with NBS/DMF to give the corresponding dibromides (**S33** and **S34**). Finally, lithium-bromide exchange with *tert*-butyllithium and addition to dimethylcarbamoyl chloride yielded the Si-xanthones **S35** and **S1** in excellent yield (73–75%). The choice of alkyllithium reagent was important, as *tert*-butyllithium gave substantially higher yields and cleaner reactions than *n*-butyllithium. This route to Si-xanthones represents a substantial improvement over the original synthesis[1] in both length (7 steps → 4 steps) and overall yield (SEM: 11% → 52%; MOM: 11% → 61%). Although the syntheses of both acetal-protected ketones were straightforward, we ultimately found $MOM_2$-Si-xanthone **S1** to be the more convenient choice due to easier chromatography, the smaller size of the protecting group, and the solid/crystalline states of the MOM dibromide **S34** and ketone **S1** (as compared to the SEM-protected viscous oils **S33** and **S35**).

(2) **Suboptimal Grignard addition (Table S1).** The key step in most existing syntheses of carbo- and Si-fluoresceins is the installation of the bottom (pendant) phenyl ring through addition of an arylmetal species to a protected dihydroxy anthrone/Si-xanthone. Although both aryllithium and aryl Grignard reagents will add to these ketones, the milder, less nucleophilic Grignards are safer and more tolerant of the *tert*-butyl esters commonly used to protect the necessary carboxyl substituents. In contrast, aryllithiums routinely require either (a) careful selection of alkyllithium reagent (*n*-/*sec*-/*tert*-butyllithium) with oftentimes extreme, inconvenient temperatures (-100 °C to -150 °C) to tolerate *t*-butyl protection,[14,17-19] or (b) more robust protecting groups (*e.g.*, oxazoline, orthoester) that can be challenging to remove.[2,18,20-22] (It is important to note that aryllithiums are typically *required* when using this approach for the synthesis of rhodamines, as most aryl Grignards are not reactive enough to add to diaminoanthrones or -xanthones.[22,23]) As a result, we have typically favored the use of Grignards for the synthesis of carbo- and Si-fluoresceins.[3-7] We previously accessed the necessary Grignard reagents via magnesium-bromide exchange of aryl bromides such as *tert*-butyl 2-bromobenzoate (**S36**) or di-*tert*-butyl 2-bromoterephthalate (**S23**) with Knochel's "Turbo Grignard" reagent *i*-PrMgCl·LiCl (**Scheme S2**, **Table S1**), as Mg/Br-exchange of aryl bromides with standard *i*-PrMgCl is often very slow.[24] We used these aryl Grignards with several dihydroxyanthrones and dihydroxy-Si-xanthones (**S22**, **S39**, **S40**) over the past decade to synthesize a number of carbofluoresceins[3,5,6] (**Table S1**, entries 1–4) and Si-fluoresceins[4,7] (**Table S1**, entries 5–6).

This method, however, presented several opportunities for improvement. Firstly, the yields obtained from the Mg/Br-exchange and ketone addition were quite variable (**Table S1**, entries 1-6), ranging from low (17%, entry 2) to moderate (56%, entries 1, 5). In our hands, the commercial Turbo Grignard reagent displayed limited and inconsistent shelf stability once opened (as compared to a standard Grignard or organolithium). Perhaps consequently, the Mg/Br-exchange rarely consumed the bromide in its entirety, regardless of temperature or extended reaction time. The addition step also seldom proceeded to completion, with significant ketone starting material recovered even when using a substantial excess of Grignard (>2 eq). In the case of the present work, the modest yield of Si-fluorescein **S24** obtained via the Turbo Grignard route (30%, entry 6) was a significant bottleneck in the synthesis of ditriflate **2** (**Scheme S2**).

Given these results, we considered whether switching to an aryl halide with more facile reactivity in the magnesium-halide exchange could significantly improve yield and conversion. It is well-established that aryl iodides undergo metal-halide exchange with Grignard reagents (*e.g, i*-PrMgCl) faster than aryl bromides; in fact, Mg/I-exchange of iodides in the *presence* of bromides is a reliable method to prepare bromoaryl Grignards from bromoiodoarenes.[25-27] Indeed, when we substituted *tert*-butyl iodobenzoate (**S37**) or di-*tert*-butyl iodoterephthalate (**S38**) for the corresponding bromides and used *i*-PrMgCl in place of *i*-PrMgCl·LiCl, we observed a dramatic improvement across all six examples (**Table S1**, entries 7–12). The Mg/I-exchanges were cleaner and faster (-40 °C → -25 °C, ≤ 2 h), negligible amounts of unreacted ketone were recovered, and the yields of the fluorescein products were greatly enhanced (up to 3× higher). Importantly, the yield of Si-fluorescein **S24**, the key intermediate in our initial route to ditriflate **2** (**Scheme S2**), was increased to 79% by this modification (entry 6 → entry 12). TBS-protected ketones were used in entries 7–12 to be most consistent with our previous work[3-7] (entries 1–6) and enable direct comparison of the efficiencies of the two Grignard protocols.

Given the improved Si-xanthone synthesis described in (1) above, we then applied the same method to the SEM- and MOM-protected Si-xanthones **S35** and **S1** (**Table S1**, entries 13–14). Both smoothly reacted with the Grignard reagent derived from di-*tert*-butyl 2-iodoterephthalate (**S38**) to afford even better yields of Si-fluoresceins **S46** (81%) and **S47** (85%). Since acetal (*e.g.*, MOM) deprotection is commonly achieved with acid, we were initially concerned that removal of the MOM group in the presence of the *tert*-butyl ester—which was required for subsequent ditriflate formation and, ultimately, the cross-coupling—might prove difficult. Gratifyingly, several conditions were found that could effectively remove the acetal without disturbing the ester, including *p*-TsOH in MeOH (24–48 h, 25 °C)[28,29] and 4 M HCl/dioxane in 2:1 EtOAc/*t*-BuOH (< 2 h, 25 °C).[30] For example, the faster HCl/dioxane conditions promoted rapid deprotection of $MOM_2$-Si-fluorescein **S47** to afford a nearly quantitative (96%) yield of 6-*tert*-butoxycarbonyl-Si-fluorescein **S48** (**Table S1**; see **Xanthenes via Optimized Grignard Addition**). Curiously, our efforts to effect a more orthogonal and chemospecific deprotection of $SEM_2$-Si-fluorescein **S46** with fluoride (*e.g.*, TBAF) were unsuccessful; nonetheless, the SEM acetals could be easily removed under the same acid-mediated conditions used for the MOM analog (see **S48** in **Xanthenes via Optimized Grignard Addition**).

(3) **Inefficient multistep reduction to install the hydroxymethyl moiety (Scheme S4).** With a dramatically improved method for installation of the bottom phenyl ring in hand, we then turned our focus to the final transformation in our initial synthesis of **2** (**Scheme S2**): the circuitous, 3-step reduction protocol used to convert Si-fluorescein **S25** to its hydroxymethyl (cyclic ether) congener **2**. In order the selectively reduce the lactone to the cyclic

ether, it was essential to remove the *tert*-butyl protecting group from the 6-carboxyl group. Unfortunately, this then required post-reduction reinstallation of the *tert*-butyl ester in the presence of two labile aryl triflates. Even the most mild reagents for carboxyl *tert*-butylation—most notably *tert*-butyl *N*,*N′*-diisopropylcarbamimidate[31] and *N*,*N*-dimethylformamide di-*tert*-butyl acetal[32]—required elevated temperature and/or extended reaction time to achieve reasonable conversion to **2**, and yields were variable and moderate at best (due in no small part to triflate hydrolysis, particularly with the DMF acetal). Moreover, the $LiBH_4$ reduction itself also exhibited variability in conversion and yield, sometimes providing ≤ 50% of the reduced intermediate **2**.

In light of these issues, we considered whether we might eschew the suboptimal reduction and protecting group shuffling entirely by directly installing the *o*-hydroxymethyl substituent during the Grignard addition (**Scheme S4**). To that end, we synthesized a candidate aryl iodide (**S2**, Scheme S4) through a simple 4-step route from the commercially available benzoate **S49**. Straightforward TIPS protection of the free hydroxymethyl group afforded a 94% yield of silyl ether **S50**; the methyl ester was converted to a *tert*-butyl ester via simple 2-step saponification and *tert*-butylation with *tert*-butyl *N*,*N′*-diisopropylcarbamimidate (**S52**; 80% yield, 2 steps). Lastly, halogen exchange of **S52** using Buchwald's Cu(I)-catalyzed aromatic Finkelstein reaction[33] cleanly provided the desired iodide intermediate **S2** in excellent yield (96%).

When subjected to the optimized Grignard conditions, iodide **S2** underwent facile magnesium-iodide exchange with *i*-PrMgCl and efficient addition to $MOM_2$-Si-xanthone **S1**. Rather than purify the initial addition product and remove the TIPS and MOM protecting groups in a stepwise fashion, we opted to directly subject the crude Grignard addition product to the MOM deprotection conditions discussed above (4 M HCl/dioxane in EtOAc/*t*-BuOH). Conveniently, all the protecting groups save the *tert*-butyl ester were cleanly removed to provide, following purification, an 87% yield of HM-Si-fluorescein **S3**. Finally, trivial installation of the triflates with trifluoromethanesulfonic anhydride afforded the long-sought HM-SiFl ditriflate **2** (94%). **Scheme S4**, then, represents our current and most optimized route to this key cross-coupling substrate (see **Supplementary Note 2**).

In comparing our original (**Scheme S2**) and final (**Scheme S4**) syntheses, we note that systematically addressing three major flaws of the initial route to ditriflate **2** has achieved both a dramatic improvement in efficiency (13 steps → 6 steps, longest linear sequence) and overall yield (< 2% → 50%). We expect that these new and modified synthetic approaches will be of significant interest and use to the many fluorophore chemists investigating not only HM-xanthenes specifically, but also the ever-widening variety of xanthene dyes in general.

#### General Experimental Information for Synthesis

Commercial reagents were obtained from reputable suppliers and used as received. All solvents were purchased in septum-sealed bottles stored under an inert atmosphere. All reactions were sealed with septa through which a nitrogen atmosphere was introduced unless otherwise noted. Reactions were conducted in round-bottomed flasks or septum-capped crimp-top vials containing Teflon-coated magnetic stir bars. Heating of reactions was accomplished with a silicon oil bath or an aluminum reaction block on top of a stirring hotplate equipped with an electronic contact thermometer to maintain the indicated temperatures.

Reactions were monitored by thin layer chromatography (TLC) on precoated TLC glass plates (silica gel 60 $F_{254}$, 250 μm thickness) or by LC/MS (Phenomenex Kinetex 2.1 mm × 30 mm 2.6 μm C18 column; 5 μL injection; 5–98% $MeCN/H_2O$, linear gradient, with constant 0.1% v/v $HCO_2H$ additive; 6 min run; 0.5 mL/min flow; ESI; positive ion mode). TLC chromatograms were visualized by UV illumination or developed with *p*-anisaldehyde, ceric ammonium molybdate, or $KMnO_4$ stain. Reaction products were purified by flash chromatography on an automated purification system using pre-packed silica gel columns or by preparative HPLC (Phenomenex Gemini–NX 30 × 150 mm 5 μm C18 column). Analytical HPLC analysis was performed with an Agilent Eclipse XDB 4.6 × 150 mm 5 μm C18 column under the indicated conditions. High-resolution mass spectrometry was performed by the High Resolution Mass Spectrometry Facility at the University of Iowa.

NMR spectra were recorded on a 400 MHz spectrometer. $^{1}H$ and $^{13}C$ chemical shifts were referenced to TMS or residual solvent peaks, and $^{19}F$ chemical shifts were referenced to $CFCl_3$. Data for $^{1}H$ NMR spectra are reported as follows: chemical shift (δ ppm), multiplicity (s = singlet, d = doublet, t = triplet, q = quartet, dd = doublet of doublets, m = multiplet), coupling constant (Hz), integration. Data for $^{13}C$ NMR spectra are reported by chemical shift (δ ppm) with hydrogen multiplicity (C, CH, $CH_2$, $CH_3$) information obtained from DEPT spectra.

#### Ketone and Aryl Iodide Intermediates

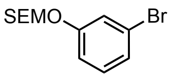


**(2-((3-Bromophenoxy)methoxy)ethyl)trimethylsilane (S29):** 3-Bromophenol (**S28**; 7.50 g, 43.35 mmol) was taken up in $CH_2Cl_2$ (100 mL); DIEA (11.33 mL, 65.03 mmol, 1.5 eq) and 2-(trimethylsilyl)ethoxymethyl chloride (11.51 mL, 65.03 mmol, 1.5 eq) were successively added, and the reaction was stirred at room temperature for 2 h. It was subsequently diluted with water and extracted with $CH_2Cl_2$ (2×). The combined organic extracts were washed with saturated $NaHCO_3$ and brine, dried over anhydrous $MgSO_4$, filtered, and evaporated. Flash chromatography (0–20% $Et_2O$/hexanes, linear gradient) afforded **S29** as a colorless oil (12.92 g, 98%). $^1H$ NMR ($CDCl_3$, 400 MHz) δ 7.24 – 7.20 (m, 1H), 7.17 – 7.10 (m, 2H), 7.02 – 6.91 (m, 1H), 5.20 (s, 2H), 3.78 – 3.71 (m, 2H), 0.99 – 0.92 (m, 2H), 0.00 (s, 9H); $^{13}C$ NMR ($CDCl_3$, 101 MHz) δ 158.4 (C), 130.7 (CH), 124.9 (CH), 122.8 (C), 119.7 (CH), 115.2 (CH), 93.1 ($CH_2$), 66.6 ($CH_2$), 18.2 ($CH_2$), -1.3 ($CH_3$); HRMS (EI) calcd for $C_{12}H_{19}BrO_2Si$ $[M]^{+\bullet}$ 302.0333/304.0312, found 302.0332/304.0311.

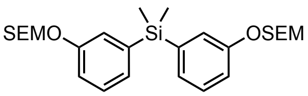


**Dimethylbis(3-((2-(trimethylsilyl)ethoxy)methoxy)phenyl)silane (S31):** A solution of (2-((3-bromophenoxy)methoxy)ethyl)trimethylsilane (**S29**; 12.50 g, 41.22 mmol, 2.4 eq) in THF (100 mL) was cooled to -78 °C under nitrogen. *n*-Butyllithium (1.6 M in hexanes, 25.76 mL, 41.22 mmol, 2.4 eq) was added, and the reaction was stirred at -78 °C for 30 min. Dichlorodimethylsilane (2.07 mL, 17.17 mmol) was then added. The dry ice bath was removed, and the reaction was stirred at room temperature for 2 h. It was subsequently quenched with saturated $NH_4Cl$, diluted with water, and extracted with EtOAc (2×). The combined organic extracts were washed with brine, dried over anhydrous $MgSO_4$, filtered, and concentrated *in vacuo*. Purification by flash chromatography on silica gel (0–10% $Et_2O$/hexanes, linear gradient) afforded 8.12 g (94%) of **S31** as a colorless oil. $^1H$ NMR ($CDCl_3$, 400 MHz) δ 7.27 (ddd, $J$ = 8.2, 7.2, 0.5 Hz, 2H), 7.18 (ddd, $J$ = 2.7, 1.1, 0.4 Hz, 2H), 7.14 (dt, $J$ = 7.2, 1.1 Hz, 2H), 7.06 (ddd, $J$ = 8.2, 2.6, 1.1 Hz, 2H), 5.20 (s, 4H), 3.78 – 3.72 (m, 4H), 0.98 – 0.92 (m, 4H), 0.53 (s, 6H), 0.00 (s, 18H); $^{13}C$ NMR ($CDCl_3$, 101 MHz) δ 157.2 (C), 140.0 (C), 129.2 (CH), 127.7 (CH), 122.3 (CH), 116.8 (CH), 93.2 ($CH_2$), 66.3 ($CH_2$), 18.2 ($CH_2$), -1.3 ($CH_3$), -2.3 ($CH_3$); HRMS (ESI) calcd for $C_{26}H_{44}O_4Si_3Na$ $[M+Na]^+$ 527.2440, found 527.2438.

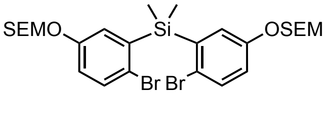


**Bis(2-bromo-5-((2-(trimethylsilyl)ethoxy)methoxy)phenyl)dimethylsilane (S33):** Dimethylbis(3-((2-(trimethylsilyl)ethoxy)methoxy)phenyl)silane (**S31**; 8.00 g, 15.85 mmol) was taken up in DMF (100 mL). *N*-Bromosuccinimide (5.64 g, 31.69 mmol, 2 eq) was added portion-wise over 5 min, and the reaction was then stirred at room temperature for 96 h. The reaction mixture was concentrated *in vacuo*; the resulting residue was diluted with water and extracted with EtOAc (2×). The combined organic extracts were washed with water and brine, dried over anhydrous $MgSO_4$, filtered, and concentrated *in vacuo*. Silica gel chromatography (0–10% $Et_2O$/hexanes, linear gradient) afforded 8.09 g (77%) of **S33** as a pale yellow oil. $^1H$ NMR ($CDCl_3$, 400 MHz) δ 7.40 (d, $J$ = 8.7 Hz, 2H), 7.14 (d, $J$ = 3.0 Hz, 2H), 6.95 (dd, $J$ = 8.7, 3.1 Hz, 2H), 5.17 (s, 4H), 3.76 – 3.70 (m, 4H), 0.97 – 0.90 (m, 4H), 0.73 (s, 6H), 0.00 (s, 18H); $^{13}C$ NMR ($CDCl_3$, 101 MHz) δ 156.3 (C), 140.2 (C), 133.8 (CH), 125.8 (CH), 121.9 (C), 118.8 (CH), 93.3 ($CH_2$), 66.4 ($CH_2$), 18.2 ($CH_2$), -1.1 ($CH_3$), -1.2 ($CH_3$); HRMS (ESI) calcd for $C_{26}H_{42}Br_2O_4Si_3Na$ $[M+Na]^+$ 683.0650/685.0630/687.0609, found 683.0646/685.0625/687.0607.

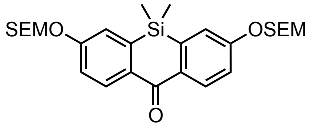


**5,5-Dimethyl-3,7-bis((2-(trimethylsilyl)ethoxy)methoxy)dibenzo[*b*,*e*]silin-10(5*H*)-one (S35):** A solution of bis(2-bromo-5-((2-(trimethylsilyl)ethoxy)methoxy)phenyl)dimethylsilane (**S33**; 5.50 g, 8.30 mmol) in THF (80 mL) was cooled to -78 °C under nitrogen. *tert*-Butyllithium (1.7 M in pentane, 21.48 mL, 36.52 mmol, 4.4 eq) was added, and the reaction was stirred at -78 °C for 30 min. Dimethylcarbamoyl chloride (841 µL, 9.13 mmol, 1.1 eq) was added; the reaction was then gradually warmed to room temperature over 4 h. It was subsequently quenched with saturated $NH_4Cl$, diluted with water, and extracted with EtOAc (2×). The combined organic extracts were washed with brine, dried over anhydrous $MgSO_4$, filtered, and concentrated *in vacuo*. Purification by flash chromatography on silica gel (0–20% $Et_2O$/hexanes, linear gradient) yielded 3.21 g (73%) of **S35** as a colorless gum. $^1H$ NMR ($CDCl_3$, 400 MHz) δ 8.42 (d, $J$ = 8.8 Hz, 2H), 7.23 (d, $J$ = 2.5 Hz, 2H), 7.20 (dd, $J$ = 8.8, 2.6 Hz, 2H), 5.31 (s, 4H), 3.82 – 3.75 (m, 4H), 1.00 – 0.93 (m, 4H), 0.47 (s, 6H), 0.00 (s, 18H); $^{13}C$ NMR ($CDCl_3$, 101 MHz) δ 186.0 (C), 160.1 (C), 141.4 (C), 134.9 (C), 132.4 (CH), 119.7 (CH), 117.7 (CH), 92.8 ($CH_2$), 66.8 ($CH_2$), 18.2 ($CH_2$), -1.2 ($CH_3$), -1.3 ($CH_3$); HRMS (ESI) calcd for $C_{27}H_{43}O_5Si_3$ $[M+H]^+$ 531.2413, found 531.2412.

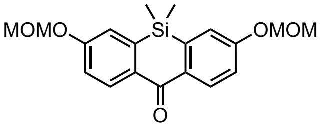


**3,7-Bis(methoxymethoxy)-5,5-dimethyldibenzo[*b*,*e*]silin-10(5*H*)-one (S1):** A solution of bis(2-bromo-5-(methoxymethoxy)phenyl)dimethylsilane[2] (**S34**; 6.00 g, 12.24 mmol) in THF (100 mL) was cooled to -78 °C under nitrogen. *tert*-Butyllithium (1.7 M in pentane, 31.68 mL, 53.85 mmol, 4.4 eq) was added, and the reaction was stirred at -78 °C for 30 min. Dimethylcarbamoyl chloride (1.24 mL, 13.46 mmol, 1.1 eq) was added; the reaction was then gradually warmed to room temperature over 4 h. It was subsequently quenched with saturated $NH_4Cl$, diluted with water, and extracted with EtOAc (2×). The combined organic extracts were washed with brine, dried over anhydrous $MgSO_4$, filtered, and concentrated *in vacuo*. Purification by flash chromatography on silica gel (0–40% EtOAc/hexanes, linear gradient) yielded 3.31 g (75%) of **S1** as a white solid. $^1H$ NMR ($CDCl_3$, 400 MHz) δ 8.43 (d, $J$ = 8.7 Hz, 2H), 7.23 (d, $J$ = 2.6 Hz, 2H), 7.20 (dd, $J$ = 8.7, 2.7 Hz, 2H), 5.28 (s, 4H), 3.52 (s, 6H), 0.48 (s, 6H); $^{13}C$ NMR ($CDCl_3$, 101 MHz) δ 185.9 (C), 159.8 (C), 141.4 (C), 135.0 (C), 132.4 (CH), 119.8 (CH), 117.6 (CH), 94.3 ($CH_2$), 56.5 ($CH_3$), -1.3 ($CH_3$); HRMS (ESI) calcd for $C_{19}H_{23}O_5Si$ $[M+H]^+$ 359.1309, found 359.1308.

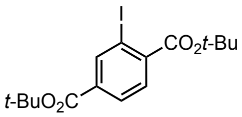

**Di-*tert*-butyl 2-iodoterephthalate (S38):** A solution of 2-iodoterephthalic acid (12.00 g, 41.1 mmol) in dioxane (80 mL) was heated to 80 °C, and *N*,*N*-dimethylformamide di-*tert*-butyl acetal (78.82 mL, 328.7 mmol, 8 eq) was added dropwise over 15 min. The reaction was stirred at 80 °C for an additional 1 h. After cooling the solution to room temperature, it was diluted with saturated $NaHCO_3$ and extracted with EtOAc (2×). The combined organic extracts were washed with water and brine, dried over anhydrous $MgSO_4$, filtered, and evaporated. Flash chromatography (0–10% $Et_2O$/hexanes, linear gradient) provided **S38** as a pale yellow, viscous oil (15.80 g, 95%). $^1$H NMR ($CDCl_3$, 400 MHz) δ 8.49 (d, *J* = 1.6 Hz, 1H), 7.96 (dd, *J* = 8.0, 1.6 Hz, 1H), 7.66 (d, *J* = 8.0 Hz, 1H), 1.63 (s, 9H), 1.59 (s, 9H); $^{13}$C NMR ($CDCl_3$, 101 MHz) δ 166.0 (C), 163.7 (C), 141.7 (CH), 141.1 (C), 134.9 (C), 130.0 (CH), 128.9 (CH), 92.7 (C), 83.4 (C), 82.3 (C), 28.3 ($CH_3$); HRMS (ESI) calcd for $C_{16}H_{21}IO_4Na$ $[M+Na]^+$ 427.0377, found 427.0381.

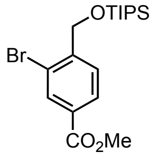

**Methyl 3-bromo-4-(((triisopropylsilyl)oxy)methyl)benzoate (S50):** To a solution of methyl 3-bromo-4-(hydroxymethyl)benzoate (**S49**; 7.95 g, 32.44 mmol) in DMF (100 mL) were added imidazole (4.42 g, 64.88 mmol, 2 eq) and TIPSCl (10.41 mL, 48.66 mmol, 1.5 eq). The reaction was stirred at room temperature for 4 h. It was subsequently diluted with water and extracted with $Et_2O$ (2×). The combined organic extracts were washed with water and brine, dried over anhydrous $MgSO_4$, filtered, and concentrated *in vacuo*. Silica gel chromatography (0–10% $Et_2O$/hexanes, linear gradient) afforded 12.19 g (94%) of **S50** as a colorless solid. $^1$H NMR ($CDCl_3$, 400 MHz) δ 8.17 (d, *J* = 1.6 Hz, 1H), 8.02 (dd, *J* = 8.1, 1.7 Hz, 1H), 7.72 (dt, *J* = 8.1, 1.1 Hz, 1H), 4.84 (d, *J* = 1.0 Hz, 2H), 3.92 (s, 3H), 1.27 – 1.17 (m, 3H), 1.11 (d, *J* = 7.0 Hz, 18H); $^{13}$C NMR ($CDCl_3$, 101 MHz) δ 166.1 (C), 145.9 (C), 133.2 (CH), 130.2 (C), 128.7 (CH), 127.3 (CH), 120.6 (C), 65.0 ($CH_2$), 52.4 ($CH_3$), 18.2 ($CH_3$), 12.1 (CH); HRMS (ESI) calcd for $C_{18}H_{30}BrO_3Si$ $[M+H]^+$ 401.1142/403.1122, found 401.1139/403.1117.

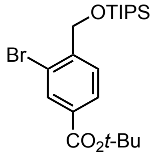

***tert*-Butyl 3-bromo-4-(((triisopropylsilyl)oxy)methyl)benzoate (S52):** Methyl 3-bromo-4-(((triisopropylsilyl)oxy)methyl)benzoate (**S50**; 11.90 g, 29.64 mmol) was dissolved in 3:1 THF/MeOH (160 mL), and 1 M NaOH (44.47 mL, 44.47 mmol, 1.5 eq) was added. The reaction was stirred at room temperature for 4 h. It was then acidified with 1 M HCl (50 mL), diluted with water, and extracted with EtOAc (2×). The combined organics were washed with brine, dried over anhydrous $MgSO_4$, filtered, and concentrated *in vacuo*. The crude carboxylic acid was taken back up in $CH_2Cl_2$ (300 mL) under nitrogen, and *tert*-butyl *N*,*N*'-diisopropylcarbamimidate (29.69 g, 148.2 mmol, 5 eq) was added. After stirring the reaction at room temperature for 18 h, the resulting white suspension was diluted with hexanes (300 mL) and concentrated to half-volume. The mixture was filtered through Celite with hexanes and concentrated *in vacuo*. Purification by flash chromatography (0–50% $CH_2Cl_2$/hexanes, linear gradient) yielded 10.48 g (80%) of **S52** as a colorless oil. $^1H$ NMR ($CDCl_3$, 400 MHz) δ 8.10 (d, *J* = 1.6 Hz, 1H), 7.96 (dd, *J* = 8.0, 1.7 Hz, 1H), 7.70 (dt, *J* = 8.1, 1.1 Hz, 1H), 4.84 (d, *J* = 1.1 Hz, 2H), 1.59 (s, 9H), 1.26 – 1.17 (m, 3H), 1.10 (d, *J* = 6.9 Hz, 18H); $^{13}C$ NMR ($CDCl_3$, 101 MHz) δ 164.7 (C), 145.3 (C), 133.0 (CH), 132.1 (C), 128.5 (CH), 127.1 (CH), 120.4 (C), 81.6 (C), 65.0 ($CH_2$), 28.3 ($CH_3$), 18.2 ($CH_3$), 12.1 (CH); HRMS (ESI) calcd for $C_{21}H_{36}BrO_3Si$ $[M+H]^+$ 443.1612/445.1592, found 443.1606/445.1584.

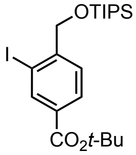


***tert*-Butyl 3-iodo-4-(((triisopropylsilyl)oxy)methyl)benzoate (S2):** An oven-dried round-bottom flask was charged with CuI (129 mg, 0.676 mmol, 0.1 eq) and NaI (2.03 g, 13.53 mmol, 2 eq). The flask was sealed and evacuated/backfilled with nitrogen (3×). A solution of *tert*-butyl 3-bromo-4-(((triisopropylsilyl)oxy)methyl)benzoate (**S52**; 3.00 g, 6.76 mmol) in dioxane (17 mL) was added, and the reaction was flushed again with nitrogen (3×). Following the addition of *trans*-*N*,*N*'-dimethylcyclohexane-1,2-diamine (213 µL, 1.35 mmol, 0.2 eq), the reaction was stirred at 110 °C for 24 h. It was then cooled to room temperature, diluted with saturated $NH_4Cl$ and water, and extracted with EtOAc (2×). The combined organic extracts were washed with brine, dried over anhydrous $MgSO_4$, filtered, and concentrated *in vacuo*. Silica gel chromatography (0–50% $CH_2Cl_2$/hexanes, linear gradient) afforded **S2** (3.19 g, 96%) as a pale yellow gum. $^1H$ NMR ($CDCl_3$, 400 MHz) δ 8.36 (d, *J* = 1.7 Hz, 1H), 8.00 (dd, *J* = 8.1, 1.7 Hz, 1H), 7.64 (dt, *J* = 8.1, 1.1 Hz, 1H), 4.72 (d, *J* = 1.0 Hz, 2H), 1.59 (s, 9H), 1.26 – 1.17 (m, 3H), 1.11 (d, *J* = 7.0 Hz, 18H); $^{13}C$ NMR ($CDCl_3$, 101 MHz) δ 164.5 (C), 147.8 (C), 139.6 (CH), 132.2 (C), 129.3 (CH), 126.8 (CH), 94.7 (C), 81.6 (C), 69.9 ($CH_2$), 28.3 ($CH_3$), 18.2 ($CH_3$), 12.1 (CH); HRMS (ESI) calcd for $C_{21}H_{36}IO_3Si$ $[M+H]^+$ 491.1473, found 491.1470.

#### XANTHENES VIA OPTIMIZED GRIGNARD ADDITIONS

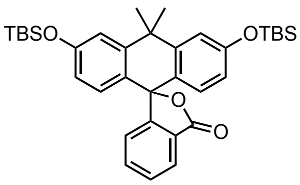


**Carbofluorescein bis(*tert*-butyldimethylsilyl ether) (S41):** To a solution of *tert*-butyl 2-iodobenzoate (**S37**; 1.65 g, 5.44 mmol, 1.5 eq) in THF (15 mL) at -40 °C was added *i*-PrMgCl (2.0 M in THF; 2.72 mL, 5.44 mmol, 1.5 eq). The reaction was then gradually warmed to -25 °C over 2 h while stirring. A solution of 3,6-bis((*tert*-butyldimethylsilyl)oxy)-10,10-dimethylanthracen-9(10*H*)-one[3] (**S39**; 1.75 g, 3.62 mmol) in THF (10 mL) was added dropwise; the reaction was then warmed to room temperature and stirred for 1 h. It was subsequently quenched with saturated $NH_4Cl$, diluted with water, and extracted with EtOAc (2×). The combined organic extracts were washed with brine, dried over anhydrous $MgSO_4$, filtered, and concentrated *in vacuo*. Flash chromatography on silica gel (0–20% $Et_2O$/hexanes, linear gradient) afforded 1.53 g (72%) of **S41** as a white solid. The characterization data for **S41** matched the previously reported spectra.[3]

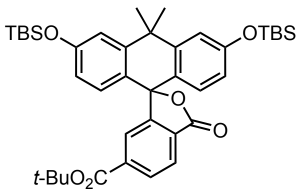


**6-(*tert*-Butoxycarbonyl)carbofluorescein bis(*tert*-butyldimethylsilyl ether) (S42):** To a solution of di-*tert*-butyl 2-iodoterephthalate (**S38**; 2.26 g, 5.59 mmol, 1.5 eq) in THF (15 mL) at -40 °C was added *i*-PrMgCl (2.0 M in THF; 2.80 mL, 5.59 mmol, 1.5 eq). The reaction was then gradually warmed to -25 °C over 2 h while stirring. A solution of 3,7-bis((*tert*-butyldimethylsilyl)oxy)-5,5-dimethyldibenzo[*b*,*e*]silin-10(5*H*)-one[3] (**S39**; 1.80 g, 3.73 mmol) in THF (10 mL) was added dropwise; the reaction was then warmed to room temperature and stirred for 3 h. It was subsequently quenched with saturated $NH_4Cl$, diluted with water, and extracted with EtOAc (2×). The combined organic extracts were washed with brine, dried over anhydrous $MgSO_4$, filtered, and concentrated *in vacuo*. Flash chromatography on silica gel (0–10% $Et_2O$/hexanes, linear gradient) afforded 1.28 g (50%) of **S42** as a white solid. The characterization data for **S42** matched the previously reported spectra.[4]

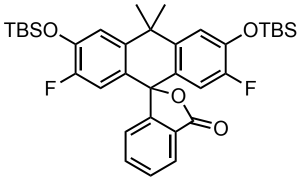


**Virginia Orange bis(*tert*-butyldimethylsilyl ether) (S43):** To a solution of *tert*-butyl 2-iodobenzoate (**S37**; 440 mg, 1.45 mmol, 1.5 eq) in THF (4 mL) at -40 °C was added *i*-PrMgCl (2.0 M in THF; 723 µL, 1.45 mmol, 1.5 eq). The reaction was then gradually warmed to -25 °C over 2 h while stirring. A solution of 3,6-bis((*tert*-

butyldimethylsilyl)oxy)-2,7-difluoro-10,10-dimethylanthracen-9(10*H*)-one[5] (**S40**; 500 mg, 0.964 mmol) in THF (2.5 mL) was added dropwise; the reaction was then warmed to room temperature and stirred for 30 min. It was subsequently quenched with saturated $NH_4Cl$, diluted with water, and extracted with EtOAc (2×). The combined organic extracts were washed with brine, dried over anhydrous $MgSO_4$, filtered, and concentrated *in vacuo*. Flash chromatography on silica gel (0–20% $Et_2O$/hexanes, linear gradient) afforded 479 mg (80%) of **S43** as a white foam. The characterization data for **S43** matched the previously reported spectra.[5]

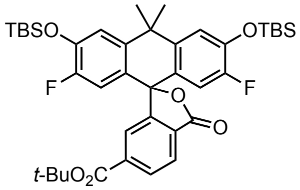


**6-(*tert*-Butoxycarbonyl)-Virginia Orange bis(*tert*-butyldimethylsilyl ether) (S44):** To a solution of di-*tert*-butyl 2-iodoterephthalate (**S38**; 2.34 g, 5.78 mmol, 1.5 eq) in THF (15 mL) at -40 °C was added *i*-PrMgCl (2.0 M in THF; 2.89 mL, 5.78 mmol, 1.5 eq). The reaction was then gradually warmed to -25 °C over 2 h while stirring. A solution of 3,6-bis((*tert*-butyldimethylsilyl)oxy)-2,7-difluoro-10,10-dimethylanthracen-9(10*H*)-one[5] (**S40**; 2.00 g, 3.86 mmol) in THF (10 mL) was added dropwise; the reaction was then warmed to room temperature and stirred for 2 h. It was subsequently quenched with saturated $NH_4Cl$, diluted with water, and extracted with EtOAc (2×). The combined organic extracts were washed with brine, dried over anhydrous $MgSO_4$, filtered, and concentrated *in vacuo*. Flash chromatography on silica gel (0–20% $Et_2O$/hexanes, linear gradient) afforded 1.90 g (68%) of **S44** as a white solid. The characterization data for **S44** matched the previously reported spectra.[6]

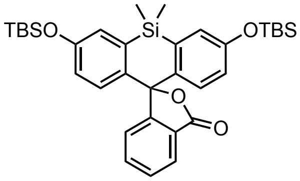


**Si-Fluorescein bis(*tert*-butyldimethylsilyl ether) (S45):** To a solution of *tert*-butyl 2-iodobenzoate (**S37**; 457 mg, 1.50 mmol, 1.5 eq) in THF (4 mL) at -40 °C was added *i*-PrMgCl (2.0 M in THF; 752 μL, 1.50 mmol, 1.5 eq). The reaction was then gradually warmed to -25 °C over 2 h while stirring. A solution of 3,7-bis((*tert*-butyldimethylsilyl)oxy)-5,5-dimethyldibenzo[*b*,*e*]silin-10(5*H*)-one[1] (**S22**; 500 mg, 1.00 mmol) in THF (2.5 mL) was added dropwise; the reaction was then warmed to room temperature and stirred for 1 h. It was subsequently quenched with saturated $NH_4Cl$, diluted with water, and extracted with EtOAc (2×). The combined organic extracts were washed with brine, dried over anhydrous $MgSO_4$, filtered, and concentrated *in vacuo*. Flash chromatography on silica gel (0–20% $Et_2O$/hexanes, linear gradient) afforded 566 mg (94%) of **S45** as a white solid. The characterization data for **S45** matched the previously reported spectra.[7]

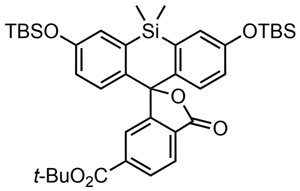

**6-(*tert*-Butoxycarbonyl)-Si-fluorescein bis(*tert*-butyldimethylsilyl ether) (S24):** To a solution of di-*tert*-butyl 2-iodoterephthalate (**S38**; 2.43 g, 6.01 mmol, 1.5 eq) in THF (15 mL) at -40 °C was added *i*-PrMgCl (2.0 M in THF; 3.01 mL, 6.01 mmol, 1.5 eq). The reaction was then gradually warmed to -25 °C over 2 h while stirring. A solution of 3,7-bis((*tert*-butyldimethylsilyl)oxy)-5,5-dimethyldibenzo[*b*,*e*]silin-10(5*H*)-one[1] (**S22**; 2.00 g, 4.01 mmol) in THF (10 mL) was added dropwise; the reaction was then warmed to room temperature and stirred for 3 h. It was subsequently quenched with saturated $NH_4Cl$, diluted with water, and extracted with EtOAc (2×). The combined organic extracts were washed with brine, dried over anhydrous $MgSO_4$, filtered, and concentrated *in vacuo*. Flash chromatography on silica gel (0–10% $Et_2O$/hexanes, linear gradient) afforded 2.22 g (79%) of **S24** as a white solid. The characterization data for **S24** matched the previously reported spectra.[7]

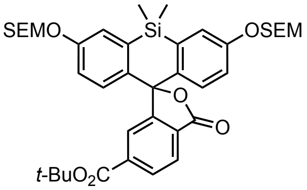

**6-(*tert*-Butoxycarbonyl)-Si-fluorescein bis(trimethylsilylethoxymethyl ether) (S46):** To a solution of di-*tert*-butyl 2-iodoterephthalate (**S38**; 3.05 g, 7.53 mmol, 2 eq) in THF (15 mL) at -40 °C was added *i*-PrMgCl (2.0 M in THF; 3.77 mL, 7.53 mmol, 2 eq). The reaction was then gradually warmed to -25 °C over 2 h while stirring. A solution of 5,5-dimethyl-3,7-bis((2-(trimethylsilyl)ethoxy)methoxy)dibenzo[*b*,*e*]silin-10(5*H*)-one (**S35**; 2.00 g, 3.77 mmol) in THF (10 mL) was added dropwise; the reaction was then warmed to room temperature and stirred for 2 h. It was subsequently quenched with saturated $NH_4Cl$, diluted with water, and extracted with EtOAc (2×). The combined organic extracts were washed with brine, dried over anhydrous $MgSO_4$, filtered, and concentrated *in vacuo*. Flash chromatography on silica gel (0–50% $Et_2O$/hexanes, linear gradient) afforded 2.24 g (81%) of **S46** as a white foam. $^1$H NMR ($CDCl_3$, 400 MHz) δ 8.14 (dd, $J$ = 8.0, 1.3 Hz, 1H), 7.99 (dd, $J$ = 8.0, 0.8 Hz, 1H), 7.85 (dd, $J$ = 1.3, 0.8 Hz, 1H), 7.33 (d, $J$ = 2.6 Hz, 2H), 7.00 (d, $J$ = 8.8 Hz, 2H), 6.95 (dd, $J$ = 8.9, 2.6 Hz, 2H), 5.23 (AB quartet, $\nu_A$ = 2092.3 Hz, $\nu_B$ = 2089.1 Hz, $J_{AB}$ = 7.0 Hz, 4H), 3.79 – 3.70 (m, 4H), 1.56 (s, 9H), 0.97 – 0.89 (m, 4H), 0.70 (s, 3H), 0.61 (s, 3H), -0.02 (s, 18H); $^{13}$C NMR ($CDCl_3$, 101 MHz) δ 170.0 (C), 164.3 (C), 157.0 (C), 155.0 (C), 137.5 (C), 137.2 (C), 136.9 (C), 130.3 (CH), 128.6 (C), 128.2 (CH), 126.0 (CH), 125.0 (CH), 121.4 (CH), 117.6 (CH), 93.0 ($CH_2$), 90.4 (C), 82.6 (C), 66.5 ($CH_2$), 28.2 ($CH_3$), 18.2 ($CH_2$), 0.0 ($CH_3$), -0.7 ($CH_3$), -1.3 ($CH_3$); HRMS (ESI) calcd for $C_{39}H_{55}O_8Si_3$ $[M+H]^+$ 735.3199, found 735.3188.

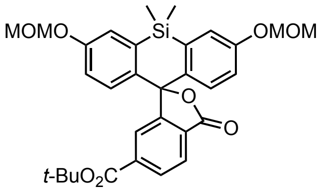


**6-(*tert*-Butoxycarbonyl)-Si-fluorescein bis(methoxymethyl ether) (S47):** To a solution of di-*tert*-butyl 2-iodoterephthalate (**S38**; 1.35 g, 3.35 mmol, 2 eq) in THF (8 mL) at -40 °C was added *i*-PrMgCl (2.0 M in THF; 1.67 mL, 3.35 mmol, 2 eq). The reaction was then gradually warmed to -25 °C over 2 h while stirring. A solution of 3,7-bis(methoxymethoxy)-5,5-dimethyldibenzo[*b*,*e*]silin-10(5*H*)-one (**S1**; 600 mg, 1.67 mmol) in THF (4 mL) was added dropwise; the reaction was then warmed to room temperature and stirred for 2 h. It was subsequently quenched with saturated $NH_4Cl$, diluted with water, and extracted with EtOAc (2×). The combined organic extracts were washed with brine, dried over anhydrous $MgSO_4$, filtered, and concentrated *in vacuo*. Flash chromatography on silica gel (0–70% $Et_2O$/hexanes, linear gradient) afforded 801 mg (85%) of **S47** as a white foam. $^1$H NMR ($CDCl_3$, 400 MHz) δ 8.13 (dd, $J$ = 8.0, 1.3 Hz, 1H), 7.98 (dd, $J$ = 8.0, 0.7 Hz, 1H), 7.85 (dd, $J$ = 1.3, 0.8 Hz, 1H), 7.32 (d, $J$ = 2.6 Hz, 2H), 7.00 (d, $J$ = 8.8 Hz, 2H), 6.94 (dd, $J$ = 8.9, 2.7 Hz, 2H), 5.18 (AB quartet, $\nu_A$ = 2076.4 Hz, $\nu_B$ = 2069.2 Hz, $J_{AB}$ = 6.9 Hz, 4H), 3.47 (s, 6H), 1.56 (s, 9H), 0.70 (s, 3H), 0.62 (s, 3H); $^{13}$C NMR ($CDCl_3$, 101 MHz) δ 170.0 (C), 164.3 (C), 156.8 (C), 154.9 (C), 137.5 (C), 137.3 (C), 137.0 (C), 130.3 (CH), 128.6 (C), 128.3 (CH), 126.0 (CH), 125.0 (CH), 121.4 (CH), 117.5 (CH), 94.5 ($CH_2$), 90.4 (C), 82.6 (C), 56.3 ($CH_3$), 28.2 ($CH_3$), 0.0 ($CH_3$), -0.7 ($CH_3$); HRMS (ESI) calcd for $C_{31}H_{35}O_8Si$ $[M+H]^+$ 563.2096, found 563.2091.

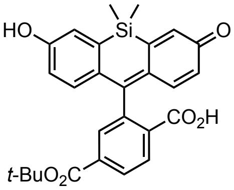


**6-(*tert*-Butoxycarbonyl)-Si-fluorescein (S48):** *Via bis-(trimethylsilylethoxymethyl ether) deprotection:* To a solution of 6-(*tert*-butoxycarbonyl)-Si-fluorescein bis(trimethylsilylethoxymethyl ether) (**S46**; 147 mg, 0.200 mmol) in 2:1 EtOAc/*t*-BuOH (3 mL) was added 4 M HCl in dioxane (1 mL), and the resulting red-pink solution was stirred at room temperature for 1 h. It was subsequently diluted with water and extracted with EtOAc (2×). The organic extracts were washed with brine, dried over anhydrous $MgSO_4$, filtered, and concentrated *in vacuo*. Silica gel chromatography (5–100% EtOAc/hexanes, linear gradient) afforded **S48** (90 mg, 95%) as an off-white solid.

*Via bis-(methoxymethyl ether) deprotection:* To a solution of 6-(*tert*-butoxycarbonyl)-Si-fluorescein bis(methoxymethyl ether) (**S47**; 270 mg, 0.480 mmol) in 2:1 EtOAc/*t*-BuOH (7.5 mL) was added 4 M HCl in dioxane (2.5 mL), and the resulting red-pink solution was stirred at room temperature for 2 h. It was subsequently diluted with water and extracted with EtOAc (2×). The organic extracts were washed with brine, dried over anhydrous $MgSO_4$, filtered, and concentrated *in vacuo*. Silica gel chromatography (5–100% EtOAc/hexanes, linear gradient) afforded **S48** (218 mg, 96%) as a white solid.

$^1H$ NMR (DMSO-$d_6$, 400 MHz) δ 9.75 (s, 2H), 8.07 (AB of ABX, $\nu_A$ = 3233.3 Hz, $J_{AX}$ = 1.3 Hz, $\nu_B$ = 3226.5 Hz, $J_{BX}$ = 0.7 Hz, $J_{AB}$ = 8.0 Hz, 2H), 7.59 (dd, $J$ = 1.4, 0.7 Hz, 1H), 7.14 (d, $J$ = 2.6 Hz, 2H), 6.82 (d, $J$ = 8.8 Hz, 2H), 6.75 (dd, $J$ = 8.8, 2.7 Hz, 2H), 1.49 (s, 9H), 0.62 (s, 3H), 0.52 (s, 3H); $^{13}C$ NMR (DMSO-$d_6$, 101 MHz) δ 169.3 (C), 163.4 (C), 156.9 (C), 155.6 (C), 136.9 (C), 135.5 (C), 133.7 (C), 129.9 (CH), 127.9 (CH), 127.2 (C), 126.2 (CH), 123.4 (CH), 120.1 (CH), 117.5 (CH), 89.8 (C), 82.2 (C), 27.6 ($CH_3$), -0.6 (CH3), -0.7 ($CH_3$); HRMS (ESI) calcd for $C_{27}H_{27}O_6Si$ $[M+H]^+$ 475.1571, found 475.1569.

#### HM-Si-Fluorescein Ditriflate 2

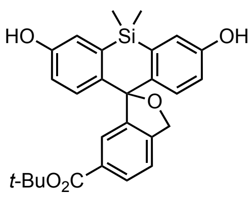


**6-*tert*-Butoxycarbonyl-hydroxymethyl-Si-fluorescein (HM-SiFl-$CO_2$*t*-Bu, S3):** To a solution of *tert*-butyl 3-iodo-4-(((triisopropylsilyl)oxy)methyl)benzoate (**S2**; 4.98 g, 10.15 mmol, 1.5 eq) in THF (25 mL) at -40 °C was added *i*-PrMgCl (2.0 M in THF; 5.08 mL, 10.15 mmol, 1.5 eq). The reaction was then gradually warmed to -25 °C over 2 h while stirring. A solution of 3,7-bis(methoxymethoxy)-5,5-dimethyldibenzo[*b*,*e*]silin-10(5*H*)-one (**S1**; 2.43 g, 6.77 mmol) in THF (10 mL) was added dropwise; the reaction was then warmed to room temperature and stirred for 2 h. It was subsequently quenched with saturated $NH_4Cl$, diluted with water, and extracted with EtOAc (2×). The combined organic extracts were washed with brine, dried over anhydrous $MgSO_4$, filtered, and concentrated *in vacuo*. The crude residue was redissolved in 2:1 EtOAc/*t*-BuOH (75 mL). Following the addition of 4 M HCl in dioxane (25 mL), the resulting red solution was stirred at room temperature for 6 h. It was then diluted with water and extracted with EtOAc (2×). The organic extracts were washed with brine, dried over anhydrous $MgSO_4$, filtered, and evaporated. Silica gel chromatography (10–75% EtOAc/hexanes, linear gradient) afforded **S3** (2.70 g, 87%) as a pale orange foam. $^1$H NMR (DMSO-$d_6$, 400 MHz) δ 9.46 (s, 2H), 7.75 (dd, *J* = 7.9, 1.5 Hz, 1H), 7.49 (d, *J* = 8.0 Hz, 1H), 7.24 (d, *J* = 1.4 Hz, 1H), 7.22 (d, *J* = 8.8 Hz, 2H), 7.01 (d, *J* = 2.7 Hz, 2H), 6.74 (dd, *J* = 8.8, 2.7 Hz, 2H), 5.62 (s, 2H), 1.43 (s, 9H), 0.61 (s, 3H), 0.45 (s, 3H); $^{13}$C NMR ($CDCl_3$, 101 MHz) δ 164.3 (C), 155.8 (C), 149.1 (C), 141.9 (C), 141.3 (C), 132.8 (C), 130.8 (C), 128.7 (CH), 128.2 (CH), 122.7 (CH), 122.3 (CH), 119.0 (CH), 117.5 (CH), 91.4 (C), 81.0 (C), 74.0 ($CH_2$), 27.7 ($CH_3$), 0.7 ($CH_3$), -0.6 ($CH_3$); HRMS (ESI) calcd for $C_{27}H_{29}O_5Si$ $[M+H]^+$ 461.1779, found 461.1779.

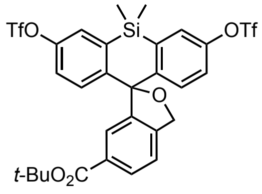


**6-*tert*-Butoxycarbonyl-hydroxymethyl-Si-fluorescein ditriflate ($Tf_2$-HM-SiFl-$CO_2$*t*-Bu, 2):** 6-*tert*-Butoxycarbonyl-hydroxymethyl-Si-fluorescein (**S3**; 2.75 g, 5.97 mmol) was taken up in DMF (60 mL). $Na_2CO_3$ (5.06 g, 47.76 mmol, 8 eq) and *N*-phenyl-bis(trifluoromethanesulfonimide) (8.53 g, 23.88 mmol, 4 eq) were added in succession, and the reaction was stirred at room temperature for 18 h. The resulting off-white suspension was diluted with water and extracted with EtOAc (2×). The combined organic extracts were washed with water and brine, dried over anhydrous $MgSO_4$, filtered, and concentrated *in vacuo*. Flash chromatography (0–20% EtOAc/hexanes, linear gradient) yielded 4.05 g (94%) of **2** as a white foam. $^1$H NMR ($CDCl_3$, 400 MHz) δ 7.99 (dd, *J* = 8.0, 1.4 Hz, 1H), 7.65 (d, *J* = 1.4 Hz, 1H), 7.50 (d, *J* = 2.7 Hz, 2H), 7.44 (d, *J* = 8.9 Hz, 2H), 7.39 (d, *J* = 8.0 Hz, 1H), 7.20 (dd, *J* = 8.9, 2.7 Hz, 2H), 5.49 (s, 2H), 1.55 (s, 9H), 0.78 (s, 3H), 0.61 (s, 3H); $^{19}$F NMR ($CDCl_3$, 376 MHz) δ -73.32 (s); $^{13}$C NMR

($CDCl_3$, 101 MHz) δ 165.0 (C), 150.1 (C), 148.9 (C), 144.8 (C), 142.4 (C), 137.0 (C), 132.6 (C), 130.1 (CH), 129.5 (CH), 126.0 (CH), 124.8 (CH), 122.8 (CH), 121.9 (CH), 118.9 (q, $^{1}J_{CF}$ = 320.9 Hz, $CF_3$), 91.8 (C), 81.8 (C), 74.1 ($CH_2$), 28.3 ($CH_3$), 0.0 ($CH_3$), -0.5 ($CH_3$); HRMS (ESI) calcd for $C_{29}H_{27}F_6O_9S_2Si$ $[M+H]^+$ 725.0764, found 725.0755.

#### References


(1) Egawa, T.; Koide, Y.; Hanaoka, K.; Komatsu, T.; Terai, T.; Nagano, T. Development of a fluorescein analogue, TokyoMagenta, as a novel scaffold for fluorescence probes in red region. *Chem. Commun.* **2011**, *47*, 4162-4164.

(2) Grimm, J. B.; Brown, T. A.; Tkachuk, A. N.; Lavis, L. D. General synthetic method for Si-fluoresceins and Si-rhodamines. *ACS Cent. Sci.* **2017**, *3*, 975-985.

(3) Grimm, J. B.; Sung, A. J.; Legant, W. R.; Hulamm, P.; Matlosz, S. M.; Betzig, E.; Lavis, L. D. Carbofluoresceins and carborhodamines as scaffolds for high-contrast fluorogenic probes. *ACS Chem. Biol.* **2013**, *8*, 1303-1310.

(4) Grimm, J. B.; Muthusamy, A. K.; Liang, Y.; Brown, T. A.; Lemon, W. C.; Patel, R.; Lu, R.; Macklin, J. J.; Keller, P. J.; Ji, N.; Lavis, L. D. A general method to fine-tune fluorophores for live-cell and in vivo imaging. *Nat. Methods* **2017**, *14*, 987-994.

(5) Grimm, J. B.; Gruber, T. D.; Ortiz, G.; Brown, T. A.; Lavis, L. D. Virginia Orange: A versatile, red-shifted fluorescein scaffold for single-and dual-input fluorogenic probes. *Bioconjugate Chem.* **2016**, *27*, 474–480.

(6) Martineau, M.; Somasundaram, A.; Grimm, J. B.; Gruber, T. D.; Choquet, D.; Taraska, J. W.; Lavis, L. D.; Perrais, D. Semisynthetic fluorescent pH sensors for imaging exocytosis and endocytosis. *Nat. Commun.* **2017**, *8*, 1412.

(7) Grimm, J. B.; English, B. P.; Chen, J.; Slaughter, J. P.; Zhang, Z.; Revyakin, A.; Patel, R.; Macklin, J. J.; Normanno, D.; Singer, R. H.; Lionnet, T.; Lavis, L. D. A general method to improve fluorophores for live-cell and single-molecule microscopy. *Nat. Methods* **2015**, *12*, 244–250.

(8) Uno, S.-n.; Kamiya, M.; Yoshihara, T.; Sugawara, K.; Okabe, K.; Tarhan, M. C.; Fujita, H.; Funatsu, T.; Okada, Y.; Tobita, S.; Urano, Y. A spontaneously blinking fluorophore based on intramolecular spirocyclization for live-cell super-resolution imaging. *Nat. Chem.* **2014**, *6*, 681-689.

(9) Iwatate, R. J.; Kamiya, M.; Umezawa, K.; Kashima, H.; Nakadate, M.; Kojima, R.; Urano, Y. Silicon rhodamine-based near-infrared fluorescent probe for γ-glutamyltransferase. *Bioconjugate Chem.* **2018**, *29*, 241-244.

(10) Bucevičius, J.; Gilat, T.; Lukinavičius, G. Far-red switching DNA probes for live cell nanoscopy. *Chem. Commun.* **2020**, *56*, 14797-14800.

(11) Chi, W.; Qiao, Q.; Wang, C.; Zheng, J.; Zhou, W.; Xu, N.; Wu, X.; Jiang, X.; Tan, D.; Xu, Z.; Liu, X. Descriptor ΔG(C-O) enables the quantitative design of spontaneously blinking rhodamines for live-cell super-resolution imaging. *Angew. Chem. Int. Edit.* **2020**, *59*, 20215-20223.

(12) Zheng, Q.; Ayala, A. X.; Chung, I.; Weigel, A. V.; Ranjan, A.; Falco, N.; Grimm, J. B.; Tkachuk, A. N.; Wu, C.; Lippincott-Schwartz, J.; Singer, R. H.; Lavis, L. D. Rational design of fluorogenic and spontaneously blinking labels for super-resolution imaging. *ACS Cent. Sci.* **2019**, *5*, 1602-1613.

(13) Best, Q. A.; Sattenapally, N.; Dyer, D. J.; Scott, C. N.; McCarroll, M. E. pH-Dependent Si-fluorescein hypochlorous acid fluorescent probe: spirocycle ring-opening and excess hypochlorous acid-induced chlorination. *J. Am. Chem. Soc* **2013**, *135*, 13365-13370.

(14) Butkevich, A. N.; Belov, V. N.; Kolmakov, K.; Sokolov, V. V.; Shojaei, H.; Sidenstein, S. C.; Kamin, D.; Matthias, J.; Vlijm, R.; Engelhardt, J.; Hell, S. W. Hydroxylated fluorescent dyes for live-cell labeling: synthesis, spectra and super-resolution STED. *Chem. Eur. J.* **2017**, *23*, 12114-12119.

(15) Smaga, L. P.; Pino, N. W.; Ibarra, G. E.; Krishnamurthy, V.; Chan, J. A photoactivatable formaldehyde donor with fluorescence monitoring reveals threshold to arrest cell migration. *J. Am. Chem. Soc* **2020**, *142*, 680-684.

(16) Bearrood, T. E.; Aguirre-Figueroa, G.; Chan, J. Rational design of a red fluorescent sensor for ALDH1A1 displaying enhanced cellular uptake and reactivity. *Bioconjugate Chem.* **2020**, *31*, 224-228.

(17) Minta, A.; Kao, J. P.; Tsien, R. Y. Fluorescent indicators for cytosolic calcium based on rhodamine and fluorescein chromophores. *J. Biol. Chem.* **1989**, *264*, 8171–8178.

(18) Butkevich, A. N.; Mitronova, G. Y.; Sidenstein, S. C.; Klocke, J. L.; Kamin, D.; Meineke, D. N.; D'Este, E.; Kraemer, P. T.; Danzl, J. G.; Belov, V. N.; Hell, S. W. Fluorescent rhodamines and fluorogenic carbopyronines for super-resolution STED microscopy in living cells. *Angew. Chem. Int. Edit.* **2016**, *55*, 3290-3294.

(19) Bucevičius, J.; Kostiuk, G.; Gerasimaitė, R.; Gilat, T.; Lukinavičius, G. Enhancing the biocompatibility of rhodamine fluorescent probes by a neighbouring group effect. *Chem. Sci.* **2020**, *11*, 7313-7323.

(20) Kolmakov, K.; Belov, V. N.; Wurm, C. A.; Harke, B.; Leutenegger, M.; Eggeling, C.; Hell, S. W. A versatile route to red-emitting carbopyronine dyes for optical microscopy and nanoscopy. *Eur. J. Org. Chem.* **2010**, *2010*, 3593-3610.

(21) Lukinavičius, G.; Umezawa, K.; Olivier, N.; Honigmann, A.; Yang, G.; Plass, T.; Mueller, V.; Reymond, L.; Corrêa Jr, I. R.; Luo, Z. G.; Schultz, C.; Lemke, E. A.; Heppenstall, P.; Eggeling, C.; Manley, S.; Johnsson, K. A near-infrared fluorophore for live-cell super-resolution microscopy of cellular proteins. *Nat. Chem.* **2013**, *5*, 132–139.

(22) Grimm, J. B.; Klein, T.; Kopek, B. G.; Shtengel, G.; Hess, H. F.; Sauer, M.; Lavis, L. D. Synthesis of a far-red photoactivatable silicon-containing rhodamine for super-resolution microscopy. *Angew. Chem. Int. Edit.* **2016**, *55*, 1723-1727.

(23) Butkevich, A. N.; Ta, H.; Ratz, M.; Stoldt, S.; Jakobs, S.; Belov, V. N.; Hell, S. W. Two-color 810 nm STED nanoscopy of living cells with endogenous SNAP-tagged fusion proteins. *ACS Chem. Biol.* **2018**, *13*, 475-480.

(24) Krasovskiy, A.; Knochel, P. A LiCl-mediated Br/Mg exchange reaction for the preparation of functionalized aryl- and heteroarylmagnesium compounds from organic bromides. *Angew. Chem. Int. Edit.* **2004**, *43*, 3333-3336.

(25) Boymond, L.; Rottländer, M.; Cahiez, G.; Knochel, P. Preparation of highly functionalized Grignard reagents by an iodine-magnesium exchange reaction and its application in solid-phase synthesis. *Angew. Chem. Int. Edit.* **1998**, *37*, 1701-1703.

(26) Abarbri, M.; Thibonnet, J.; Bérillon, L.; Dehmel, F.; Rottländer, M.; Knochel, P. Preparation of new polyfunctional magnesiated heterocycles using a chlorine-, bromine-, or iodine-magnesium exchange. *J. Org. Chem.* **2000**, *65*, 4618-4634.

(27) Knochel, P.; Dohle, W.; Gommermann, N.; Kneisel, F. F.; Kopp, F.; Korn, T.; Sapountzis, I.; Vu, V. A. Highly functionalized organomagnesium reagents prepared through halogen-metal exchange. *Angew. Chem. Int. Edit.* **2003**, *42*, 4302-4320.

(28) Hari, Y.; Kondo, R.; Date, K.; Aoyama, T. Facile synthesis of 2-unsubstituted benzofuran-3-carboxylates using diazo(trimethylsilyl)methylmagnesium bromide. *Tetrahedron* **2009**, *65*, 8708-8713.

(29) Cheng, H. G.; Zhang, R. M.; Wang, M.; Zeng, X. F.; Xie, C. S. Convergent assembly of enantioenriched tetrahydrobenzofuro[2,3-b]pyrrole scaffolds by Ag-catalyzed asymmetric domino reaction of isocyanoacetates. *Asian J Org Chem* **2018**, *7*, 1075-1079.

(30) Kuribayashi, T.; Kubota, H.; Fukuda, T.; Takano, R.; Tsuji, T.; Sasaki, K.; Tanaka, N. 5-Hydroxypyrimidine-4-carboxamide compound. US 2011/0112103 A1, May 12, 2011.

(31) Mathias, L. J. Esterification and alkylation reactions employing isoureas. *Synthesis* **1979**, 561-576.

(32) Widmer, U. A convenient preparation of t-butyl esters. *Synthesis* **1983**, *1983*, 135-136.

(33) Klapars, A.; Buchwald, S. L. Copper-catalyzed halogen exchange in aryl halides: an aromatic Finkelstein reaction. *J. Am. Chem. Soc.* **2002**, *124*, 14844-14845.
