## Supplementary Note 1 for "A series of spontaneously blinking dyes for super-resolution microscopy"

SUPPLEMENTARY NOTE 2

### A series of spontaneously blinking dyes for super-resolution microscopy

Katie L. Holland, Sarah E. Plutkis, Timothy A. Daugird, Abhishek Sau, Jonathan B. Grimm, Brian P. English, Qinsi Zheng, Sandeep Dave, Fariha Rahman, Liangqi Xie, Peng Dong, Ariana N. Tkachuk, Timothy A. Brown, Robert H. Singer, Catherine G. Galbraith, Siegfried M. Musser, Wesley R. Legant, and Luke D. Lavis*

*

#### Experimental Information: Synthesis, Characterization, and Application of JFb Dyes

| Page | Contents |
|---|---|
| S2 | General Experimental Information for Synthesis |
| S3–S17 | Experimentals and Characterization Data for All Compounds |
| S3 | JFb Synthesis via Pd-Catalyzed C–N Cross-Coupling |
| S7 | Deprotection of JFb *tert*-Butyl Esters |
| S11 | Synthesis of JFb–HaloTag Ligands and Other Derivatives |

#### General Experimental Information for Synthesis

Commercial reagents were obtained from reputable suppliers and used as received. All solvents were purchased in septum-sealed bottles stored under an inert atmosphere. All reactions were sealed with septa through which a nitrogen atmosphere was introduced unless otherwise noted. Reactions were conducted in round-bottomed flasks or septum-capped crimp-top vials containing Teflon-coated magnetic stir bars. Heating of reactions was accomplished with a silicon oil bath or an aluminum reaction block on top of a stirring hotplate equipped with an electronic contact thermometer to maintain the indicated temperatures.

Reactions were monitored by thin layer chromatography (TLC) on precoated TLC glass plates (silica gel 60 $F_{254}$, 250 μm thickness) or by LC/MS (Phenomenex Kinetex 2.1 mm × 30 mm 2.6 μm C18 column; 5 μL injection; 5–98% $MeCN/H_2O$, linear gradient, with constant 0.1% v/v $HCO_2H$ additive; 6 min run; 0.5 mL/min flow; ESI; positive ion mode). TLC chromatograms were visualized by UV illumination or developed with *p*-anisaldehyde, ceric ammonium molybdate, or $KMnO_4$ stain. Reaction products were purified by flash chromatography on an automated purification system using pre-packed silica gel columns or by preparative HPLC (Phenomenex Gemini–NX 30 × 150 mm 5 μm C18 column). Analytical HPLC analysis was performed with an Agilent Eclipse XDB 4.6 × 150 mm 5 μm C18 column under the indicated conditions. High-resolution mass spectrometry was performed by the High Resolution Mass Spectrometry Facility at the University of Iowa.

NMR spectra were recorded on a 400 MHz spectrometer. $^{1}H$ and $^{13}C$ chemical shifts were referenced to TMS or residual solvent peaks, and $^{19}F$ chemical shifts were referenced to $CFCl_3$. Data for $^{1}H$ NMR spectra are reported as follows: chemical shift (δ ppm), multiplicity (s = singlet, d = doublet, t = triplet, q = quartet, dd = doublet of doublets, m = multiplet), coupling constant (Hz), integration. Data for $^{13}C$ NMR spectra are reported by chemical shift (δ ppm) with hydrogen multiplicity (C, CH, $CH_2$, $CH_3$) information obtained from DEPT spectra.

#### JFB Synthesis via Pd-Catalyzed C–N Cross-Couplings

**General method S1: C–N cross-coupling of ditriflate.** The following procedure for 6-*tert*-butoxycarbonyl-$JF_{614}$b (**S11**) is representative. A vial was charged with ditriflate **2** (300 mg, 0.414 mmol), 3,3-difluoroazetidine hydrochloride (129 mg, 0.994 mmol, 2.4 eq), RuPhos-G3-palladacycle (34.6 mg, 41.4 μmol, 0.1 eq), RuPhos (19.3 mg, 41.4 μmol, 0.1 eq), and $Cs_2CO_3$ (647 mg, 1.99 mmol, 4.8 eq). The vial was sealed and evacuated/backfilled with nitrogen (3×). Dioxane (2.5 mL) was added, and the reaction was flushed again with nitrogen (3×). The reaction was then stirred at 100 °C for 3 h. It was subsequently cooled to room temperature, filtered through Celite with $CH_2Cl_2$, and concentrated to dryness. Purification by silica gel chromatography (0–30% EtOAc/hexanes, linear gradient) afforded **S11** (232 mg, 92%) as a white solid.

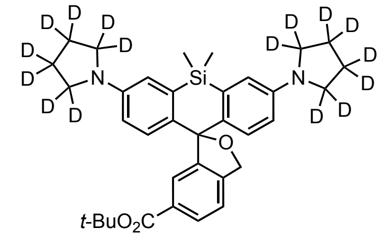


**6-*tert*-Butoxycarbonyl-$JFX_{650}$b (S4):** The title compound (27%, pale yellow solid) was prepared from ditriflate **2** and pyrrolidine-2,2,3,3,4,4,5,5-$d_8$ according to method S1. $^1$H NMR ($CDCl_3$, 400 MHz) δ 7.89 (dd, $J$ = 7.9, 1.5 Hz, 1H), 7.58 (d, $J$ = 1.4 Hz, 1H), 7.30 (d, $J$ = 7.9 Hz, 1H), 7.08 (d, $J$ = 8.8 Hz, 2H), 6.77 (d, $J$ = 2.8 Hz, 2H), 6.49 (dd, $J$ = 8.8, 2.7 Hz, 2H), 5.40 (s, 2H), 1.50 (s, 9H), 0.66 (s, 3H), 0.54 (s, 3H); $^{13}$C NMR ($CDCl_3$, 101 MHz) δ 165.6 (C), 148.1 (C), 146.3 (C), 143.1 (C), 137.8 (C), 134.7 (C), 131.8 (C), 128.7 (CH), 128.5 (CH), 125.1 (CH), 121.0 (CH), 115.6 (CH), 113.3 (CH), 92.8 (C), 81.0 (C), 73.1 ($CH_2$), 28.3 ($CH_3$), 0.3 ($CH_3$), -0.0 ($CH_3$); HRMS (ESI) calcd for $C_{35}H_{27}D_{16}N_2O_3Si$ $[M+H]^+$ 583.4042, found 583.4034.

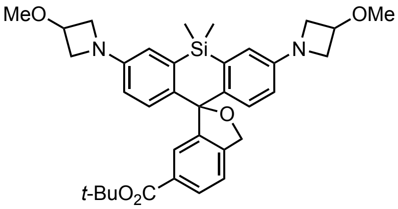


**6-*tert*-Butoxycarbonyl-$JF_{639}$b (S5):** The title compound (91%, white solid) was prepared from ditriflate **2** and 3-methoxyazetidine hydrochloride according to method S1. $^1$H NMR ($CDCl_3$, 400 MHz) δ 7.91 (dd, $J$ = 7.9, 1.5 Hz, 1H), 7.59 (d, $J$ = 1.6 Hz, 1H), 7.31 (d, $J$ = 8.0 Hz, 1H), 7.06 (d, $J$ = 8.6 Hz, 2H), 6.67 (d, $J$ = 2.7 Hz, 2H), 6.38 (dd, $J$ = 8.6, 2.6 Hz, 2H), 5.37 (s, 2H), 4.33 (tt, $J$ = 6.1, 4.5 Hz, 2H), 4.14 – 4.07 (m, 4H), 3.75 – 3.68 (m, 4H), 3.32 (s, 6H), 1.51 (s, 9H), 0.64 (s, 3H), 0.52 (s, 3H); $^{13}$C NMR ($CDCl_3$, 101 MHz) δ 165.6 (C), 149.8 (C), 147.3 (C), 143.2 (C), 139.8 (C), 134.8 (C), 131.8 (C), 128.9 (CH), 128.2 (CH), 125.2 (CH), 121.2 (CH), 115.9 (CH), 113.3 (CH), 92.7 (C), 81.2 (C), 73.1 ($CH_2$), 70.2 (CH), 59.0 ($CH_2$), 56.2 ($CH_3$), 28.3 ($CH_3$), 0.3 ($CH_3$), -0.3 ($CH_3$); HRMS (ESI) calcd for $C_{35}H_{43}N_2O_5Si$ $[M+H]^+$ 599.2936, found 599.2933.

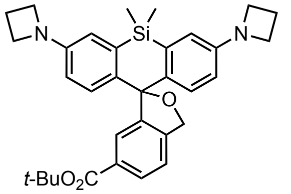


**6-*tert*-Butoxycarbonyl-JF$_{646}$b (S6):** The title compound (91%, off-white solid) was prepared from ditriflate **2** and azetidine according to method S1. $^1$H NMR (CDCl$_3$, 400 MHz) δ 7.90 (dd, *J* = 7.9, 1.5 Hz, 1H), 7.60 (d, *J* = 1.4 Hz, 1H), 7.30 (d, *J* = 8.0 Hz, 1H), 7.05 (d, *J* = 8.6 Hz, 2H), 6.65 (d, *J* = 2.7 Hz, 2H), 6.35 (dd, *J* = 8.7, 2.7 Hz, 2H), 5.36 (s, 2H), 3.88 (t, *J* = 7.2 Hz, 8H), 2.34 (p, *J* = 7.2 Hz, 4H), 1.51 (s, 9H), 0.64 (s, 3H), 0.52 (s, 3H); $^{13}$C NMR (CDCl$_3$, 101 MHz) δ 165.6 (C), 150.4 (C), 147.4 (C), 143.3 (C), 139.4 (C), 134.7 (C), 131.8 (C), 128.9 (CH), 128.1 (CH), 125.3 (CH), 121.1 (CH), 115.5 (CH), 112.9 (CH), 92.8 (C), 81.2 (C), 73.0 (CH$_2$), 52.5 (CH$_2$), 28.3 (CH$_3$), 17.1 (CH$_2$), 0.3 (CH$_3$), -0.3 (CH$_3$); HRMS (ESI) calcd for $C_{33}H_{39}N_2O_3Si$ [M+H]$^+$ 539.2724, found 539.2725.

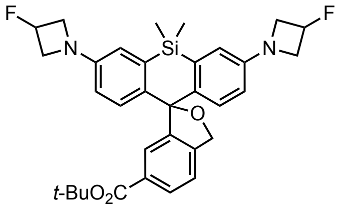


**6-*tert*-Butoxycarbonyl-JF$_{635}$b (S7):** The title compound (25%, light blue solid) was prepared from ditriflate **2** and 3-fluoroazetidine hydrochloride according to method S1. $^1$H NMR (CDCl$_3$, 400 MHz) δ 7.92 (dd, *J* = 7.9, 1.5 Hz, 1H), 7.60 (d, *J* = 1.4 Hz, 1H), 7.32 (d, *J* = 8.0 Hz, 1H), 7.09 (d, *J* = 8.6 Hz, 2H), 6.68 (d, *J* = 2.7 Hz, 2H), 6.39 (dd, *J* = 8.7, 2.7 Hz, 2H), 5.41 (dtt, $^2J_{HF}$ = 57.0 Hz, *J* = 6.0, 3.7 Hz, 2H), 5.37 (s, 2H), 4.24 – 4.14 (m, 4H), 4.02 – 3.91 (m, 4H), 1.52 (s, 9H), 0.66 (s, 3H), 0.53 (s, 3H); $^{19}$F NMR (CDCl$_3$, 376 MHz) δ -180.44 (dtt, $J_{FH}$ = 56.9, 23.9, 18.0 Hz); $^{13}$C NMR (CDCl$_3$, 101 MHz) δ 165.5 (C), 149.36 (d, $^4J_{CF}$ = 1.5 Hz, C), 147.1 (C), 143.2 (C), 140.3 (C), 134.9 (C), 131.9 (C), 129.0 (CH), 128.2 (CH), 125.2 (CH), 121.2 (CH), 116.1 (CH), 113.5 (CH), 92.6 (C), 82.94 (d, $^1J_{CF}$ = 204.5 Hz, CHF), 81.3 (C), 73.1 (CH$_2$), 59.68 (d, $^2J_{CF}$ = 23.5 Hz, CH$_2$), 28.3 (CH$_3$), 0.3 (CH$_3$), -0.3 (CH$_3$); HRMS (ESI) calcd for $C_{33}H_{37}F_2N_2O_3Si$ [M+H]$^+$ 575.2536, found 575.2535.

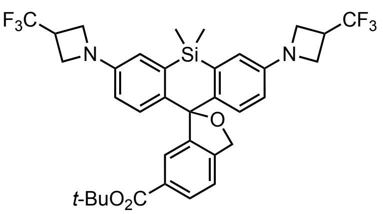


**6-*tert*-Butoxycarbonyl-JF$_{626}$b (S8):** The title compound (92%, white solid) was prepared from ditriflate **2** and 3-trifluoromethyl-azetidine hydrochloride according to method S1. $^1$H NMR (CDCl$_3$, 400 MHz) δ 7.92 (dd, *J* = 7.9, 1.4 Hz, 1H), 7.60 (d, *J* = 1.4 Hz, 1H), 7.32 (d, *J* = 8.0 Hz, 1H), 7.10 (d, *J* = 8.6 Hz, 2H), 6.66 (d, *J* = 2.6 Hz, 2H), 6.37 (dd, *J* = 8.6, 2.7 Hz, 2H), 5.37 (s, 2H), 4.09 – 4.02 (m, 4H), 3.99 – 3.92 (m, 4H), 3.45 – 3.30 (m, 2H), 1.52 (s, 9H),

0.65 (s, 3H), 0.53 (s, 3H); $^{19}$F NMR ($CDCl_3$, 376 MHz) δ -73.30 (d, $^3J_{FH}$ = 8.8 Hz); $^{13}$C NMR ($CDCl_3$, 101 MHz) δ 165.5 (C), 148.9 (C), 147.0 (C), 143.2 (C), 140.4 (C), 135.0 (C), 131.9 (C), 129.0 (CH), 128.3 (CH), 126.5 (q, $^1J_{CF}$ = 275.2 Hz, $CF_3$), 125.2 (CH), 121.2 (CH), 115.6 (CH), 112.9 (CH), 92.6 (C), 81.3 (C), 73.1 ($CH_2$), 51.4 (q, $^3J_{CF}$ = 3.6 Hz, $CH_2$), 33.2 (q, $^2J_{CF}$ = 32.0 Hz, CH), 28.3 ($CH_3$), 0.3 ($CH_3$), -0.4 ($CH_3$); HRMS (ESI) calcd for $C_{35}H_{37}F_6N_2O_3Si$ $[M+H]^+$ 675.2472, found 675.2471.

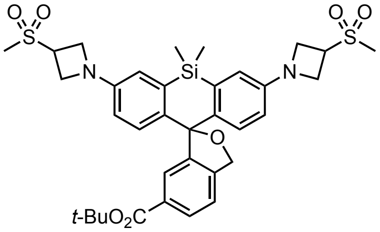


**6-*tert*-Butoxycarbonyl-JF$_{630}$b (S9):** The title compound (41%, light blue solid) was prepared from ditriflate **2** and 3-methylsulfonyl-azetidine hydrochloride according to method S1. $^1$H NMR ($CDCl_3$, 400 MHz) δ 7.91 (dd, $J$ = 7.9, 1.4 Hz, 1H), 7.57 (d, $J$ = 1.7 Hz, 1H), 7.32 (d, $J$ = 8.0 Hz, 1H), 7.13 (d, $J$ = 8.6 Hz, 2H), 6.68 (d, $J$ = 2.7 Hz, 2H), 6.41 (dd, $J$ = 8.6, 2.7 Hz, 2H), 5.40 (s, 2H), 4.27 – 4.17 (m, 8H), 4.10 – 4.01 (m, 2H), 2.99 (s, 6H), 1.52 (s, 9H), 0.66 (s, 3H), 0.53 (s, 3H); $^{13}$C NMR ($CDCl_3$, 101 MHz) δ 165.4 (C), 148.5 (C), 147.0 (C), 142.9 (C), 141.2 (C), 134.8 (C), 132.0 (C), 129.1 (CH), 128.3 (CH), 125.0 (CH), 121.3 (CH), 115.9 (CH), 113.4 (CH), 92.5 (C), 81.3 (C), 73.3 ($CH_2$), 52.6 ($CH_2$), 51.9 (CH), 38.1 ($CH_3$), 28.3 ($CH_3$), 0.3 ($CH_3$), -0.2 ($CH_3$); HRMS (ESI) calcd for $C_{35}H_{43}N_2O_7S_2Si$ $[M+H]^+$ 695.2275, found 695.2274.

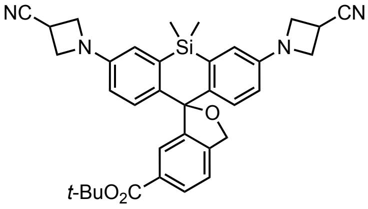


**6-*tert*-Butoxycarbonyl-JF$_{629}$b (S10):** The title compound (49%, pale green solid) was prepared from ditriflate **2** and azetidine-3-carbonitrile hydrochloride according to method S1. $^1$H NMR ($CDCl_3$, 400 MHz) δ 7.92 (dd, $J$ = 8.0, 1.5 Hz, 1H), 7.59 (d, $J$ = 1.4 Hz, 1H), 7.33 (d, $J$ = 7.9 Hz, 1H), 7.12 (d, $J$ = 8.6 Hz, 2H), 6.66 (d, $J$ = 2.6 Hz, 2H), 6.38 (dd, $J$ = 8.6, 2.7 Hz, 2H), 5.39 (s, 2H), 4.23 – 4.17 (m, 4H), 4.11 – 4.06 (m, 4H), 3.58 (tt, $J$ = 8.4, 6.4 Hz, 2H), 1.52 (s, 9H), 0.67 (s, 3H), 0.53 (s, 3H); $^{13}$C NMR ($CDCl_3$, 101 MHz) δ 165.4 (C), 148.7 (C), 146.8 (C), 143.0 (C), 141.2 (C), 134.9 (C), 132.0 (C), 129.1 (CH), 128.3 (CH), 125.0 (CH), 121.3 (CH), 119.9 (C), 115.9 (CH), 113.4 (CH), 92.4 (C), 81.3 (C), 73.3 ($CH_2$), 55.4 ($CH_2$), 28.3 ($CH_3$), 18.5 (CH), 0.3 ($CH_3$), -0.3 ($CH_3$); HRMS (ESI) calcd for $C_{35}H_{37}N_4O_3Si$ $[M+H]^+$ 589.2629, found 589.2630.

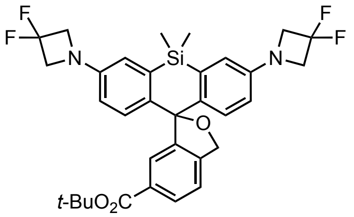


**6-*tert*-Butoxycarbonyl-JF$_{614}$b (S11):** The title compound (92%, white solid) was prepared from ditriflate **2** and 3,3-difluoroazetidine hydrochloride according to method S1. $^1$H NMR ($CDCl_3$, 400 MHz) δ 7.93 (dd, $J$ = 7.9, 1.4 Hz, 1H), 7.61 (d, $J$ = 1.4 Hz, 1H), 7.33 (d, $J$ = 7.9 Hz, 1H), 7.14 (d, $J$ = 8.6 Hz, 2H), 6.71 (d, $J$ = 2.7 Hz, 2H), 6.42 (dd, $J$ = 8.6, 2.7 Hz, 2H), 5.39 (s, 2H), 4.23 (t, $^3J_{HF}$ = 11.8 Hz, 8H), 1.52 (s, 9H), 0.67 (s, 3H), 0.54 (s, 3H); $^{19}$F NMR ($CDCl_3$, 376 MHz) δ -99.78 (p, $^3J_{FH}$ = 12.0 Hz); $^{13}$C NMR ($CDCl_3$, 101 MHz) δ 165.5 (C), 148.1 (t, $^4J_{CF}$ = 2.6 Hz, C), 146.8 (C), 143.1 (C), 141.1 (C), 135.1 (C), 133.0 (C), 129.1 (CH), 128.3 (CH), 125.1 (CH), 121.3 (CH), 116.6 (CH), 116.10 (t, $^1J_{CF}$ = 274.6 Hz, $CF_2$), 114.0 (CH), 92.5 (C), 81.3 (C), 73.2 ($CH_2$), 63.5 (t, $^2J_{CF}$ = 25.6 Hz, $CH_2$), 28.3 ($CH_3$), 0.3 ($CH_3$), -0.3 ($CH_3$); HRMS (ESI) calcd for $C_{33}H_{35}F_4N_2O_3Si$ $[M+H]^+$ 611.2348, found 611.2342.

#### DEPROTECTION OF JFB *TERT*-BUTYL ESTERS

**General method S2: *tert*-Butyl ester deprotection.** The following procedure for 6-carboxy-$JF_{614}b$ (**S19**) is representative. 6-*tert*-Butoxycarbonyl-$JF_{614}b$ (**S11**; 215 mg, 0.352 mmol) was taken up in $CH_2Cl_2$ (5 mL), and trifluoroacetic acid (1 mL) was added. The reaction was stirred at room temperature for 4 h. Toluene (5 mL) was added; the reaction mixture was concentrated to dryness and then azeotroped with MeOH (3×). The residue was redissolved in 1:1 THF/MeOH (5 mL), and 1 M NaOH (1.06 mL, 1.06 mmol, 3 eq) was added. After stirring the reaction at room temperature for 1 h, it was acidified with 1 M HCl (1.1 mL), diluted with water, and extracted with EtOAc (2×). The combined organic extracts were washed with brine, dried over anhydrous $MgSO_4$, filtered, and concentrated *in vacuo* to afford 186 mg (95%) of **S19** as a blue solid. Analytical HPLC and NMR indicated that the material was >95% pure and did not require further purification prior to amide coupling.

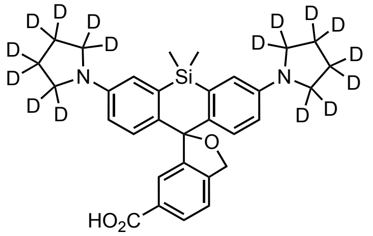

**6-Carboxy-$JFX_{650}b$ (S12):** The title compound (67%, blue solid) was prepared from 6-*tert*-butoxycarbonyl-$JFX_{650}b$ (**S4**) according to method S2. $^1$H NMR ($CDCl_3$, 400 MHz) δ 7.92 (dd, *J* = 7.9, 1.5 Hz, 1H), 7.68 (d, *J* = 1.5 Hz, 1H), 7.30 (d, *J* = 8.0 Hz, 1H), 6.92 (d, *J* = 8.8 Hz, 2H), 6.71 (d, *J* = 2.8 Hz, 2H), 6.39 (dd, *J* = 8.8, 2.8 Hz, 2H), 5.27 (s, 2H), 0.58 (s, 3H), 0.47 (s, 3H); Analytical HPLC: $t_R$ = 11.7 min, >99% purity (10–95% $MeCN/H_2O$, linear gradient, with constant 0.1% v/v TFA additive; 20 min run; 1 mL/min flow; ESI; positive ion mode; detection at 650 nm); HRMS (ESI) calcd for $C_{31}H_{19}D_{16}N_2O_3Si$ $[M+H]^+$ 527.3416, found 527.3409.

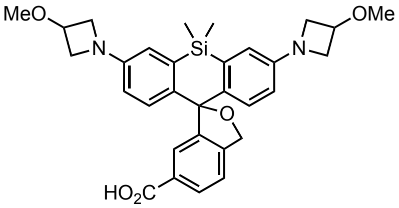

**6-Carboxy-$JF_{639}b$ (S13):** The title compound (95%, dark blue solid) was prepared from 6-*tert*-butoxycarbonyl-$JF_{639}b$ (**S5**) according to method S2. $^1$H NMR ($CDCl_3$, 400 MHz) δ 8.00 (dd, *J* = 8.0, 1.5 Hz, 1H), 7.75 (d, *J* = 1.5 Hz, 1H), 7.37 (d, *J* = 8.0 Hz, 1H), 6.99 (d, *J* = 8.6 Hz, 2H), 6.69 (d, *J* = 2.7 Hz, 2H), 6.36 (dd, *J* = 8.6, 2.7 Hz, 2H), 5.32 (s, 2H), 4.37 – 4.28 (m, 2H), 4.15 – 4.05 (m, 4H), 3.71 (dd, *J* = 7.9, 4.6 Hz, 4H), 3.32 (s, 6H), 0.63 (s, 3H), 0.52 (s, 3H); $^{13}$C NMR ($CDCl_3$, 101 MHz) δ 170.9 (C), 149.8 (C), 147.2 (C), 145.2 (C), 139.5 (C), 135.3 (C), 129.6 (CH), 129.2 (C), 128.1 (CH), 126.5 (CH), 121.5 (CH), 116.3 (CH), 113.3 (CH), 92.7 (C), 72.7 ($CH_2$), 70.2 (CH), 59.0 ($CH_2$), 56.2 ($CH_3$), 0.6 ($CH_3$), -0.9 ($CH_3$); Analytical HPLC: $t_R$ = 10.5 min, >99% purity (10–95% $MeCN/H_2O$, linear gradient, with constant 0.1% v/v TFA additive; 20 min run; 1 mL/min flow; ESI; positive ion mode; detection at 650 nm); HRMS (ESI) calcd for $C_{31}H_{35}N_2O_5Si$ $[M+H]^+$ 543.2310, found 543.2316.

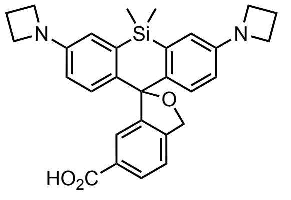


**6-Carboxy-JF$_{646}$b (S14):** The title compound (87%, dark blue solid) was prepared from 6-*tert*-butoxycarbonyl-JF$_{646}$b (**S6**) according to method S2. $^1$H NMR ($CDCl_3$, 400 MHz) δ 7.93 (dd, $J$ = 7.9, 1.4 Hz, 1H), 7.68 (d, $J$ = 1.4 Hz, 1H), 7.29 (d, $J$ = 8.0 Hz, 1H), 6.90 (d, $J$ = 8.6 Hz, 2H), 6.60 (d, $J$ = 2.6 Hz, 2H), 6.27 (dd, $J$ = 8.7, 2.7 Hz, 2H), 5.25 (s, 2H), 3.81 (t, $J$ = 7.2 Hz, 8H), 2.27 (p, $J$ = 7.2 Hz, 4H), 0.56 (s, 3H), 0.45 (s, 3H); $^{13}$C NMR ($CDCl_3$, 101 MHz) δ 170.8 (C), 150.5 (C), 147.3 (C), 145.2 (C), 139.1 (C), 135.2 (C), 129.5 (CH), 129.2 (C), 128.1 (CH), 126.5 (CH), 121.4 (CH), 116.0 (CH), 113.0 (CH), 92.7 (C), 72.6 ($CH_2$), 52.6 ($CH_2$), 17.1 ($CH_2$), 0.6 ($CH_3$), -0.9 ($CH_3$); Analytical HPLC: $t_R$ = 10.8 min, 98.1% purity (10–95% MeCN/$H_2O$, linear gradient, with constant 0.1% v/v TFA additive; 20 min run; 1 mL/min flow; ESI; positive ion mode; detection at 650 nm); HRMS (ESI) calcd for $C_{29}H_{31}N_2O_3Si$ $[M+H]^+$ 483.2098, found 483.2105.

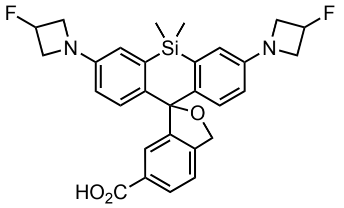


**6-Carboxy-JF$_{635}$b (S15):** The title compound (~100%, blue green solid) was prepared from 6-*tert*-butoxycarbonyl-JF$_{635}$b (**S7**) according to method S2. $^1$H NMR ($CDCl_3$, 400 MHz) δ 8.02 (dd, $J$ = 7.9, 1.5 Hz, 1H), 7.75 (dd, $J$ = 1.4, 0.6 Hz, 1H), 7.38 (dd, $J$ = 8.0, 0.6 Hz, 1H), 7.02 (d, $J$ = 8.6 Hz, 2H), 6.69 (d, $J$ = 2.6 Hz, 2H), 6.37 (dd, $J$ = 8.6, 2.7 Hz, 2H), 5.40 (dtt, $^2J_{FH}$ = 57.1 Hz, $J$ = 5.8, 3.8 Hz, 2H), 5.34 (s, 2H), 4.27 – 4.13 (m, 4H), 4.04 – 3.88 (m, 4H), 0.65 (s, 3H), 0.53 (s, 3H); $^{19}$F NMR ($CDCl_3$, 376 MHz) δ -180.44 (dtt, $J_{FH}$ = 56.9, 23.7, 18.0 Hz); Analytical HPLC: $t_R$ = 12.2 min, >99% purity (10–95% MeCN/$H_2O$, linear gradient, with constant 0.1% v/v TFA additive; 20 min run; 1 mL/min flow; ESI; positive ion mode; detection at 650 nm); HRMS (ESI) calcd for $C_{29}H_{29}F_2N_2O_3Si$ $[M+H]^+$ 519.1910, found 519.1908.

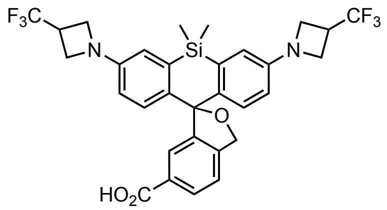


**6-Carboxy-JF$_{626}$b (S16):** The title compound (97%, blue solid) was prepared from 6-*tert*-butoxycarbonyl-JF$_{626}$b (**S8**) according to method S2. $^1$H NMR ($CDCl_3$, 400 MHz) δ 8.02 (dd, *J* = 7.9, 1.5 Hz, 1H), 7.75 (d, *J* = 1.7 Hz, 1H), 7.38 (d, *J* = 7.9 Hz, 1H), 7.03 (d, *J* = 8.6 Hz, 2H), 6.67 (d, *J* = 2.7 Hz, 2H), 6.35 (dd, *J* = 8.7, 2.7 Hz, 2H), 5.34 (s, 2H), 4.11 – 4.00 (m, 4H), 3.99 – 3.91 (m, 4H), 3.44 – 3.31 (m, 2H), 0.64 (s, 3H), 0.52 (s, 3H); $^{19}$F NMR ($CDCl_3$, 376 MHz) δ -73.31 (d, $^3J_{FH}$ = 8.5 Hz); $^{13}$C NMR ($CDCl_3$, 101 MHz) δ 171.3 (C), 149.0 (C), 147.1 (C), 145.3 (C), 140.0 (C), 135.5 (C), 129.7 (CH), 129.1 (C), 128.2 (CH), 126.5 (q, $^1J_{CF}$ = 275.1 Hz, $CF_3$), 126.4 (CH), 121.6 (CH), 115.8 (CH), 112.8 (CH), 92.6 (C), 72.8 ($CH_2$), 51.4 (q, $^3J_{CF}$ = 3.2 Hz, $CH_2$), 33.2 (q, $^2J_{CF}$ = 32.0 Hz, CH), 0.5 ($CH_3$), -0.9 ($CH_3$); Analytical HPLC: $t_R$ = 14.4 min, >99% purity (10–95% MeCN/$H_2O$, linear gradient, with constant 0.1% v/v TFA additive; 20 min run; 1 mL/min flow; ESI; positive ion mode; detection at 650 nm); HRMS (ESI) calcd for $C_{31}H_{29}F_6N_2O_3Si$ [M+H]$^+$ 619.1846, found 619.1849.

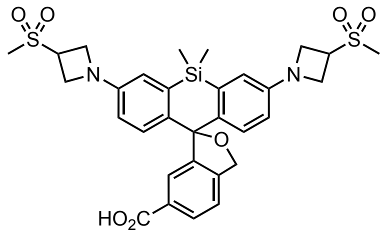


**6-Carboxy-JF$_{630}$b (S17):** The title compound (84%, blue solid) was prepared from 6-*tert*-butoxycarbonyl-JF$_{630}$b (**S9**) according to method S2. $^1$H NMR ($CDCl_3$, 400 MHz) δ 8.01 (dd, *J* = 8.0, 1.5 Hz, 1H), 7.72 (d, *J* = 1.5 Hz, 1H), 7.38 (d, *J* = 8.0 Hz, 1H), 7.07 (d, *J* = 8.6 Hz, 2H), 6.69 (d, *J* = 2.7 Hz, 2H), 6.38 (dd, *J* = 8.6, 2.7 Hz, 2H), 5.37 (s, 2H), 4.26 – 4.14 (m, 8H), 4.12 – 4.02 (m, 2H), 2.98 (s, 6H), 0.65 (s, 3H), 0.52 (s, 3H); Analytical HPLC: $t_R$ = 11.2 min, >99% purity (10–95% MeCN/$H_2O$, linear gradient, with constant 0.1% v/v TFA additive; 20 min run; 1 mL/min flow; ESI; positive ion mode; detection at 650 nm); HRMS (ESI) calcd for $C_{31}H_{35}N_2O_7S_2Si$ [M+H]$^+$ 639.1649, found 639.1644.

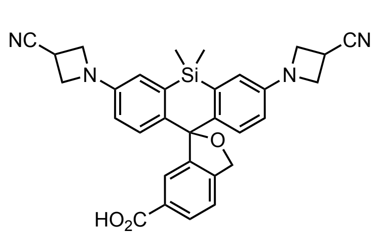


**6-Carboxy-JF$_{629}$b (S18):** The title compound (88%, pale blue solid) was prepared from 6-*tert*-butoxycarbonyl-JF$_{629}$b (**S10**) according to method S2. $^1$H NMR ($CD_3OD$, 400 MHz) δ 7.91 (dd, *J* = 7.9, 1.5 Hz, 1H), 7.47 (d, *J* = 1.4 Hz, 1H), 7.44 (d, *J* = 8.0 Hz, 1H), 7.14 (d, *J* = 8.7 Hz, 2H), 6.74 (d, *J* = 2.6 Hz, 2H), 6.46 (dd, *J* = 8.7, 2.7 Hz, 2H), 5.49

(s, 2H), 4.18 – 4.12 (m, 4H), 4.04 – 3.98 (m, 4H), 3.70 (tt, *J* = 8.3, 5.6 Hz, 2H), 0.65 (s, 3H), 0.49 (s, 3H); Analytical HPLC: $t_R$ = 12.4 min, >99% purity (10–95% MeCN/$H_2O$, linear gradient, with constant 0.1% v/v TFA additive; 20 min run; 1 mL/min flow; ESI; positive ion mode; detection at 650 nm); HRMS (ESI) calcd for $C_{31}H_{29}N_4O_3Si$ $[M+H]^+$ 533.2003, found 533.2007.

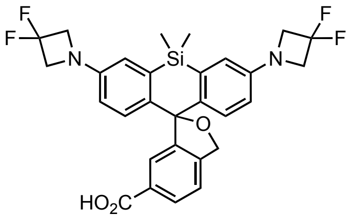


**6-Carboxy-JF$_{614}$b (S19):** The title compound (95%, blue solid) was prepared from 6-*tert*-butoxycarbonyl-JF$_{614}$b (**S11**) according to method S2. $^1$H NMR ($CDCl_3$, 400 MHz) δ 8.03 (dd, *J* = 8.0, 1.5 Hz, 1H), 7.75 (d, *J* = 1.6 Hz, 1H), 7.39 (d, *J* = 8.0 Hz, 1H), 7.07 (d, *J* = 8.6 Hz, 2H), 6.71 (d, *J* = 2.7 Hz, 2H), 6.40 (dd, *J* = 8.6, 2.7 Hz, 2H), 5.36 (s, 2H), 4.22 (t, $^3J_{HF}$ = 11.8 Hz, 8H), 0.66 (s, 3H), 0.54 (s, 3H); $^{19}$F NMR ($CDCl_3$, 376 MHz) δ -99.77 (p, $^3J_{FH}$ = 12.0 Hz); $^{13}$C NMR ($CDCl_3$, 101 MHz) δ 171.4 (C), 148.2 (t, $^4J_{CF}$ = 2.5 Hz, C), 147.0 (C), 145.1 (C), 140.6 (C), 135.5 (C), 129.8 (CH), 129.2 (C), 128.3 (CH), 126.3 (CH), 121.7 (CH), 116.8 (CH), 116.1 (t, $^1J_{CF}$ = 274.7 Hz, $CF_2$), 113.9 (CH), 92.4 (C), 72.9 ($CH_2$), 63.5 (t, $^2J_{CF}$ = 25.7 Hz, $CH_2$), 0.5 ($CH_3$), -0.8 ($CH_3$); Analytical HPLC: $t_R$ = 15.0 min, >99% purity (10–95% MeCN/$H_2O$, linear gradient, with constant 0.1% v/v TFA additive; 20 min run; 1 mL/min flow; ESI; positive ion mode; detection at 254 nm); HRMS (ESI) calcd for $C_{29}H_{27}F_4N_2O_3Si$ $[M+H]^+$ 555.1722, found 555.1734.

#### SYNTHESIS OF JFB–HALOTAG LIGANDS AND OTHER DERIVATIVES

**General method S3: Preparation of HaloTag ligands and maleimides.** The following procedure for $JF_{614}$b–HaloTag ligand (**$10_{HTL}$**) is representative. 6-Carboxy-$JF_{614}$b (**S19**; 50 mg, 90.2 μmol) and HATU (51.4 mg, 0.135 mmol, 1.5 eq) were combined in DMF (4 mL); 2-(2-((6-chlorohexyl)oxy)ethoxy)ethanamine hydrochloride (**S20**; 35.2 mg, 0.135 mmol, 1.5 eq) and DIEA (47.1 μL, 0.270 mmol, 3 eq) were added, and the reaction was stirred at room temperature for 4 h. It was subsequently diluted with saturated $NaHCO_3$ and extracted with EtOAc (2×). The combined organic extracts were washed with brine, dried over anhydrous $MgSO_4$, filtered, and concentrated *in vacuo*. Silica gel chromatography (10–100% EtOAc/toluene, linear gradient) afforded 55.8 mg (81%) of **$10_{HTL}$** as a white solid.

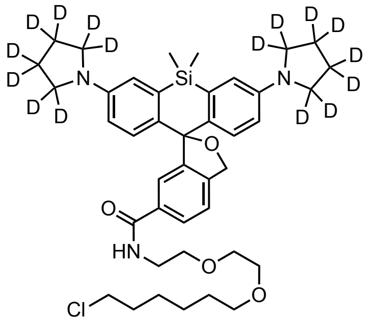


**$JFX_{650}$b–HaloTag ligand ($3_{HTL}$):** The title compound (63%, blue solid) was prepared from 6-carboxy-$JFX_{650}$b (**S12**) according to method S3. $^1$H NMR ($CDCl_3$, 400 MHz) δ 7.75 (dd, $J$ = 7.8, 1.6 Hz, 1H), 7.39 (d, $J$ = 1.6 Hz, 1H), 7.34 (d, $J$ = 7.9 Hz, 1H), 6.95 (d, $J$ = 8.8 Hz, 2H), 6.77 (d, $J$ = 2.8 Hz, 2H), 6.56 (t, $J$ = 4.5 Hz, 1H), 6.45 (dd, $J$ = 8.8, 2.8 Hz, 2H), 5.29 (s, 2H), 3.64 – 3.55 (m, 6H), 3.55 – 3.51 (m, 2H), 3.49 (t, $J$ = 6.7 Hz, 2H), 3.37 (t, $J$ = 6.6 Hz, 2H), 1.76 – 1.68 (m, 2H), 1.55 – 1.47 (m, 2H), 1.43 – 1.34 (m, 2H), 1.33 – 1.26 (m, 2H), 0.64 (s, 3H), 0.53 (s, 3H); Analytical HPLC: $t_R$ = 13.4 min, >99% purity (10–95% $MeCN/H_2O$, linear gradient, with constant 0.1% v/v TFA additive; 20 min run; 1 mL/min flow; ESI; positive ion mode; detection at 650 nm); HRMS (ESI) calcd for $C_{41}H_{39}D_{16}ClN_3O_4Si$ $[M+H]^+$ 732.4649, found 732.4638.

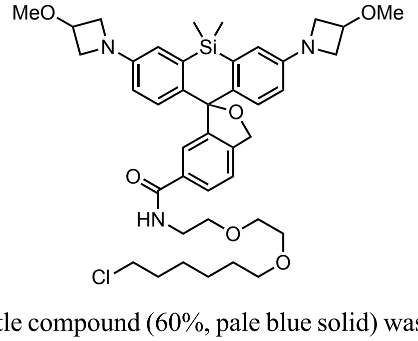


**$JF_{639}$b–HaloTag ligand ($4_{HTL}$):** The title compound (60%, pale blue solid) was prepared from 6-carboxy-$JF_{639}$b (**S13**) according to method S3. $^1$H NMR ($CDCl_3$, 400 MHz) δ 7.62 (dd, $J$ = 7.9, 1.5 Hz, 1H), 7.30 (d, $J$ = 1.5 Hz, 1H), 7.21 (d, $J$ = 7.9 Hz, 1H), 6.82 (d, $J$ = 8.6 Hz, 2H), 6.55 (d, $J$ = 2.6 Hz, 2H), 6.48 (t, $J$ = 4.4 Hz, 1H), 6.21 (dd, $J$ = 8.6, 2.7 Hz, 2H), 5.14 (s, 2H), 4.25 – 4.14 (m, 2H), 4.02 – 3.91 (m, 4H), 3.63 – 3.54 (m, 4H), 3.52 – 3.43 (m, 6H), 3.42 – 3.38

(m, 2H), 3.37 (t, $J$ = 6.6 Hz, 2H), 3.24 (t, $J$ = 6.6 Hz, 2H), 3.19 (s, 6H), 1.63 – 1.55 (m, 2H), 1.41 – 1.34 (m, 2H), 1.30 – 1.21 (m, 2H), 1.20 – 1.14 (m, 2H), 0.50 (s, 3H), 0.38 (s, 3H); Analytical HPLC: $t_R$ = 12.6 min, >99% purity (10–95% MeCN/$H_2O$, linear gradient, with constant 0.1% v/v TFA additive; 20 min run; 1 mL/min flow; ESI; positive ion mode; detection at 650 nm); HRMS (ESI) calcd for $C_{41}H_{55}ClN_3O_6Si$ $[M+H]^+$ 748.3543, found 748.3548.

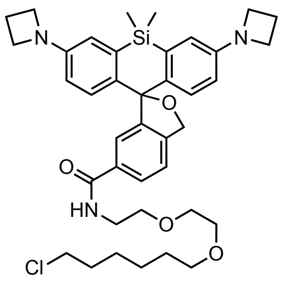

**JF$_{646}$b–HaloTag ligand (5$_{HTL}$):** The title compound (54%, blue solid) was prepared from 6-carboxy-JF$_{646}$b (**S14**) according to method S3. $^1H$ NMR ($CDCl_3$, 400 MHz) δ 7.75 (dd, $J$ = 7.8, 1.5 Hz, 1H), 7.43 (d, $J$ = 1.5 Hz, 1H), 7.35 (d, $J$ = 7.9 Hz, 1H), 6.94 (d, $J$ = 8.6 Hz, 2H), 6.66 (d, $J$ = 2.6 Hz, 2H), 6.62 (t, $J$ = 4.4 Hz, 1H), 6.32 (dd, $J$ = 8.6, 2.7 Hz, 2H), 5.28 (s, 2H), 3.88 (t, $J$ = 7.2 Hz, 8H), 3.66 – 3.55 (m, 6H), 3.56 – 3.52 (m, 2H), 3.50 (t, $J$ = 6.7 Hz, 2H), 3.38 (t, $J$ = 6.6 Hz, 2H), 2.35 (p, $J$ = 7.2 Hz, 4H), 1.77 – 1.69 (m, 2H), 1.55 – 1.48 (m, 2H), 1.43 – 1.35 (m, 2H), 1.33 – 1.27 (m, 2H), 0.63 (s, 3H), 0.52 (s, 3H); Analytical HPLC: $t_R$ = 12.8 min, >99% purity (10–95% MeCN/$H_2O$, linear gradient, with constant 0.1% v/v TFA additive; 20 min run; 1 mL/min flow; ESI; positive ion mode; detection at 650 nm); HRMS (ESI) calcd for $C_{39}H_{51}ClN_3O_4Si$ $[M+H]^+$ 688.3332, found 688.3330.

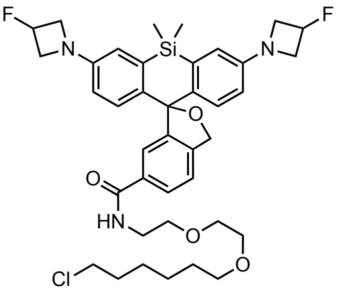

**JF$_{635}$b–HaloTag ligand (6$_{HTL}$):** The title compound (51%, pale blue solid) was prepared from 6-carboxy-JF$_{635}$b (**S15**) according to method S3. $^1H$ NMR ($CDCl_3$, 400 MHz) δ 7.66 (dd, $J$ = 7.9, 1.5 Hz, 1H), 7.38 (d, $J$ = 1.5 Hz, 1H), 7.28 (d, $J$ = 7.9 Hz, 1H), 6.93 (d, $J$ = 8.6 Hz, 2H), 6.62 (d, $J$ = 2.7 Hz, 2H), 6.55 (t, $J$ = 4.4 Hz, 1H), 6.29 (dd, $J$ = 8.6, 2.7 Hz, 2H), 5.33 (dtt, $^2J_{HF}$ = 57.1 Hz, $J$ = 5.9, 3.8 Hz, 2H), 5.23 (s, 2H), 4.19 – 4.05 (m, 4H), 3.96 – 3.82 (m, 4H), 3.58 – 3.50 (m, 6H), 3.49 – 3.45 (m, 2H), 3.43 (t, $J$ = 6.7 Hz, 2H), 3.32 (t, $J$ = 6.7 Hz, 2H), 1.70 – 1.61 (m, 2H), 1.48 – 1.41 (m, 2H), 1.36 – 1.28 (m, 2H), 1.27 – 1.20 (m, 2H), 0.58 (s, 3H), 0.46 (s, 3H); $^{19}F$ NMR ($CDCl_3$, 376 MHz) δ -180.43 (dtt, $J_{FH}$ = 57.2, 24.0, 18.0 Hz); Analytical HPLC: $t_R$ = 14.1 min, 98.1% purity (10–95% MeCN/$H_2O$, linear gradient, with constant 0.1% v/v TFA additive; 20 min run; 1 mL/min flow; ESI; positive ion mode; detection at 650 nm); HRMS (ESI) calcd for $C_{39}H_{49}ClF_2N_3O_4Si$ $[M+H]^+$ 724.3143, found 724.3141.

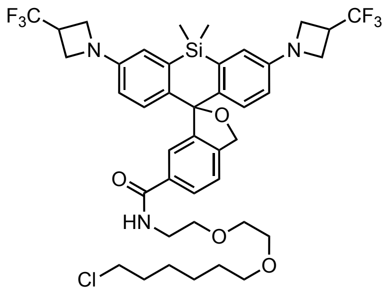

**JF$_{626}$b–HaloTag ligand (7$_{HTL}$):** The title compound (75%, white solid) was prepared from 6-carboxy-JF$_{626}$b (**S16**) according to method S3. $^1$H NMR (CDCl$_3$, 400 MHz) δ 7.73 (dd, *J* = 7.9, 1.6 Hz, 1H), 7.45 (d, *J* = 1.7 Hz, 1H), 7.35 (d, *J* = 7.9 Hz, 1H), 7.00 (d, *J* = 8.6 Hz, 2H), 6.67 (d, *J* = 2.7 Hz, 2H), 6.60 (t, *J* = 4.8 Hz, 1H), 6.34 (dd, *J* = 8.7, 2.7 Hz, 2H), 5.29 (s, 2H), 4.09 – 4.02 (m, 4H), 3.99 – 3.91 (m, 4H), 3.65 – 3.57 (m, 6H), 3.56 – 3.52 (m, 2H), 3.49 (t, *J* = 6.7 Hz, 2H), 3.44 – 3.32 (m, 2H), 3.38 (t, *J* = 6.6 Hz, 2H), 1.77 – 1.68 (m, 2H), 1.54 – 1.48 (m, 2H), 1.43 – 1.35 (m, 2H), 1.33 – 1.25 (m, 2H), 0.64 (s, 3H), 0.52 (s, 3H); $^{19}$F NMR (CDCl$_3$, 376 MHz) δ -73.30 (d, $^3J_{FH}$ = 8.6 Hz); Analytical HPLC: $t_R$ = 15.9 min, >99% purity (10–95% MeCN/$H_2O$, linear gradient, with constant 0.1% v/v TFA additive; 20 min run; 1 mL/min flow; ESI; positive ion mode; detection at 650 nm); HRMS (ESI) calcd for $C_{41}H_{48}ClF_6N_3O_4SiNa$ [M+Na]$^+$ 846.2899, found 846.2908.

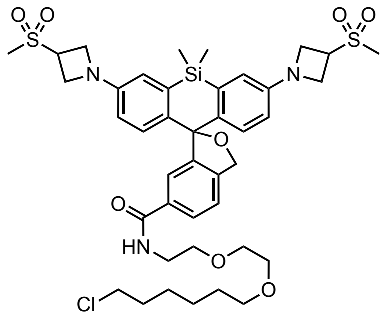

**JF$_{630}$b–HaloTag ligand (8$_{HTL}$):** The title compound (64%, pale blue solid) was prepared from 6-carboxy-JF$_{630}$b (**S17**) according to method S3. $^1$H NMR (CDCl$_3$, 400 MHz) δ 7.58 (dd, *J* = 7.9, 1.5 Hz, 1H), 7.31 (d, *J* = 1.6 Hz, 1H), 7.22 (d, *J* = 7.9 Hz, 1H), 6.92 (d, *J* = 8.6 Hz, 2H), 6.56 (d, *J* = 2.7 Hz, 2H), 6.50 (t, *J* = 4.3 Hz, 1H), 6.24 (dd, *J* = 8.6, 2.7 Hz, 2H), 5.19 (s, 2H), 4.17 – 4.01 (m, 8H), 3.97 – 3.88 (m, 2H), 3.51 – 3.43 (m, 6H), 3.43 – 3.39 (m, 2H), 3.37 (t, *J* = 6.7 Hz, 2H), 3.26 (t, *J* = 6.6 Hz, 2H), 2.86 (s, 6H), 1.63 – 1.55 (m, 2H), 1.41 – 1.35 (m, 2H), 1.30 – 1.23 (m, 2H), 1.21 – 1.15 (m, 2H), 0.52 (s, 3H), 0.39 (s, 3H); Analytical HPLC: $t_R$ = 12.7 min, >99% purity (10–95% MeCN/$H_2O$, linear gradient, with constant 0.1% v/v TFA additive; 20 min run; 1 mL/min flow; ESI; positive ion mode; detection at 650 nm); HRMS (ESI) calcd for $C_{41}H_{55}ClN_3O_8S_2SiNa$ [M+Na]$^+$ 866.2702, found 866.2715.

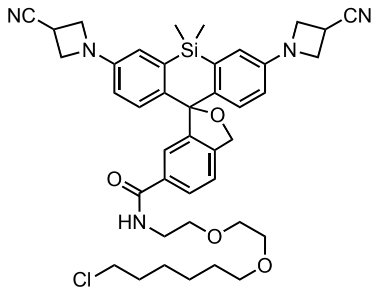


**$JF_{629}$b–HaloTag ligand ($9_{HTL}$):** The title compound (57%, pale blue solid) was prepared from 6-carboxy-$JF_{629}$b (**S18**) according to method S3. $^1$H NMR ($CDCl_3$, 400 MHz) δ 7.70 (dd, $J$ = 7.9, 1.5 Hz, 1H), 7.46 (d, $J$ = 1.5 Hz, 1H), 7.36 (d, $J$ = 7.9 Hz, 1H), 7.07 (d, $J$ = 8.7 Hz, 2H), 6.71 (d, $J$ = 2.6 Hz, 2H), 6.56 (t, $J$ = 4.5 Hz, 1H), 6.40 (dd, $J$ = 8.7, 2.7 Hz, 2H), 5.33 (s, 2H), 4.23 (ddd, $J$ = 8.4, 7.1, 2.6 Hz, 4H), 4.10 (td, $J$ = 6.8, 2.7 Hz, 4H), 3.65 – 3.53 (m, 10H), 3.50 (t, $J$ = 6.7 Hz, 2H), 3.40 (t, $J$ = 6.7 Hz, 2H), 1.77 – 1.68 (m, 2H), 1.57 – 1.49 (m, 2H), 1.44 – 1.35 (m, 2H), 1.35 – 1.25 (m, 2H), 0.66 (s, 3H), 0.53 (s, 3H); Analytical HPLC: $t_R$ = 14.2 min, >99% purity (10–95% MeCN/$H_2O$, linear gradient, with constant 0.1% v/v TFA additive; 20 min run; 1 mL/min flow; ESI; positive ion mode; detection at 650 nm); HRMS (ESI) calcd for $C_{41}H_{49}ClN_5O_4Si$ $[M+H]^+$ 738.3237, found 738.3233.

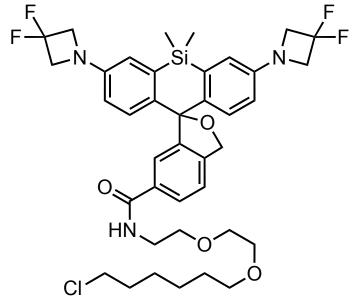


**$JF_{614}$b–HaloTag ligand ($10_{HTL}$):** The title compound (81%, white solid) was prepared from 6-carboxy-$JF_{614}$b (**S19**) according to method S3. $^1$H NMR ($CDCl_3$, 400 MHz) δ 7.72 (dd, $J$ = 7.9, 1.6 Hz, 1H), 7.46 (d, $J$ = 1.5 Hz, 1H), 7.36 (d, $J$ = 7.9 Hz, 1H), 7.05 (d, $J$ = 8.6 Hz, 2H), 6.71 (d, $J$ = 2.7 Hz, 2H), 6.61 (t, $J$ = 4.8 Hz, 1H), 6.39 (dd, $J$ = 8.6, 2.7 Hz, 2H), 5.32 (s, 2H), 4.22 (t, $^3J_{HF}$ = 11.8 Hz, 8H), 3.65 – 3.57 (m, 6H), 3.57 – 3.52 (m, 2H), 3.50 (t, $J$ = 6.7 Hz, 2H), 3.39 (t, $J$ = 6.6 Hz, 2H), 1.77 – 1.68 (m, 2H), 1.56 – 1.48 (m, 2H), 1.44 – 1.35 (m, 2H), 1.34 – 1.27 (m, 2H), 0.66 (s, 3H), 0.54 (s, 3H); $^{19}$F NMR ($CDCl_3$, 376 MHz) δ -99.79 (p, $^3J_{FH}$ = 11.8 Hz); Analytical HPLC: $t_R$ = 15.0 min, >99% purity (30–95% MeCN/$H_2O$, linear gradient, with constant 0.1% v/v TFA additive; 20 min run; 1 mL/min flow; ESI; positive ion mode; detection at 254 nm); HRMS (ESI) calcd for $C_{39}H_{46}ClF_4N_3O_4SiNa$ $[M+Na]^+$ 782.2774, found 782.2784.

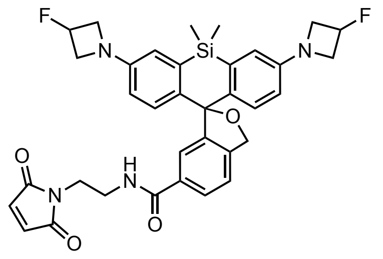

**JF$_{635}$b–maleimide (6$_{MAL}$):** The title compound (83%, yellow solid) was prepared from 6-carboxy-JF$_{635}$b (**S15**) and *N*-(2-aminoethyl)maleimide trifluoroacetate salt according to method S3. $^{1}$H NMR ($CDCl_3$, 400 MHz) δ 7.66 (dd, *J* = 7.9, 1.6 Hz, 1H), 7.47 (dd, *J* = 1.6, 0.6 Hz, 1H), 7.35 (dd, *J* = 7.9, 0.5 Hz, 1H), 6.98 (d, *J* = 8.6 Hz, 2H), 6.69 (d, *J* = 2.6 Hz, 2H), 6.68 (s, 2H), 6.55 (t, *J* = 5.4 Hz, 1H), 6.35 (dd, *J* = 8.6, 2.6 Hz, 2H), 5.40 (dtt, $^{2}J_{HF}$ = 57.1 Hz, *J* = 5.9, 3.7 Hz, 2H), 5.27 (s, 2H), 4.24 – 4.13 (m, 4H), 4.01 – 3.89 (m, 4H), 3.80 – 3.74 (m, 2H), 3.64 – 3.57 (m, 2H), 0.64 (s, 3H), 0.53 (s, 3H); $^{19}$F NMR ($CDCl_3$, 376 MHz) δ -180.43 (dtt, $J_{FH}$ = 57.0, 24.2, 18.1 Hz); $^{13}$C NMR ($CDCl_3$, 101 MHz) δ 171.1 (C), 167.5 (C), 149.5 (d, $^{4}J_{CF}$ = 1.1 Hz, C), 146.9 (C), 143.2 (C), 139.7 (C), 135.6 (C), 134.4 (CH), 133.9 (C), 128.3 (CH), 126.2 (CH), 123.6 (CH), 121.6 (CH), 116.3 (CH), 113.3 (CH), 92.6 (C), 82.9 (d, $^{1}J_{CF}$ = 204.5 Hz, CHF), 72.5 ($CH_2$), 59.7 (d, $^{2}J_{CF}$ = 23.5 Hz, $CH_2$), 39.8 ($CH_2$), 37.5 ($CH_2$), 0.6 ($CH_3$), -1.0 ($CH_3$); Analytical HPLC: $t_R$ = 11.9 min, >99% purity (10–95% MeCN/$H_2O$, linear gradient, with constant 0.1% v/v TFA additive; 20 min run; 1 mL/min flow; ESI; positive ion mode; detection at 650 nm); HRMS (ESI) calcd for $C_{35}H_{35}F_2N_4O_4Si$ $[M+H]^+$ 641.2390, found 641.2385.

**General method S4: NHS ester formation.** The following procedure for JF$_{630}$b-NHS (**8$_{NHS}$**) is representative. 6-Carboxy-JF$_{630}$b (**S17**; 30 mg, 47.0 μmol) was combined with DSC (28.9 mg, 0.113 mmol, 2.4 eq) in DMF (1.5 mL). After adding $Et_3N$ (39.3 μL, 0.282 mmol, 6 eq) and DMAP (0.6 mg, 4.7 μmol, 0.1 eq), the reaction was stirred at room temperature for 1 h. It was subsequently concentrated *in vacuo* and purified by reverse phase HPLC (20–60% MeCN/$H_2O$, linear gradient, with constant 0.1% TFA additive). The pooled product fractions were partially concentrated to remove MeCN, neutralized with 1–2 mL saturated $NaHCO_3$, and extracted with $CH_2Cl_2$ (2×). The combined organic layers were washed with brine, dried over anhydrous $MgSO_4$, filtered, and evaporated to yield 17.8 mg (52%) of **8$_{NHS}$** as a pale blue solid.

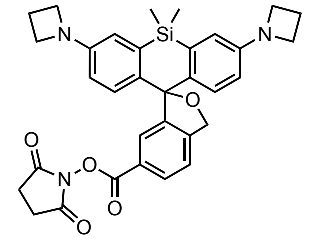

**JF$_{646}$b–NHS (5$_{NHS}$):** The title compound (32%, yellow-green solid) was prepared from 6-carboxy-JF$_{646}$b (**S14**) according to method S4. $^{1}$H NMR ($CDCl_3$, 400 MHz) δ 7.88 (dd, *J* = 7.9, 1.5 Hz, 1H), 7.59 (d, *J* = 1.5 Hz, 1H), 7.24 (d, *J* = 8.0 Hz, 1H), 6.78 (d, *J* = 8.6 Hz, 2H), 6.47 (d, *J* = 2.6 Hz, 2H), 6.17 (dd, *J* = 8.6, 2.7 Hz, 2H), 5.17 (s, 2H), 3.71 (t, *J* = 7.2 Hz, 8H), 2.74 – 2.63 (m, 4H), 2.16 (p, *J* = 7.3 Hz, 4H), 0.43 (s, 3H), 0.32 (s, 3H); $^{13}$C NMR ($CDCl_3$,

101 MHz) δ 169.4 (C), 161.8 (C), 150.7 (C), 148.1 (C), 146.8 (C), 138.5 (C), 135.0 (C), 129.8 (CH), 128.1 (CH), 126.9 (CH), 124.7 (C), 121.9 (CH), 115.8 (CH), 112.9 (CH), 92.8 (C), 72.7 ($CH_2$), 52.6 ($CH_2$), 25.8 ($CH_2$), 17.1 ($CH_2$), 0.5 ($CH_3$), -0.8 ($CH_3$); Analytical HPLC: $t_R$ = 11.6 min, 96.5% purity (10–95% MeCN/$H_2O$, linear gradient, with constant 0.1% v/v TFA additive; 20 min run; 1 mL/min flow; ESI; positive ion mode; detection at 650 nm); HRMS (ESI) calcd for $C_{33}H_{34}N_3O_5Si$ $[M+H]^+$ 580.2262, found 580.2260.

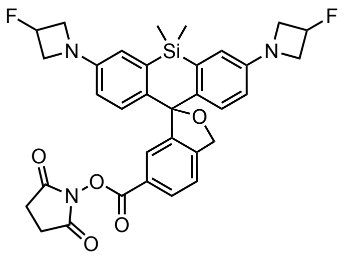

**$JF_{635}$b–NHS ($6_{NHS}$):** The title compound (56%, off-white solid) was prepared from 6-carboxy-$JF_{635}$b (**S15**) according to method S4. $^1H$ NMR ($CDCl_3$, 400 MHz) δ 8.10 (dd, $J$ = 8.0, 1.6 Hz, 1H), 7.79 (d, $J$ = 1.5 Hz, 1H), 7.46 (d, $J$ = 8.0 Hz, 1H), 7.04 (d, $J$ = 8.7 Hz, 2H), 6.71 (d, $J$ = 2.7 Hz, 2H), 6.42 (dd, $J$ = 8.7, 2.7 Hz, 2H), 5.43 (dtt, $^2J_{HF}$ = 57.1 Hz, $J$ = 6.0, 3.7 Hz, 1H), 5.39 (s, 2H), 5.38 – 5.33 (m, 1H), 4.28 – 4.16 (m, 4H), 4.06 – 3.94 (m, 4H), 2.90 (s, 4H), 0.65 (s, 3H), 0.55 (s, 3H); $^{19}F$ NMR ($CDCl_3$, 376 MHz) δ -180.53 (dtt, $J_{FH}$ = 57.4, 24.1, 18.1 Hz); $^{13}C$ NMR ($CDCl_3$, 101 MHz) δ 169.3 (C), 161.7 (C), 149.6 (d, $^4J_{CF}$ = 1.2 Hz, C), 147.9 (C), 146.5 (C), 139.4 (C), 135.2 (C), 129.9 (CH), 128.2 (CH), 126.8 (CH), 124.8 (C), 122.1 (CH), 116.3 (CH), 113.5 (CH), 92.6 (C), 83.0 (d, $^1J_{CF}$ = 204.5 Hz, CHF), 72.9 ($CH_2$), 59.7 (d, $^2J_{CF}$ = 23.6 Hz, $CH_2$), 25.8 ($CH_2$), 0.5 ($CH_3$), -0.8 ($CH_3$); Analytical HPLC: $t_R$ = 12.6 min, >99% purity (10–95% MeCN/$H_2O$, linear gradient, with constant 0.1% v/v TFA additive; 20 min run; 1 mL/min flow; ESI; positive ion mode; detection at 650 nm); HRMS (ESI) calcd for $C_{33}H_{32}F_2N_3O_5Si$ $[M+H]^+$ 616.2074, found 616.2073.

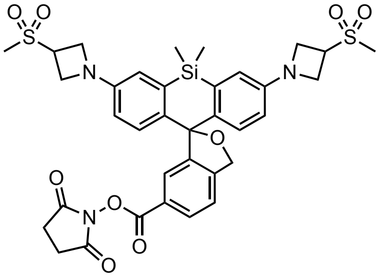

**$JF_{630}$b–NHS ($8_{NHS}$):** The title compound (52%, pale blue solid) was prepared from 6-carboxy-$JF_{630}$b (**S17**) according to method S4. $^1H$ NMR ($CDCl_3$, 400 MHz) δ 8.07 (dd, $J$ = 8.0, 1.5 Hz, 1H), 7.74 (dd, $J$ = 1.6, 0.7 Hz, 1H), 7.44 (dd, $J$ = 8.0, 0.6 Hz, 1H), 7.06 (d, $J$ = 8.6 Hz, 2H), 6.69 (d, $J$ = 2.6 Hz, 2H), 6.41 (dd, $J$ = 8.7, 2.7 Hz, 2H), 5.39 (s, 2H), 4.27 – 4.17 (m, 8H), 4.09 – 4.01 (m, 2H), 3.00 (s, 6H), 2.87 (s, 4H), 0.63 (s, 3H), 0.51 (s, 3H); $^{13}C$ NMR ($CDCl_3$, 101 MHz) δ 169.4 (C), 161.7 (C), 148.7 (C), 147.7 (C), 146.2 (C), 140.3 (C), 135.1 (C), 130.0 (CH), 128.2 (CH), 126.6 (CH), 124.8 (C), 122.2 (CH), 116.2 (CH), 113.4 (CH), 92.5 (C), 73.0 ($CH_2$), 52.7 ($CH_2$), 52.0 (CH), 37.9 ($CH_3$), 25.8 ($CH_2$), 0.4 ($CH_3$), -0.7 ($CH_3$); Analytical HPLC: $t_R$ = 12.3 min, 98.3% purity (10–95% MeCN/$H_2O$, linear gradient, with constant 0.1% v/v TFA additive; 20 min run; 1 mL/min flow; ESI; positive ion mode; detection at 650 nm); HRMS (ESI) calcd for $C_{35}H_{38}N_3O_9S_2Si$ $[M+H]^+$ 736.1813, found 736.1814.

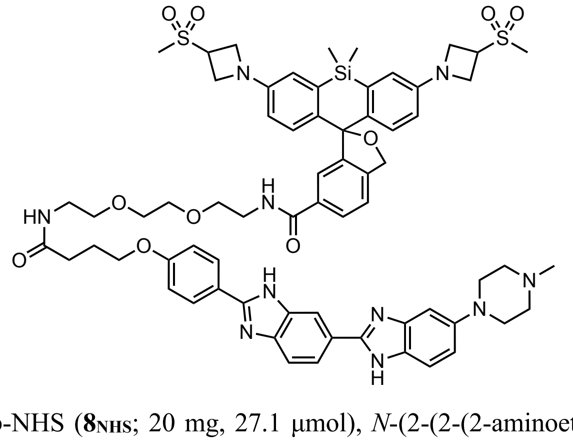


**JF$_{630}$b–Hoechst (8$_{HST}$):** JF$_{630}$b-NHS (**8$_{NHS}$**; 20 mg, 27.1 µmol), *N*-(2-(2-(2-aminoethoxy)ethoxy)ethyl)-4-(4-(5-(4-methylpiperazin-1-yl)-1*H*,3'*H*-[2,5'-bibenzo[*d*]imidazol]-2'-yl)phenoxy)butanamide[1] (4·TFA salt; 35.8 mg, 32.6 µmol, 1.2 eq), and DIEA (47.3 µL, 0.272 mmol, 10 eq) were combined in DMF (1 mL). The reaction was stirred for 4 h at room temperature, then concentrated *in vacuo*. The crude residue was purified by reverse phase HPLC (10–75% MeCN/$H_2O$, linear gradient, with constant 0.1% TFA additive). The pooled product fractions were partially concentrated to remove MeCN, neutralized with saturated $NaHCO_3$, and extracted with 10% MeOH/$CH_2Cl_2$ (2×). The combined organic layers were dried over anhydrous $MgSO_4$, filtered, and evaporated to yield 5.0 mg (15%) of **8$_{HST}$** as a yellow solid. $^1H$ NMR ($CD_3OD$, 400 MHz) δ 8.26 (s, 1H), 8.04 (d, *J* = 8.9 Hz, 2H), 7.96 (d, *J* = 8.4 Hz, 1H), 7.70 (dd, *J* = 8.0, 1.6 Hz, 1H), 769 (bs, 1H), 7.52 (d, *J* = 8.8 Hz, 1H), 7.39 (d, *J* = 8.0 Hz, 1H), 7.33 (d, *J* = 1.5 Hz, 1H), 7.19 – 7.13 (m, 1H), 7.09 – 7.05 (m, 1H), 7.06 (d, *J* = 8.9 Hz, 2H), 7.03 (d, *J* = 8.7 Hz, 2H), 6.72 (d, *J* = 2.6 Hz, 2H), 6.41 (dd, *J* = 8.7, 2.7 Hz, 2H), 5.37 (s, 2H), 4.32 – 4.24 (m, 2H), 4.19 – 4.08 (m, 8H), 4.01 (t, *J* = 6.1 Hz, 2H), 3.58 – 3.49 (m, 6H), 3.48 – 3.41 (m, 4H), 3.29 – 3.23 (m, 6H), 2.98 (s, 6H), 2.81 – 2.72 (m, 4H), 2.44 (s, 3H), 2.34 (t, *J* = 7.3 Hz, 2H), 2.04 (p, *J* = 6.7 Hz, 2H), 0.61 (s, 3H), 0.46 (s, 3H); Analytical HPLC: $t_R$ = 7.9 min, >99% purity (10–95% MeCN/$H_2O$, linear gradient, with constant 0.1% v/v TFA additive; 20 min run; 1 mL/min flow; ESI; positive ion mode; detection at 650 nm); HRMS (ESI) calcd for $C_{66}H_{77}N_{10}O_{10}S_2Si$ $[M+H]^+$ 1261.5029, found 1261.5026.

[1] Nakamura, A.; Takigawa, K.; Kurishita, Y.; Kuwata, K.; Ishida, M.; Shimoda, Y.; Hamachi, I.; Tsukiji, S. *Chem. Commun.* **2014**, *50*, 6149–6152.
